# A Metabolic Labeling Strategy for Tracking Protein Synthesis in Complex Biological Systems

**DOI:** 10.64898/2026.08.30.747940

**Authors:** Yong Jia Bu, Samuel P. Nyandwi, Flavia de Lima Alves, Rasangi Tennakoon, Tabea V. Stamm, David J. Schneider, Alexander Eddenden, Timothy Wing Yin Ma, Yoo-jeong Chun, Jolie Marie Miller, Hui Peng, Aaron R. Wheeler, Scott A. Yuzwa, Mark Nitz, Haissi Cui

## Abstract

Protein synthesis supports most biological processes. In the brain in particular, protein synthesis plays a critical role in physiological and pathological states. Here, we describe <u>Te</u>llurophene-<u>A</u>lkyne <u>C</u>ycloaddition-mediated <u>A</u>mino acid <u>T</u>agging (TeACAT), a versatile strategy for fast, facile, and flexible tagging of newly synthesized proteins in mice.

TeACAT is based on metabolic incorporation of the non-canonical amino acid TePhe into proteins by the endogenous protein synthesis machinery. Due to their high similarity, TePhe can efficiently replace canonical Phe without dietary or genetic manipulation. The subsequent bio-orthogonal reaction of TePhe with either fluorescent dyes or affinity handles enables both visualization and affinity enrichment of proteins synthesized during TePhe exposure. TeACAT is compatible with immunofluorescence for cell-type specific visualization of protein synthesis with subcellular resolution and can be used in conjunction with routine proteomics to identify and quantify newly synthesized proteins. Robust incorporation into the mouse proteome was observed on the scale of hours to days, allowing the interrogation of various biological processes.

In summary, TeACAT enables the visualization and quantification of protein synthesis with minimal perturbation for biological discoveries.

## Introduction

mRNA translation determines cellular phenotype through the production of functional protein ensembles. In turn, precise spatial and temporal coordination of protein synthesis between cells allows complex tissues to form and to carry out specialized functions. A prime example is the brain, where specialized translation in differentiated cells^1–4^ orchestrate the formation of elaborate three-dimensional networks^4–6^ and ultimately enable behaviour and learning.^7^ Consequently, aberrant protein synthesis is highly detrimental and perturbations can cause neurodevelopmental^8–11^ and neurodegenerative diseases.^11–13^ Given the central role of protein synthesis in all living systems, tools for its characterization are key for enabling new discoveries.

An ideal methodology for tracking global protein synthesis would minimally perturb the underlying biology, be readily applied across a range of model systems at varying timescales, and employ simple workflows using readily accessible reagents and equipment. The experimental readouts should preserve the spatial context that is key to revealing changes in cellular protein synthesis and relationships within complex tissues. Furthermore, the method should provide the ability to enrich newly synthesized proteins to facilitate their identification.

Existing methodologies for tracking protein synthesis in the mammalian brain address some of the above criteria, but are limited with regards to versatility and feasibilty.^14–18^ Metabolic incorporation of non-canonical amino acids (ncaas), such as the L-Azidohomoalanine (AHA)-based BONCAT method^19,20^, present the most powerful and generalizable strategy to provide both spatial and sequence-level readouts of new protein synthesis. However, challenges remain due to low incorporation efficiency under physiological conditions and need for genetically engineered animal models.

L-tellurienylalanine (TePhe)^21^, as a tellurophene-bearing analogue of L-phenylalanine (Phe), offers a solution. The 5-membered tellurophene ring is an effective phenyl bio-isostere whose inclusion into model proteins has minimal impacts on folding and stability.^22–25^ TePhe is readily taken up via the endogenous translation machinery^21^ and has been explored as a protein synthesis probe in imaging mass cytometry (IMC). However, IMC is limited in spatial resolution, imaging speed^26,27^, and requires to specialized instrumentation. To broaden the applicability of TePhe, we recently established a novel tellurophene-based bioconjugation reaction (<u>o</u>xidation-controlled, <u>s</u>train-promoted tellurophene <u>a</u>lkyne <u>c</u>ycloaddition — OSTAC) to interface TePhe incorporation with fluorescence- and affinity-tagging.^28^

Here, we leverage OSTAC chemistry for fluorescent visualization of protein synthesis in murine tissues, as well as for the enrichment of tagged proteins for mass spectrometry-based proteomics. Our approach provides the first demonstration of facile tagging and functionalization of newly synthesized proteins from the mammalian brain, accomplished entirely through endogenous translation machinery and free from dietary or genetic manipulation. We term this combination of TePhe metabolic incorporation and subsequent OSTAC bioconjugation TeACAT: <u>Te</u>llurophene-<u>A</u>lkyne <u>C</u>ycloaddition-mediated <u>A</u>mino acid <u>T</u>agging. TeACAT is fast, facile, and versatile, enabling the study of protein synthesis under minimally perturbing conditions for a diverse range of biological contexts.

## Results

### TePhe efficiently and robustly tags newly synthesized proteins in murine tissues

To explore the potential of TeACAT for fluorescent detection of newly synthesized proteins in murine tissues (Fig. 1a), we delivered TePhe to mice through intravenous (IV) injection, at 60 mg/kg TePhe formulated in 40% Captisol in PBS, as previously described for IMC.^21^ TePhe and vehicle-injected animals were sacrificed after 24 hr. OSTAC chemistry (Fig. 1b) was performed on tissues expected to have high rates of protein turnover, namely the liver and small intestine, where the highest TePhe incorporation was expected. A clear increase in fluorescence intensity could be observed in formalin-fixed cryosections from TePhe-treated mice over vehicle controls (Fig. S1; detailed workflow in Fig. S3). This suggested that OSTAC chemistry protocols, initially established in cell culture, could be transferred to mouse tissues.

**Figure 1.**
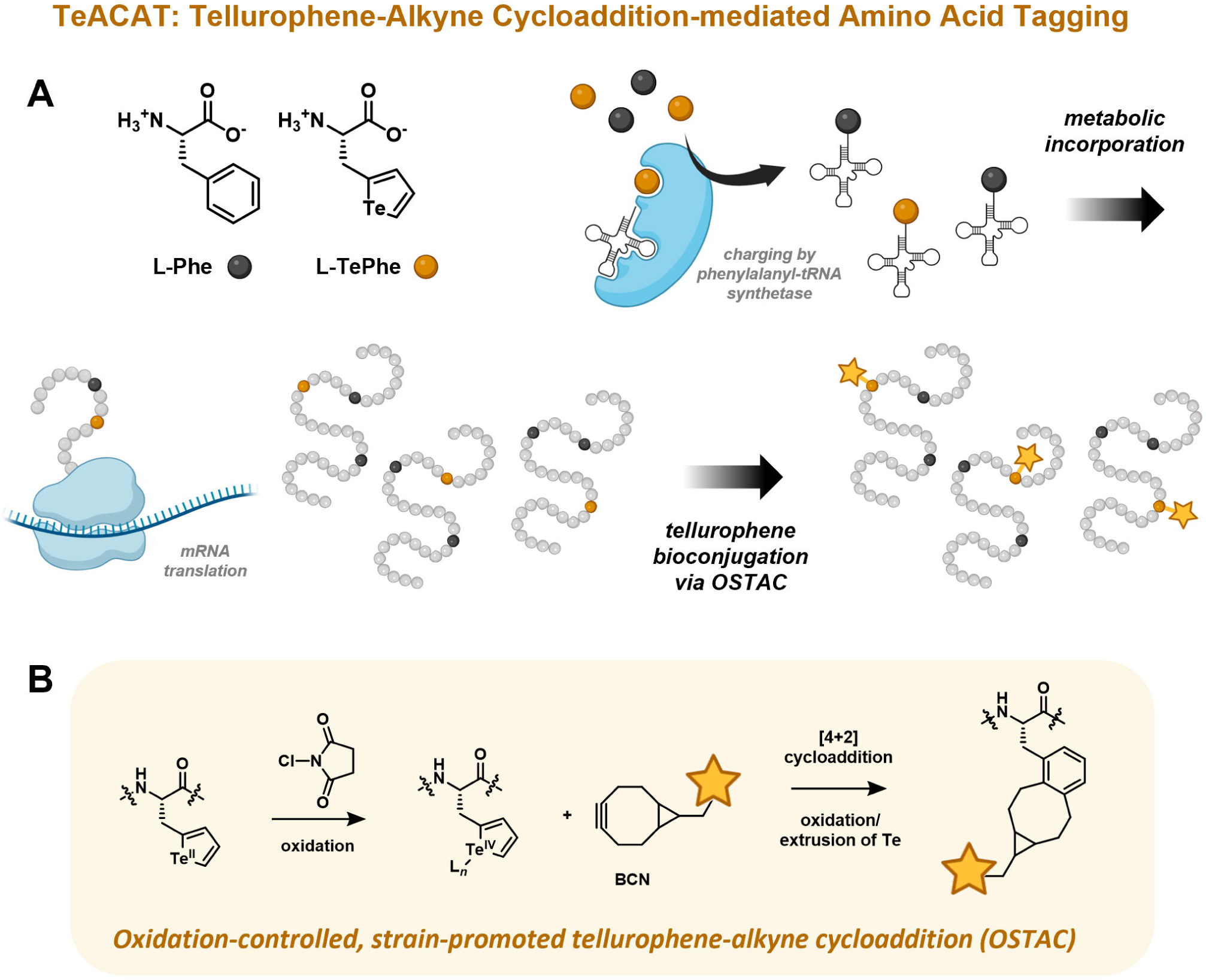
Overview of TeACAT (Tellurophene-alkyne cycloaddition-mediated amino acid tagging). A) Structures of L-Phe and L-TePhe. TePhe closely mimics Phe and is incorporated into newly synthesized proteins by the endogenous translation machinery. Incorporated TePhe can then be detected using the tellurophene-targeting bioconjugation shown in B). B) Oxidation-controlled, strain-promoted tellurophene alkyne cycloaddition (OSTAC) reaction sequence. The tellurophene side chain of TePhe undergoes a 2-electron oxidation reaction on the Te centre, which activates the ring for [4+2] cycloaddition with bicyclo[6.1.0]nonyne (BCN). Rapid extrusion of Te and re-aromatization of the system results in a highly stable benzocyclooctane adduct.

Encouraged by these results, we turned our attention to the central nervous system, which is challenging due to lower rates of protein synthesis^29–31^ and restricted import of labeling reagents through the blood-brain barrier.^32–34^ We explored the relationship between TePhe dose and TePhe incorporation by fluorescent labeling of lysates (using the segment of spinal cord directly below the brainstem) and quantification after SDS-PAGE (Fig. 2a; detailed workflow in Fig. S2a, gels shown in Fig. S4). For these and all subsequent experiments, we used intraperitoneal (IP) TePhe injection as it was more amenable to repeat dosing (dosing regime overview in Fig. 2b).

**Figure 2.**
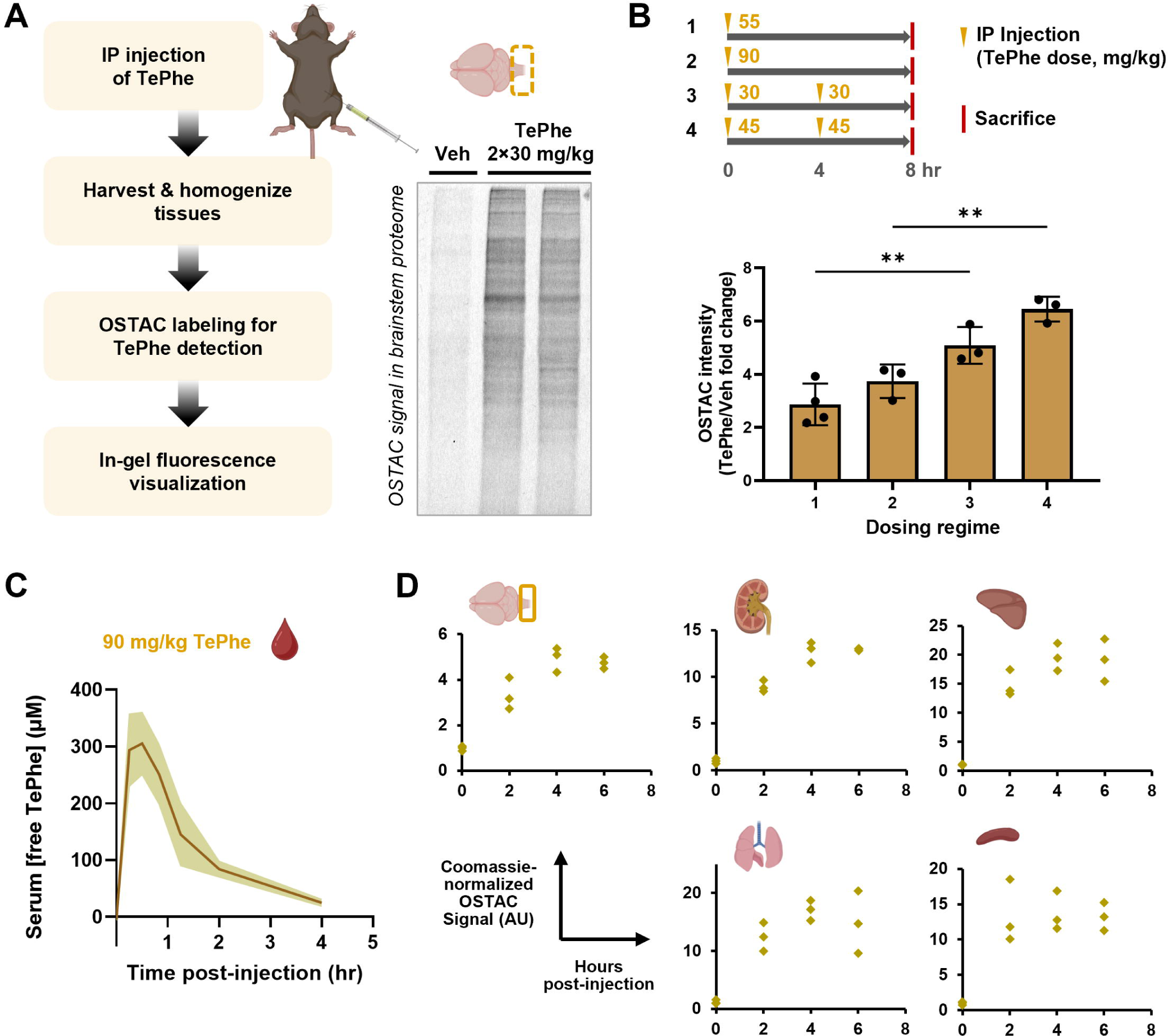
TePhe dosing regimes and incorporation characteristics. A) Overview of workflow to quantify TePhe incorporation in tissues, with example SDS-PAGE showing TAMRA fluorescence in OSTAC-labeled brainstem homogenates. B) TePhe dosing regimes and resultant brainstem TePhe content. Each datapoint represents one mouse (*n* = 4, 3, 3, 3 for conditions 1-4 respectively; one-way ANOVA, ** *p* <0.01). C) Free TePhe concentration in mouse serum after single IP injection (90 mg/kg), as determined by OSTAC-based fluorescent TLC assay. *n* = 3 per timepoint, 2 technical replicates each. Shaded region represents = standard deviation between all replicates. D) Time course of TePhe incorporation in major organs, determined by in-gel fluorescence of OSTAC-labeled tissue homogenates. Each datapoint represents one animal. *n* = 3 for all timepoints.

A single TePhe dose at 55 mg/kg yielded clearly discernible signal in spinal cord lysates over background; the signal did not increase significantly at 90 mg/kg (Fig. 2b, regimes 1-2). As amino acids are quickly cleared from the blood stream^35,36^, we hypothesized that repeated dosing might further increase incorporation compared to an increased bolus injection. Indeed, higher signal was obtained by administering two smaller doses of TePhe 4 hr apart (Fig. 2b, regimes 3-4): TePhe contents in mice receiving 2 x 30 mg/kg and 2 x 45 mg/kg were on average 1.7- to 1.8-fold higher than those receiving single doses of 55 mg/kg and 90 mg/kg, respectively.

To establish pharmacokinetics of TePhe distribution, we determined the concentration of free TePhe in mouse serum post-IP injection (90 mg/kg TePhe) using an OSTAC-based fluorescent TLC assay (Fig. 2c, Fig. S5a). Serum TePhe concentration peaked at ∼300 µM between 15 to 50 min post-IP injection, then gradually fell to ∼30 µM by 4 hr, leading to an estimated half-life of TePhe in serum of 1.1 hr (Fig. S5b). This clearance profile explains the higher TePhe incorporation upon repeat dosing at 4 hr intervals compared to a single bolus injection. Urine analysis by OSTAC-based fluorescent TLC suggested TePhe processing through Phe metabolic pathways and excretion of tellurophene-bearing analogues of Phe metabolites^37–40^ (Fig. S6-7). This suggests that TePhe can compete with systemic Phe levels. *In vitro* enzymatic assays using purified human mitochondrial phenylalanyl-tRNA synthetase (FARS2) confirmed comparably efficient activation of Phe and TePhe by FARS2 (Fig. S8).

To explore whether TePhe could be used to track protein synthesis over shorter periods, we assessed TePhe content in different tissues over time (Fig. 2d, Fig. S9a-f). Across the spinal cord, kidney, liver, lungs, and spleen, a common trend emerged, with half or more of the maximum TePhe incorporation occurring within the first 2 hr. This was followed by slower incorporation and a plateau after 4 hr, again in agreement with the clearance profile of free TePhe shown in Fig. 2c. Incorporation of TePhe into serum proteins was also clearly detectable by 2 hr (Fig. S9g), which presumably occurred through TePhe uptake in the liver, followed by liver protein synthesis, and secretion back into the blood stream, all within a short time frame.

We also probed the lifetime of TePhe-labeled proteins in the brain by increasing the spacing between sequential doses (3 x 30 mg/kg over 24 or 72 hr). Total TePhe levels in the brain proteome were comparable to 2 x 45 mg/kg injected within 8 hr (Fig. S10), suggesting that incorporated TePhe persists over days. Together, these results demonstrate rapid and systemic incorporation of TePhe across murine tissues and highlight its usefulness as a probe for protein synthesis over a range of timescales.

### TePhe is a minimally perturbing probe for brain protein synthesis

To ensure that TePhe is minimally perturbing, we performed detailed molecular and metabolic characterization. We observed minor discolouration of the spleen, gall bladder, and most consistently, kidneys of TePhe-treated animals, suggesting TePhe breakdown by Phe-catabolizing enzymes, possibly to elemental tellurium. We assessed kidney integrity and activity through immunochemistry and the biomarker creatine, both of which did not reveal obvious changes in TePhe-treated mice (Fig. S11-12). A reduction in blood urea nitrogen to the lower percentile and an elevation of aspartate aminotransferase (AST) levels was observed^41,42^, both of which could indicate liver damage, while alanine aminotransferase (ALT) levels remained comparable (Fig. S12). The magnitude of AST change matches mild hepatotoxicity.^43–46^ An acute high dose of TePhe thus mildly affected the liver while kidney function remained unchanged.

To exclude potential subtle perturbations in the brain upon TePhe exposure, we performed in-depth characterization of the brain’s response to TePhe incorporation. We compared the proteomes and transcriptomes of 2 mm coronal brain slices from vehicle and TePhe-treated mice (Fig. S13a, S14a). We chose the 2x 45 mg/kg TePhe dosing regime, which led to the highest level of incorporation and was therefore most likely to induce aberrant changes. Proteomics analysis revealed highly similar bulk proteomes with very few proteins of significantly altered abundance (14 down-regulated and 17 up-regulated at ≥1.5-fold change and *p-*value of <0.05, Student’s t-test, Fig. S13b-c). Overrepresentation testing did not indicate the induction of any stress-related pathways by TePhe (Fig. S13d). These results indicate that TePhe incorporation led to minimal perturbation of the brain proteome.

A stress response could alter gene expression before manifesting detectable proteome changes. RNA-seq showed minimal differential gene expression between vehicle and TePhe-treated groups, and no separation between groups was found in principle component analysis (Fig. S14b-c). Only one gene exceeded both abundance change and statistical significance cutoffs (*p*adj <0.05; fold change ≥2): Cdkn1a, encoding the cyclin-dependent kinase p21, which is up-regulated in response to damaged DNA and inflammation.^47,48^ We also combined all 114 genes meeting the *p*adj <0.05 significance cutoff, regardless of fold-change, and tested for GO term enrichment. Few were identified (Fig. S14d) such as loose associations with response to a stimulus and changing nutrient conditions, as expected. We specifically interrogated targets of ATF4 (a high-level regulator of the integrated stress response)^49^, myelination (elemental tellurium can cause dose-dependent demyelination)^50^, and Phe-associated pathways, but did not observe differential expression changes (Fig. S14e-g). The only observed clustering reflecting treatment groups were in Phe import-associated genes, which were upregulated in response to TePhe injections (Fig. S14g). Altogether, these results validated that working concentrations of TePhe were minimally perturbing to brain physiology.

### TePhe visualizes protein synthesis in brain sections with subcellular resolution

Following the extensive validation of TePhe as a minimally perturbing metabolic probe, we explored the use of OSTAC chemistry to visualize protein synthesis. Our existing OSTAC labeling protocol was adapted for 4 µm formalin-fixed, paraffin-embedded (FFPE) brain sections (Fig. S3), leading to a flexible protocol for visualization of newly synthesized proteins with TeACAT. After deparaffinization and rehydration, the optimized reaction protocol consisted of three simple steps, which can be completed in 1 hr: 1) alkylation of free thiols with iodoacetamide (IAA) for 20 min; 2) sample oxidation with *N-*chlorosuccinimide (NCS) and blocking of remaining off-target reactive sites with a decoy strained alkyne for 10 min; and 3) reaction of oxidized TePhe with BCN-fluorophore for 10 min. The first step was conducted under mildly basic (pH 8.5) conditions to promote deprotonation and alkylation of free thiols which can interfere with OSTAC through consumption of the oxidant and off-target thiol-yne reactivity. This step can be omitted for further timing savings but is recommended for minimizing background. Following alkylation, the sample was switched to a pH 5 sodium acetate buffer (low pH maximizes labeling intensity^28^). In the second step, oxidation was initiated; residual off-target functionalities which can react with BCN under oxidative conditions were blocked through the introduction of a non-fluorescent dibenzoazacycooctyne (DIBAC or DBCO) moiety, which reacts readily with off-targets while having extremely poor reactivity against oxidized TePhe.^28^ Finally, BCN-fluorophore addition resulted in the covalent attachment of fluorophores at TePhe sites. As the reaction time for OSTAC is very short^28^, we performed labeling by submerging slides into bulk reaction mix to ensure consistency between samples. Surface staining within a hydrophobic barrier, which reduced reagent consumption, also yielded good results.

Representative results of fluorescent labeling on vehicle and TePhe-treated (2 x 30 mg/kg over 8 hr) coronal sections are shown in Fig. 3a. Macroscopically, elevated fluorescence intensity was observed across brain regions in TePhe-treated samples relative to vehicle-treated controls (Fig. 3a, left panel). White matter regions and areas surrounding the ventricles was more prone to unspecific labeling by both OSTAC chemistry and immunofluorescence (IF). Nevertheless, TePhe incorporation at the single cell level was readily distinguishable from background fluorescence when imaging at higher magnification, as demonstrated by the “sprinkled” appearance of the neocortex, where OSTAC signal conformed to the shapes of presumptive neuronal cell bodies (Fig. 3a, centre). The widespread and even distribution of bright neurons suggests that TePhe readily permeated throughout brain structures (Fig. S15). At even higher magnification, signal along proximal neuronal projections could be discerned, while vehicle controls showed only a low, uniform fluorescence background (Fig. 3a, right; Fig. S16). We observed strong signal in hippocampal neurons (particularly in the CA3 subfield pyramidal neurons, Fig. S15e1, f1), consistent with known patterns from radioactive amino acid incorporation^51^ and single-cell analysis of translation states via Ribo-STAMP.^17^ Elevated signal in the choroid plexus of lateral ventricles was also observed, which can be rationalized on the basis of choroidal epithelia being exposed to and incorporating appreciable quantities of TePhe in its role as the blood-cerebrospinal fluid (CSF) barrier and secretor of CSF proteins (Fig. S15c2, d2).^52,53^ Of note, images shown in Fig. 3 were not subjected to background subtraction, demonstrating the excellent dynamic range of labeling, especially in gray matter regions.

**Figure 3.**
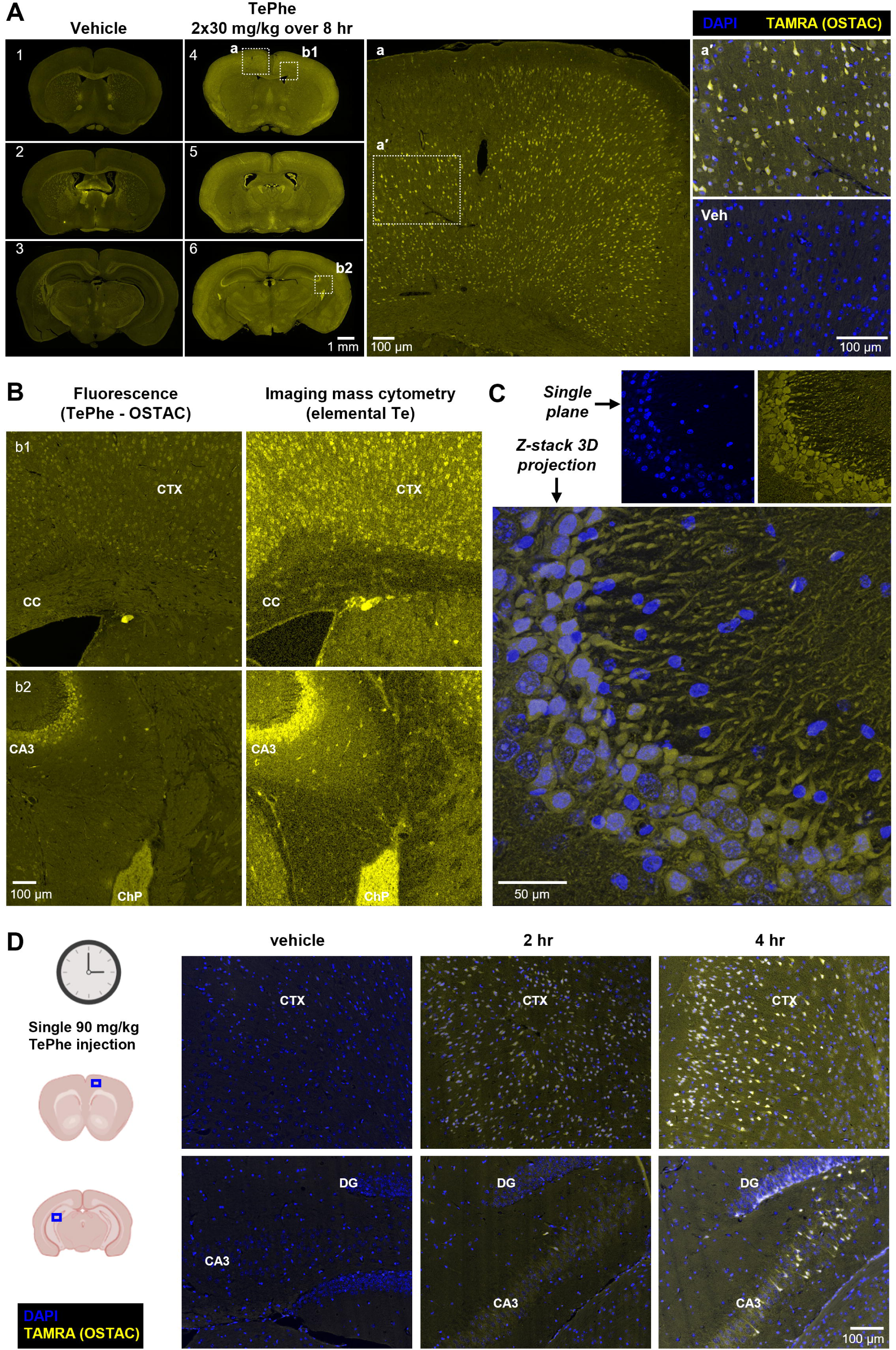
OSTAC-based fluorescent detection of protein synthesis in FFPE mouse brain sections. Images are from a 10-month old female mouse receiving 2x 30 mg/kg IP injections 4 hr apart and sacrificed at 8 hr. A) OSTAC detection of TePhe in 4 µm FFPE coronal sections. Left: whole section view showing OSTAC labeling background and signal (TAMRA channel only). Centre: region **a** of Section 4 magnified. Right: region **a’** of centre panel magnified, with overlay of TAMRA and DAPI channels. Matching region from vehicle Section 1 shown for comparison. B) Comparison of OSTAC labeling and imaging mass cytometry, using matched 1 mm^2^ ROIs from near-adjacent sections. Fluorescence: TAMRA intensity. IMC: Te intensity (geometric mean of ^126^Te, ^128^Te, ^130^Te; ^131^Xe-corrected). C) Confocal imaging of OSTAC-labeled CA3 subfield of hippocampus (20 µm section, z-stack with 0.4 µm stepsize). Abbreviation of brain substructures: Ctx = cortex; CC = corpus callosum, ChP = choroid plexus; CA3 = subfield 3 of Ammon’s horn. D) OSTAC labeling in 4 µm sections from 7-month old mice receiving a single dose of 90 mg/kg TePhe and sacrificed at short time points indicated above. CTX = cerebral cortex. DG = dentate gyrus. CA3 = subfield 3 of Ammon’s horn.

Fluorescent labeling patterns were corroborated by imaging mass cytometry (IMC) on a near-adjacent section (Fig. 3b, Fig. S17). IMC provides an orthogonal TePhe detection method through laser ablation, atomization and ionization of the specimen in an argon plasma, followed by mass spectrometric detection of elemental tellurium (Te).^21^ Fig. 3b shows 1 mm^2^ regions covering the cortex and corpus callosum (inset b1) and the area surrounding the CA3 subfield of the hippocampus (inset b2). Matching patterns of strong intensity in the cerebral cortex, hippocampus, and cells of the choroid plexus were observed, while fluorescence within the corpus callosum and white matter tracts were confirmed to be primarily originating from background fluorescence. Overall, IMC confirmed the presence of TePhe in the expected regions and validated the fluorescent readout.

As 4 µm sections span less than the width of a single neuronal soma^54^, we imaged OSTAC-labeled 20 µm sections to demonstrate greater depth of protein synthesis visualization (Fig. 3c, Fig. S18). Confocal imaging of the CA3 hippocampal subfield allowed generation of a 3-D projection in which the dense collection of CA3 pyramidal neuron cell bodies and proximal dendrites were readily visible. Our technique could therefore visualize protein synthesis while preserving the intricate 3-D architecture of neurons.

Finally, to determine the minimum duration of TePhe exposure required for imaging, brain sections from mice receiving a single 90 mg/kg IP dose and sacrificed at 2 and 4 hr post-injection were examined (Fig. 3d, Fig S19). At 2 hr, individual neurons with bright cell bodies were visible, consistent with TePhe incorporation by 2 hr in lysates (Fig. 2d). Very bright signal was observed by 4 hr. Despite the modest signal intensity indicated by spinal cord lysates of animals receiving a single TePhe injection (Fig. 2b), microscopy data demonstrated that single injection and short exposure regimes can yield strong signal within brain tissues due to spatially non-uniform incorporation. The durations investigated here were on par with the shortest labeling times reported for *in vivo* BONCAT-based protocols^16,55,56^. Our labeling strategy can thus effectively capture protein synthesis on short timescales, while forgoing the need for genetic or dietary manipulation. Taken together, these results demonstrate that TeACAT offers a facile and robust approach to the detection of protein synthesis in mouse brains.

### OSTAC chemistry can be multiplexed with detection of other biomarkers

We next demonstrated compatibility of fluorescent TePhe detection with the simultaneous immunofluorescence-based visualization of common cell markers within the brain (TeACAT-IF). OSTAC chemistry could be combined sequentially with Nissl dye staining of the somata of neurons (Fig. S20), which highlighted the enrichment of TePhe within neuronal cell bodies. We then tested sequential combination of OSTAC chemistry and IF: while OSTAC labeling is largely compatible with immunocytochemistry^28^, heat-induced antigen retrieval (HIAR) is needed to unmask epitopes in FFPE sections. TePhe was resilient toward HIAR conditions (20 min at 95 °C in pH 6 citrate or pH 9 Tris-EDTA antigen retrieval buffers; Fig. S21) and samples could be processed with routine antibody staining, with the integration of an OSTAC chemistry step between HIAR and blocking steps. To determine the effects of OSTAC conditions on subsequent antibody binding, we chose representative cell markers, including the post-mitotic neuronal marker NeuN, oligodendrocyte marker Olig2, activation marker c-Fos, and dendritic marker MAP2 (Fig. 4). The oxidative conditions used for OSTAC labeling did not abrogate expected IF patterns and NeuN, MAP2, and c-Fos staining were easily discerned (Figs. S22, 24-25). Only anti-Olig2 exhibited a notable decrease in immunofluorescence intensity after OSTAC treatment, but with preservation of its expected nuclear localization (Fig. S23), it remained serviceable for oligodendrocyte identification. This was consistent with previous observations that TePhe incorporation and the chemical environment needed for OSTAC coupling can both reduce and increase immunofluorescence intensity for select antigens^28^, possibly due to changes in epitope oxidation state or accessibility. Additional antibody targets tested and found to be minimally affected by OSTAC treatment include α-tubulin and eIF2α (Fig. S26-27). Across all antibodies tested, the specificity of antigen binding was preserved. Based on these results, we expect compatibility with IF, broadening the applicability of TeACAT for protein synthesis studies in complex tissues.

**Figure 4.**
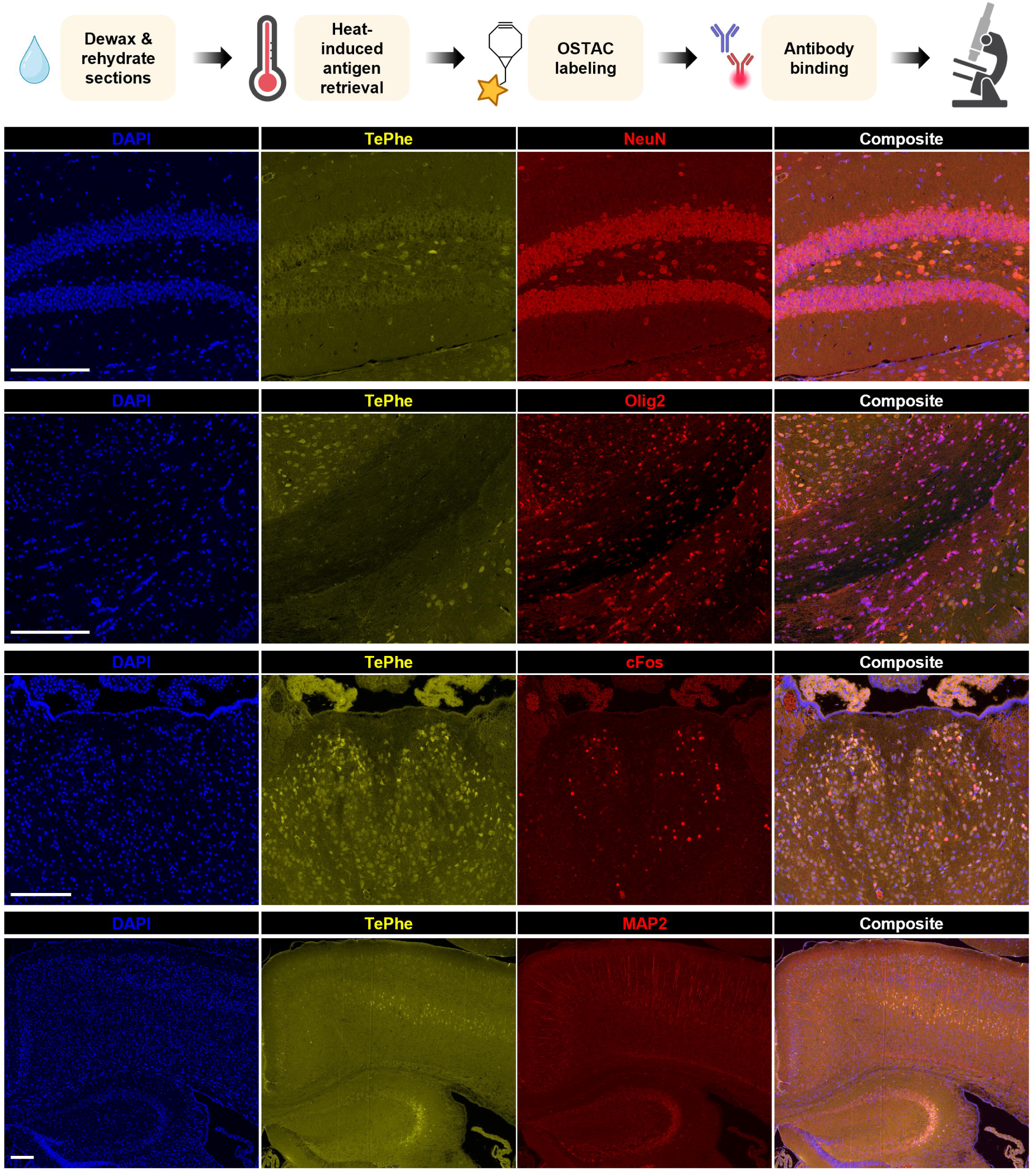
Combination of OSTAC labeling with select immunofluorescence targets within the mouse brain. TePhe detected in TAMRA channel; all antibody targets visualized by Dylight 650-conjugated 2 °C antibody. NeuN = neuronal nuclear protein, marker of post-mitotic neurons. Olig2 = oligodendrocyte transcription factor 2, oligodendroglial lineage marker. cFos = transcription factor associated with cell activity. MAP2 = microtubule-associated protein 2, marker of cytoskeletal structures within somata and dendrites of neurons. All scale bars = 100 µm.

We further used TeACAT-IF to validate patterns of TePhe incorporation within the brain. We focused on the subventricular zone (SVZ) as an example, which harbours neural stem cells capable of self-renewal and neurogenesis in adult mice. Baser et al. demonstrated that global protein synthesis rates fluctuate during neural differentiation *ex vivo,* as determined by O-propargyl-puromycin labeling of isolated cell populations (Fig. 5a).^57^ Here, we employed Sox2 as a marker for neural stem and progenitor cells and doublecortin (Dcx) as a marker for early neuroblasts, in conjunction with TePhe labeling, to visualize the protein synthesis timeline. Both Sox2 and Dcx staining were unperturbed when combined with OSTAC chemistry (Figs. S28-29). In tissues from 3-month old mice, a fraction of Sox2-positive cells co-localized with high TePhe signal (Fig. 5b-c); this is consistent with reports of increased protein synthesis in the subset of activated neural stem cells. In turn, less frequent co-localization of Dcx and TePhe signal was observed (Fig. 5b,d), consistent with reduced global protein synthesis and greater regulation of translation in neuroblasts^57^. We could thus corroborate published findings^57^ within intact tissues, which was not possible with established techniques due to limited permeation within the brain.^32^

**Figure 5.**
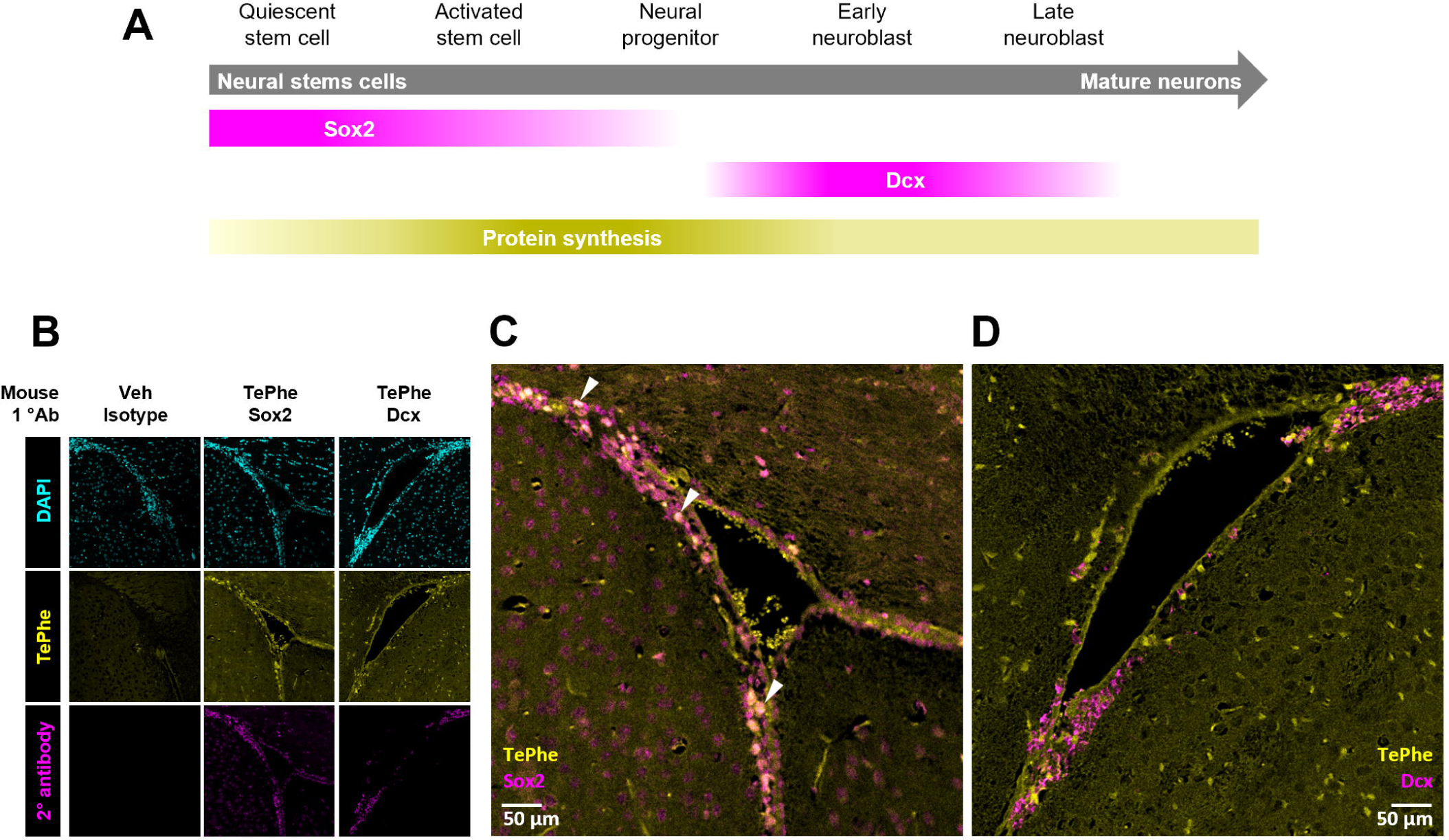
Combination of OSTAC chemistry and immunofluorescence for visualizing protein synthesis in neurogenic niche (subventricular zone) in adult mouse brains. A) Overview of expected patterns of protein synthesis, Sox2 and Dcx expression during adult neurogenesis. B) Patterns of protein synthesis (TePhe signal) combined with Sox2 and Dcx staining. Images of overlaid TePhe with Sox2 (C) and Dcx (D) respectively. White arrowheads in C) highlight co-localization between TePhe and Sox2 signal.

### TePhe-tagged proteins can be enriched and identified using proteomics

TeACAT can readily be adapted for affinity tagging: we previously demonstrated biotinylation of TePhe for visualization with fluorescent streptavidin conjugates in fixed cell cultures.^28^ We therefore explored TeACAT in conjunction with mass spectrometry (MS)-based proteomics (workflow shown in Fig. S30, TeACAT-MS). The reagents used were largely identical to in-solution fluorescent labeling, but with TePhe coupling to a BCN-biotin sulfone probe, where the thioether of biotin is pre-oxidized to avoid formation of heterogeneous sulfoxides during the OSTAC reaction.^28^ Briefly, the steps included sample pre-treatment by standard DTT reduction and iodoacetamide alkylation, acetone precipitation to remove excess reductant, biotin conjugation to TePhe, and a second round of acetone precipitation to remove excess biotin probe prior to streptavidin complexation. Routine conditions for streptavidin capture, digest, and peptide clean-up were then used. TePhe is expected to replace only a small fraction of Phe sites across the proteome, enabling pull-down of newly synthesized proteins while the majority of peptides remain unmodified for MS analysis.

We began by testing our workflow on TePhe-treated HEK293T cell lysates. TeACAT-MS led to the detection of 3831 protein groups detected in TePhe-treated vs. 2470 in untreated samples (Fig. S31a). While the number of protein groups detected in the negative control was comparable, the majority (77%) exhibited significantly increased (≥2-fold change, *p* <0.05) intensity in the TePhe condition, indicating enrichment of newly synthesized proteins over background (Fig. S31b). The degree of enrichment did not correlate with Phe content or sequence length (*R*^2^ = 0.058 and 0.107, respectively; Fig. S31c).

We functionally validated this workflow through the characterization of newly synthesized proteomes in undifferentiated and differentiating neuroblasts. Working concentrations of TePhe did not impact on Neuro2a (N2a) mouse neuroblastoma cell viability (Fig. S32a). N2A cells were differentiated for 4 days, after which protein synthesis over an 8-hr window was captured through addition of 400 µM TePhe. OSTAC-based enrichment was used to capture the newly synthesized proteomes, resulting in quantification of 2314 protein groups across differentiated vs undifferentiated N2a samples (LFQ; Fig. S32b). Applying cutoffs (≥1.5-fold change, *p*-value <0.05) resulted in 195 (8.4%) down-regulated and 268 (11.6%) up-regulated protein groups (Fig. 6a). Gene ontology analysis showed a clear down-regulation of proteins involved in DNA organization and replication, as expected for neurons exiting the mitotic cycle. Transport and lipid metabolism-associated proteins were enriched in the up-regulated set (Fig. 6b, Fig. S32c). The ability to discern these expected biological changes encouraged us to apply our enrichment workflow to mouse brain proteomes.

**Figure 6.**
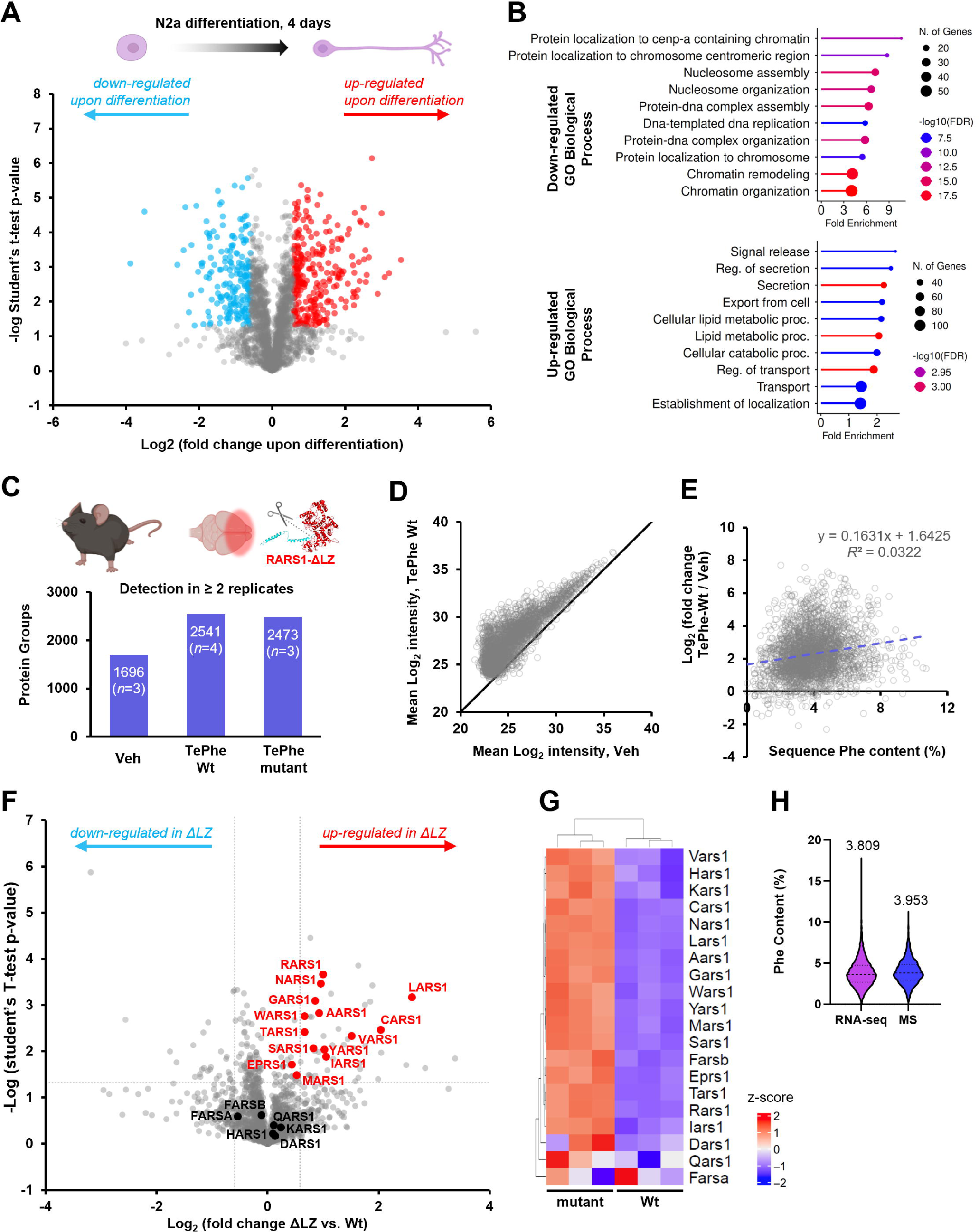
Identification of newly synthesized proteins via TeACAT. A) Comparison of newly synthesized proteomes of undifferentiated and 4-day-differentiated murine neuroblastoma (N2a) cells, tagged by 400 µM TePhe treatment for 8 hr. Protein groups passing LFQ criteria in at least 2 replicates in any treatment group were included. Student’s t-test was performed after imputation; significance cutoffs at fold change >1.5 and *p* <0.05. B) Pathway analysis of significantly down- and up-regulated proteins from A), performed using ShinyGo 0.85. Top 10 most enriched pathways with FDR <0.05 are shown. C) OSTAC-based enrichment for newly synthesized proteins from mouse forebrains. Wt = animals expressing normal, non-truncated RARS1; mutant = homozygous truncation of RARS1. Veh = vehicle-treated, Wt mice. TePhe = mice receiving 2 x 45 mg/kg injections 4 hr apart. Numbers of protein groups identified in at least 2 replicates per group are shown. D) Comparison of mean MS/MS intensities of protein groups detected across Veh and TePhe-treated Wt samples, showing enrichment in the TePhe sample. E) Relationship between sequence Phe content and degree of enrichment, showing no bias for proteins with more Phe sites. F) Comparison of newly synthesized proteins in the forebrains of Wt and mutant mice. Cytosolic aminoacyl-tRNA synthetases are highlighted. G) Differential expression analysis of Wt and ΔLZ transcriptomes showing up-regulation of mRNAs for most cytosolic aminoacyl-tRNA synthetases.^58^ H) Comparison of Phe content of all proteins expected from coding regions identified by RNA-seq against all proteins identified by OSTAC-based enrichment, showing highly similar distributions

TeACAT-MS was ultimately applied to a mouse model of hypomyelinating leukodystrophy caused by truncation of arginyl-tRNA synthetase, RARS1, in the forebrain.^58^ Bulk TePhe incorporation did not differ significantly between control and mutant animals, suggesting that global protein synthesis was not impaired (Fig. S33). To investigate differences in the active synthesis of specific proteins, we injected control littermates and mutant mice with 2 doses of 45 mg/kg TePhe at 4 hr intervals and harvested forebrain tissue at 8 hr. Increased numbers and intensities of proteins in TePhe-treated samples over vehicle controls confirmed enrichment of tagged proteins (Fig. 6c-d). As with HEK293T proteomes, enrichment showed no bias for Phe content (Fig. 6e, *R*^2^ = 0.03; 3.8% median Phe in TeACAT-MS proteome vs 3.5% for the full mouse proteome).^59^ Additionally, comparison of Phe content across proteins identified by TeACAT-MS with proteins expected from all coding sequences detected in RNA-seq yielded highly similar distributions (Fig. 6h).

We identified 27 down-regulated and 47 up-regulated proteins (≥1.5-fold change, *p* <0.05) out of 2006 identified protein groups. Strikingly, 12 cytosolic aminoacyl-tRNA synthetases (ARSs) were up-regulated (Fig. 6f). These findings complement transcriptome data, where most cytosolic ARSs were upregulated in mutant animal brains (Fig. 6g).^58^ Conversely, the 3 ARSs, which were unchanged on RNA level, were also unchanged in proteomics. This included the phenylalanyl-tRNA synthetase α subunit (FARSA). The phenylalanyl-tRNA synthetase β subunit (FARSB) showed unchanged proteomic abundance despite mRNA up-regulation, which may reflect instability of excess FARSB in the absence of a corresponding up-regulation of its binding partner FARSA.^60,61^ (This consistency in FARSA/B levels was fortuitous as equal TePhe charging is needed for effective comparison of active translation between the two groups.)

Taken together, we demonstrate that TeACAT allowed capture and direct characterization of newly synthesized mouse brain proteomes and could discern pathologic changes in disease states. As is, we expect our method to be applicable to a diverse range of models and biological questions.

## Discussion

In this work, we established a methodology for robust and systemic *in vivo* tagging of newly synthesized proteins followed by their fluorescent detection or proteomic identification. We could visualize protein synthesis in brain sections and concomitantly identify cell type. Newly synthesized proteins could be biotinylated and enriched from the bulk proteome for sequence-level characterization, allowing comparison of protein synthesis in distinct biological states. The facile implementation of TeACAT in otherwise unmanipulated animals opens many possibilities for studies in complex systems where protein synthesis is currently challenging to interrogate.

A key feature of TeACAT is the high structural similarity between TePhe and Phe, resulting in effective activation of TePhe by endogenous translation machinery. This is advantageous when compared to azide-alkyne cycloaddition-based BONCAT probes such as AHA and homopropargylglycine (HPG). Endogenous methionyl-tRNA synthetase processes AHA and HPG significantly less efficiently than the cognate substrate^62^, necessitating genetic engineering or canonical amino acid depletion to boost probe incorporation. As TePhe effectively competes with Phe, protein synthesis can be studied under more physiological conditions and with minimal effort. In contrast to amino acid isotopologues (e.g. radiotracers^63–66^, heavy amino acids for MS^67–70^, and deuterated Raman probes^71–73^), which are by definition chemically equivalent to canonical amino acids and efficiently incorporated, TePhe can be efficiently functionalized with OSTAC chemistry.

TePhe is well-tolerated in mice. The highest single dose, 90 mg/kg TePhe, resulted in a spike in serum concentration to approximately 300 µM. This is multiple times the expected concentration of Phe in blood and favours uptake and protein incorporation, but is not so high as to cause concerns of phenylketonuria (>1 mM Phe).^74,75^ Thorough characterization on both transcriptomic and proteomic levels revealed essentially no perturbation of brain physiology after 2 x 45 mg/kg injections over 8 hr. Only a single gene was found to be significantly up-regulated by RNAseq: Cdkn1a, encoding the p21 protein involved in inhibition of cell cycle progression in response to DNA damage, at a 2.1-fold increase. Excess Phe can induce p21 expression in cell culture^76^, and given the absence of other indicators of DNA damage, we assume that p21 is upregulated due to the sudden influx of a Phe-like molecule, rather than due to DNA damage. No gross morphological indications of damage were found in any of the studied organs (Fig. S1, S10), positioning TeACAT as a minimally disruptive method to study protein synthesis.

Our data suggests that TeACAT is uniquely suited for probing protein synthesis in complex tissues, such as for neuroscience questions: TePhe was efficiently transported into the central nervous system after IP injection and diffused across brain regions, circumventing the need for specialized procedures, such as intracranial injections. TePhe uptake may be aided by the relatively low concentration of Phe in blood^77^, as well as high affinity of LAT1 (L-type amino acid transporter 1) for Phe and Phe-like substrates^78,79^ facilitating TePhe transport across the blood-brain barrier. Additionally, TeACAT enables experimentation ranging from 2 hr to several days, allowing processes occurring on varying timescales to be examined. The shortest labeling period is on par with the best temporal resolution achieved with BONCAT.^56^

TeACAT provides a straightforward workflow for capturing spatial patterns of global protein synthesis in complex tissues. OSTAC chemistry could be easily integrated with routine tissue processing and immunofluorescence workflows for combined visualization of protein synthesis and cellular biomarkers at high spatial resolution. Steps unique to fluorescent OSTAC labeling can be performed in under 1 hr. Our optimized protocol for on-slide, immersion staining generated equally clean results on 4 µm and 20 µm FFPE sections, and can most likely be extended to even thicker tissues. Compared to mass cytometry-based methods, fluorescence imaging benefits from accessible instrumentation, superior spatial resolution, and quicker imaging of large areas of tissues, which allows for whole mount imaging. Laser ablation of the sample in IMC in turn limits resolution and imaging area (typically 1 hr acquisition time for 1 mm^2^ imaged at 1 µm resolution).^26,27^ In turn, IMC retains the advantage of lower background due to the absence of autofluorescence, the ability to be multiplexed, and the potential of using isotopologues in pulsed experiments^80^.

Furthermore, TeACAT is compatible with other methods: spatial patterns generated by TeACAT can complement data from sequencing-based methods, such as RiboTag^15^ and Ribo-STAMP.^17^, where spatial connectivity might be lost during sample preparation. TeACAT provides protein level information to match spatially resolved methods of mRNA translation such as RIBOmap.^81^ Additionally, we expect that TeACAT can be multiplexed with other biorthogonal labeling techniques, namely BONCAT^19^ and THRONCAT^82^, as OSTAC chemistry can be performed consecutively with azide-alkyne cycloaddition.^28^

While TeACAT-IF allows for visualization of protein synthesis patterns and the unbiased identification of cell types with varying TePhe incorporation, TeACAT combined with MS enables identification and quantitation of proteins synthesized during a defined treatment period. Our experiments in a murine leukodystrophy model demonstrated that TeACAT can effectively capture changes in translational output for individual proteins. Chen et al. have previously adapted OSTAC chemistry to enrich lipidated proteins for MS analysis^83^, confirming the versatility of tellurophenes as proteomic probes. We anticipate that TeACAT-MS will be highly effective for uncovering molecular differences between healthy and disease states, as well as responses to environmental or developmental cues.

To conclude, TeACAT is a protein synthesis tracking methodology suitable for complex biological systems. We see TeACAT not only as a standalone methodology with ideal attributes for *in vivo* studies, but also a ‘plug-and-play’ method that can be easily integrated to address diverse and complex research questions.

## Methods

### Reagents

TePhe is currently licensed to Standard BioTools for mass cytometry. It is available from the authors upon request. Large-scale TePhe synthesis was achieved by modification of existing protocols^21^ (for details, see Supplementary Information, Section 1). “BCN-TAMRA” refers to either in-house synthesized BCN-PEG3-TAMRA^28^ or commercial BCN-PEG4-TAMRA purchased from AAT Bioquest through CedarLane (70512(AAT)); the two probes were used interchangeably. BCN-PEG3-biotin sulfone (“BCN-BtnSO2”) was synthesized in-house.^28^ DBCO-PEG4-OH (“DBCO”, Click Chem Tools), Captisol® (CyDex Pharmaceuticals), *N-*chlorosuccinimide and organic solvents (Sigma), buffers (Bioshop) and TLC plates (Santai) were all used as received from commercial suppliers.

### Animals

Animals were housed in a temperature-controlled facility with a 12-hr day-night cycle and given access to food (LabDiet rodent chow 5001) and water *ad libitum*. All mice are of the C56BL/6J background. For studies on TePhe uptake and TeACAT, individuals of different sexes were evenly distributed among conditions to minimize confounding factors. Male and female animals between 2.5 to 11 months old were used. For RARS1 truncation mutants, animals in which one or both copies of exon 2 of the *Rars1* gene were flanked by LoxP sequences were used, which were crossed with animals expressing Cre recombinase from a forebrain-specific promoter (*Emx1^tm1(cre)Krj^*). A detailed table of animal age, sex, and genotypes is provided in the Supplementary Information, Section 2. All experiments were conducted in accordance with University of Toronto’s animal use protocol 20012885.

### TePhe injections

TePhe was dissolved at up to 18 mg/mL in a solution of 40% Captisol in PBS. The solution was sterilized through a 0.22 µm filter, aliquoted, and stored at -80 °C until use. IV injections were limited at <100 µL volume per mouse delivered through the tail vein, while IP injections ranged between 90 and 150 µL per injection, depending on the weight of the mouse. Repeated IP injections were given on alternating sides of the abdomen. Animals were returned to their home cage after injection and monitored for any indications of discomfort.

### Tissue/biofluid collection and processing

#### Tissue collection

Animals were euthanized through CO2 inhalation. 30 sec after cessation of breathing, secondary euthanasia was performed by cervical dislocation (except in cases of blood collection). Animals were not perfused. After decapitation, whole brains were collected for fixation, with the exceptions of olfactory bulb tissue and the segment of spinal cord immediately below the brainstem, which were separately flash frozen. Other major organs were dissected and rinsed in PBS prior to fixation or flash freezing. Drop fixation was performed in 4% *p-*formaldehyde in PBS at 4 °C with gentle rocking. Fresh fixative was replenished 24 hr after initial immersion of tissues. Whole brain and the largest lobe of the liver were fixed for up to 72 hr, while smaller organs were fixed for up to 48 hr. For RNA-seq and bulk proteomics, freshly harvested whole brains were cut into coronal slices using a 2 mm stainless steel brain matrix (Zivic Instruments 5325). Sections were individually stored in 1.5 mL tubes, immediately flash frozen in liquid nitrogen, and stored at -80 °C until use. For OSTAC-based proteomics of Wt and ΔLZ forebrains, the first 1 mm-thick coronal slice from the rostral end of the brain (excluding olfactory bulb) was dissected.

#### Blood collection

Terminal blood collection was performed after primary euthanasia by CO2 inhalation. The animal’s chest cavity was opened and blood drawn by cardiac puncture. Whole blood was transferred to a 1.5 mL tube and centrifuged at 1500 x g for 3 min at RT. Separated blood was allowed to stand at room temperature (RT) for 5 min, after which the clear top layer was transferred to a new tube, immediately frozen on dry ice, and stored at -80 °C until assayed. Assessment of serum creatinine, blood urea nitrogen, and aminotransferase levels were performed by The Centre for Phenogenomics (Toronto, Canada) using standard protocols.

#### Urine collection

Urine samples were collected by placing animals onto the lid of a sterile 15 cm petri dish and performing routine restraining motions. Most animals produced appreciable volumes (40-100 µL) of urine during this brief handling process. The urine was transferred to a 1.5 mL tube by pipet, frozen on dry ice, and stored at -80 °C until assayed. Animals were then returned to home cages if urine collection at later time points were desired, or sacrificed to collect matched blood samples.

#### Tissue lysis in preparation for in-solution OSTAC labeling

Spinal cord samples were thawed for 2-3 min at RT, briefly crushed with a pipet tip, suspended in lysis buffer (PBS + 0.5% SDS + complete mini protease inhibitor) and homogenized using a syringe fitted with a 22-gauge hypodermic needle. All other tissues were homogenized by mashing partially thawed tissue against a cell strainer (70 µm mesh, placed over a 50 mL falcon tube) using the back end of a syringe plunger, suspending the resulting paste in lysis buffer, and passing the mixture through the strainer by a combination of gravity and aspiration. The thick mixture was further homogenized by being drawn through a 22-gauge needle, transferred to microcentrifuge tubes, and pelleted at 2000 x g for 1 min to remove large debris. Protein concentration was determined by BCA assay (Pierce #23225) according to kit instructions in 96-well plates and read using a BioTek Synergy H1 microplate reader. Bulk lysates were stored at -80 °C until use.

#### Frozen sections

After fixation, tissues were rinsed in PBS, transferred to sterile 30% sucrose in PBS, and incubated at 4 °C until tissues lost buoyancy. Samples were then embedded in OCT and sectioned on a Thermo Scientific HM525 NX Cryostat.

#### FFPE sections

After fixation, tissues were washed in PBS (2 × 30 min) and, if necessary, cut into smaller pieces prior to dehydration. Whole brain was divided into 2 mm coronal slices. Kidney was split into two coronal halves; the largest lobe of the liver was divided into 4-5 segments by cuts along the short axis of the tissue. Tissues were washed with deionized water for at least 30 min, then dehydrated on a rocking platform using the following changes of solvents, for at least the durations listed: 70% ethanol (EtOH, overnight), 80% EtOH (30, 40, 40 min), 90% EtOH (30, 40, 40 min), 100% EtOH, (30, 40, 30 min), 1:1 EtOH/xylenes (30 min, 1 hr), and 100% xylenes (2 × 45 min). Hot wax infiltration was performed at 60 °C in a Leica TP 1020 tissue processor (30, 45, 45 min). Tissues were embedded in Paraplast Plus and sectioned at 4 µm thickness using a Leica rotary microtome. Select brain tissues were processed and sectioned by UHN STTARR histopathology services (images shown in Figs. 4, 5, S14-17, S19-25).

### Protein expression and purification

Chemically competent *Escherichia coli* BL21(DE3) cells (Agilent Technologies) were transformed with 60 ng plasmid DNA encoding codon-optimized human FARS2 residues 37–451, lacking the predicted N-terminal mitochondrial targeting sequence and fused to an N-terminal hexahistidine tag, in a pET-28a-derived expression vector (Twist Bioscience). Briefly, 1.5 µL DNA was added to competent cells and incubated on ice for 30 min, followed by heat shock at 42°C for 30 s. Cells were recovered in 700 µL SOC medium at 37°C for 60 min, plated on LB agar supplemented with 50 µg/mL kanamycin (BioShop), and incubated overnight at 37°C.

A single colony was inoculated into 200 mL Terrific Broth (TB; BioShop) containing 50 µg/mL kanamycin and cultured overnight at 37°C with shaking at 180 rpm. The pre-culture was used to inoculate 2 L TB supplemented with 50 µg/mL kanamycin. Cultures were grown at 37°C and 200 rpm to an OD₆₀₀ of 0.7–0.8, after which expression was induced with isopropyl β-D-1-thiogalactopyranoside (IPTG; BioShop) to a final concentration of 1 mM. Cultures were incubated overnight at 21°C and 200 rpm, harvested by centrifugation at 3,900 x g for 45 min at 4°C, and stored at −20°C until purification. Samples collected before and after induction, normalized to an OD₆₀₀ of 1.0, were retained for SDS-PAGE analysis.

Cell pellets derived from 2 L culture were resuspended in 40 mL ice-cold lysis buffer consisting of 40 mM HEPES, pH 8.0, 300 mM NaCl, and 5 mM imidazole. Cells were lysed using an EmulsiFlex-C3 microfluidizer (Avestin) at 15,000 psi for two passes with cooling between passes. The lysate was clarified by centrifugation at 10,200 x g for 60 min at 4°C.

The clarified lysate was applied to a 5 mL Ni-NTA agarose column (Qiagen) pre-equilibrated with lysis buffer. The column was washed with 5 column volumes (CV) of lysis buffer, and bound protein was eluted stepwise using 30, 55, 100, 200, 300, and 500 mM imidazole in 40 mM HEPES, pH 8.0, and 300 mM NaCl. Fractions containing FARS2 were identified by SDS-PAGE and pooled.

Pooled Ni-NTA fractions were dialyzed overnight at 4°C against 4 L low-salt buffer containing 20 mM HEPES, pH 8.6, 20 mM NaCl, 7 mM MgCl₂, and 10% (v/v) glycerol using cellulose dialysis tubing with a 14 kDa molecular-weight cutoff (Sigma-Aldrich). Protein was loaded onto a 5 mL HiTrap Q HP column (Cytiva) equilibrated with low-salt buffer using an ÄKTA pure™ chromatography system (Cytiva). After washing with 5 CV of low-salt buffer, protein was eluted over 10 CV with a linear gradient to 50% high-salt buffer containing 20 mM HEPES, pH 8.6, 2 M NaCl, 7 mM MgCl₂, and 10% (v/v) glycerol at a flow rate of 2.5 mL/min. Fractions containing FARS2 were identified by SDS-PAGE and pooled.

Pooled anion-exchange fractions were concentrated to 5 mL using a 30 kDa molecular-weight-cutoff centrifugal concentrator (Amicon Ultra; Millipore) and loaded onto a HiLoad 16/600 Superdex 200 pg size-exclusion column (Cytiva) equilibrated with SEC buffer containing 20 mM HEPES, pH 8.0, 200 mM NaCl, 5 mM MgCl₂, and 5% (v/v) glycerol. Size-exclusion chromatography was performed on an ÄKTA pure™ system at 0.5 mL/min. Fractions corresponding to the main elution peak were pooled. Protein concentration was determined by absorbance at 280 nm using a NanoDrop ND-1000 spectrophotometer (Thermo Fisher Scientific) and a calculated extinction coefficient of 66.73×10^3^ M^−1^⋅cm^−1^. Purified protein was aliquoted, flash-frozen in liquid nitrogen, and stored at −80°C.

### ATP-consumption assay

ATP consumption by FARS2 was measured using the Kinase-Glo® Luminescent Kinase Assay (Promega) as an endpoint measure of residual ATP. Reactions were performed in assay buffer containing 50 mM HEPES, 50 mM KCl, and 10 mM MgCl₂. To reduce ATP turnover by FARS2 in the absence of amino acid, a 10× FARS2 stock containing 10 µM FARS2 and 10 µM ATP in assay buffer was pre-incubated at 37°C for 30 min. Unless otherwise indicated, reactions contained 10 µM ATP, 1 µM FARS2, and 62.5 µM of the indicated amino acid. Control reactions lacking FARS2 and/or the amino acid were included. Reaction mixtures were incubated at 37°C for 30 min.

Following incubation, 20 µL aliquots of each reaction were transferred to white, flat-bottom 96-well plates containing 20 µL Kinase-Glo reagent. Plates were centrifuged at 3,900 × *g* for 1 min and incubated at room temperature for 10 min, protected from light, before measurement using a Synergy H1 microplate reader (BioTek) with a 1 s integration time. ATP standards containing 2.5, 5, and 10 µM ATP were included on each plate. Residual ATP concentrations were determined by interpolation from the ATP standard curve, and ATP consumption was calculated as the difference between the initial and residual ATP concentrations.

### RNA-seq

RNA extraction was performed on 2 mm coronal brain slices (∼60 mg) using the Qiagen miRNeasy mini kit according to manufacturer’s instructions. Buffer volumes were scaled up 2-fold and contents of each brain slice distributed over 2 columns to ensure adequate binding capacity. RNA integrity was assessed using Bioanalyzer through Centre for Applied Genomics, at the Hospital for Sick Children, Toronto, Canada.

RNA-seq libraries were constructed using the NEBNext® Ultra™ II RNA Library Prep with Sample Purification Beads (NEB #E7775S) following the manufacturer’s instructions. The USER enzyme (NEB #M5505S) was added during adapter ligation to resolve the hairpin-loop adaptor design, ensuring efficient downstream amplification. Finalized libraries were sequenced at the Centre for Applied Genomics, at the Hospital for Sick Children, Toronto, Canada on an Illumina NovaSeq 6000 SP platform to produce ∼30 million paired-end reads of 150 nucleotides per sample. Data processing was performed on the Trillium supercomputing cluster (hosted by Scinet and the Digital Research Alliance of Canada). Analysis and visualization were performed in RStudio. Reads were mapped to Mus_musculus.GRCm39.114^84^ using STAR aligner^85^. Counts were obtained using RSubread’s FeatureCount function (Version 2.26.0)^86^ and differentially expressed genes were identified with DESeq2 1.52.0^87^. Volcano plots were exported with EnhancedVolcano 1.30.0.

### Fluorescent OSTAC labeling

#### General considerations for OSTAC chemistry

NCS stock solutions (50 mM in dH2O) were freshly prepared for each OSTAC experiment. DBCO-PEG4-OH was prepared as a 50 mM stock in DMSO and stored at -20 °C. BCN-TAMRA was prepared as 5 mM stocks in DMSO and stored at -80 °C. Due to the short duration of OSTAC blocking and labeling steps, a 30-second stagger in start times between samples was employed to ensure sufficient time for handling. Unless otherwise stated, all steps were performed at RT.

#### In-solution fluorescent OSTAC chemistry of tissue lysates

One of two types of pre-treatment leading up to OSTAC labeling was performed: for 1-pot labeling, samples were normalized to 1.0 mg/mL protein by dilution in PBS + 0.5% SDS and mixed with EDTA (3 mM). Free thiols were alkylated with iodoacetamide (10 mM) in the dark for 20 min, after which samples were used directly for OSTAC labeling. For 2-step labeling with DTT reduction, samples were normalized to a common concentration between 1.0 to 2.0 mg/mL by dilution in PBS + 0.5% SDS, followed by 2-fold dilution with 200 mM Tris + 200 mM NaCl, pH 8.5 + 0.5% SDS (150 to 300 µg input). Samples were reduced with DTT (6 mM) at 55 °C for 45 min, cooled, alkylated with iodoacetamide (15 mM) in the dark for 25 min, and quenched with additional DTT (6 mM). To remove excess DTT and iodoacetamide, proteins were precipitated by addition of 4-fold volume of reagent grade acetone (1 hr to overnight). Precipitates were collected at 18000 × g for 5 min, resuspended in at least 3 initial volumes of pure acetone by sonication in a Transsonic 420 bath sonicator for 10-15 seconds. The process was repeated for at least 3 acetone washes. Pellets were air-dried until no flowing solvent was visible upon inversion of the tube, redissolved in PBS + 0.5% SDS, quantified and adjusted to 1.0 mg/mL. To initiate the OSTAC labeling sequence, DBCO-PEG4-OH (150 µM) and NCS (500 µM) were added in quick succession and the solution immediately vortexed. After exactly 10 min, BCN-TAMRA (1 to 2 µM) and additional NCS (100 µM) were added in quick succession, the solution vortexed, and again incubated for exactly 10 min. Reactions were quenched by the addition of 5x Laemmli loading buffer (300 mM Tris pH 6.8, 50% v/v glycerol, 10% w/v SDS, 25% v/v β-mercaptoethanol, trace bromophenol blue) and boiled for 1 min. Samples were fractionated on home-made 1.5 mm thick 10% polyacrylamide gels at 120 V for 45-60 min.

#### OSTAC-based determination of free TePhe in mouse serum

TePhe standards in serum were generated by mixing concentrated TePhe and serum from untreated mice at a ratio of 1:20 to attain final concentrations of 0 to 450 µM. Serum from TePhe injected mice was used directly. All standards and samples were mixed with 40 µL methanol at RT, vortexed, and pelleted at 13000 × g for 10 min to remove proteins. 5 µL of cleared supernatant was transferred to a new 1.5 mL tube and allowed to air-dry in a fume hood for 1.5 to 2 hr. The residue was redissolved in 10 µL PBS containing BCN-PEG3-TAMRA (25 µM), followed by addition of NCS (1.25 mM) to initiate OSTAC labeling. After 15 min, the reaction was quenched by addition of DTT (3 mM). The reaction mixture was acidified by addition of 20% acetic acid solution (1.5 µL) and spotted onto TLC plates. Plates were allowed to air-dry for 10 min before developing twice in 15% MeOH in DCM + 3% AcOH. Plates were dried briefly after removal from the development chamber and immediately imaged. Densitometry was performed using GelAnalyzer. To account for variations during precipitation, 2 technical replicates were processed per sample.

#### OSTAC-based analysis of TePhe-derived metabolites in mouse urine

Standards were generated by mixing concentrated TePhe and urine from untreated mice to attain final concentrations of 0 to 400 µM TePhe. Urine from treated mice was used directly. Standards and samples were precipitated with methanol as described for mouse serum. Due to the high concentrations of OSTAC-reactive metabolites in TePhe-treated samples, solutions were diluted by a further 2 to 4-fold in PBS before proceeding to OSTAC labeling and TLC analysis as described above. To collect OSTAC adducts of putative TePhe metabolites for mass spectrometric analysis, the labeling reaction was performed using the equivalent of 2 µL urine collected at 2 hr post-injection. After methanol precipitation as described above, 10 µL cleared supernatant was diluted to 200 µL in 10 mM sodium acetate buffer, pH 5 and treated with BCN-TAMRA (50 µM) + NCS (1 mM) for 20 min. The reaction was then diluted 2-fold in H2O and acidified with 0.5% formic acid. The mixture was loaded onto a C18 disk (Empore) which had been activated with methanol and 70% acetonitrile + 0.1% formic acid in H2O, and equilibrated in 0.1% formic acid in H2O. Adducts were eluted using a gradient of 5, 10, 15, 20, 25, 33, and 50 % acetonitrile + 0.1% formic acid in H2O. Concentrated metabolite adduct fractions were used directly for electrospray ionization mass spectrometry analysis. Spectra were acquired on an Agilent 6538 UHD hybrid quadrupole time-of-flight mass spectrometer in positive mode.

#### Fluorescent OSTAC labeling of FFPE tissues

Liver and small intestine cryosections were thawed at RT for up to 30 min, followed by rehydration in PBS (3 x 3 min), permeabilization in 0.3% Triton X-100 in PBS (2 x 30 min), and brief washes in PBS (2 x 1 min) before proceeding to OSTAC labeling. FFPE brain sections were baked at 60-65 °C for 30 min before dewaxing in xylenes (2 x 5 min), followed by rehydration in 100% EtOH (2 x 5 min), 90% EtOH (2 x 5 min), 80% EtOH (2 x 5 min), 70% EtOH (2 x 5 min), and dH2O (5 min) before proceeding to OSTAC labeling or heat-induced antigen retrieval (see below). OSTAC staining was performed in glass coplin jars with slides vertically immersed, with multiple jars used to facilitate rapid transfer of slides from one solution to the next. Rehydrated tissues were equilibrated in alkylation buffer (100 mM Tris + 100 mM NaCl, pH 8.5 + 0.025% Triton X-100) for 5 min, followed by iodoacetamide (10 mM) in alkylation buffer for 20 min in the dark (this step is optional). Sections were briefly rinsed in PBS, then equilibrated in OSTAC buffer (10 mM sodium acetate, pH 5 + 0.025% Triton X-100) for 5 min. OSTAC labeling consisted of blocking with DBCO (100 µM) + NCS (500 µM) in OSTAC buffer for 10 min, followed by labeling with BCN-TAMRA (1.5 µM) + NCS (100 µM) in OSTAC buffer for 10 min (reaction solutions prepared immediately before use). Labeling was stopped by 3 rapid washes in PBS + 0.025% Triton X-100 and 2 washes in PBS. Sections were then subjected to blocking and antibody staining if desired (see below). Lastly, sections were stained with DAPI (2.5 µg/mL in PBS, 10 min, dark), washed with PBS (2 x 10 seconds), and mounted in ProLong Gold antifade (Invitrogen).

#### Heat-induced antigen retrieval (HIAR)

Rehydrated FFPE sections were equilibrated in antigen retrieval buffer (10 mM Tris + 1 mM EDTA, pH 9 + 0.05% Tween-20, or 10 mM sodium citrate, pH 6 + 0.05% Tween-20) for 5 min, then transferred to antigen retrieval buffer pre-equilibrated to to 95 °C in a water bath. Slides were maintained at 95 °C for up to 30 min, at which point heated jars were removed from the bath and allowed to cool at ambient temperature for 15-30 min, followed by further cooling within an RT water bath for up to 30 min. Slides were washed once with PBS, and either subjected to OSTAC labeling as described above, or subjected directly to blocking and antibody binding in the case of staining pattern controls.

#### Antibody binding

All steps were performed at RT. Sections were blocked using immunofluorescence buffer (IF buffer; 1% BSA, 2% FBS in PBS or TBS) for at least 1.5 hr. Antibodies were diluted in 1:1 IF buffer/PBS (or TBS) and centrifuged at 13000 x g for 5 min to remove aggregates prior to use. Primary antibodies were allowed to bind overnight, followed by PBS or TBS wash (2 x 3 min), application of secondary antibodies for 2 hr, and PBS or TBS wash (2 x 3 min). Sections were DAPI stained and mounted as described above prior to imaging. A table of antibody suppliers, lot numbers and working concentrations can be found in the Supplementary Information, Section 3.

### Fluorescence imaging and analysis

#### Gel imaging

SDS-PAGE gels were imaged using a Syngene G:BOX Chemi XT 4 gel documentation system. TAMRA fluorescence was collected under green LED illumination with UV filter (1 second exposure). Coomassie stain was imaged by white light transillumination. Densitometry analysis was performed in GelAnalyzer. The fold change in TAMRA intensity of TePhe-treated samples over vehicle (after normalization within each lane by Coomassie intensity) was calculated as a value enabling comparison between gels regardless of absolute intensities.

#### Fluorescence microscopy

Unless otherwise specified, images were acquired on a Mica Widefield Microhub automated microscope with spectral deconvolution capabilities (Leica), using 10x and 20x air objectives. Consistent illumination intensity was achieved using the “Relight” function. Confocal imaging was performed on a Leica TCS SP8 microscope, access was provided through the Cell & Systems Biology Imaging Facility at the University of Toronto. Samples were mounted with Type F immersion liquid and imaged using 20x and 40x oil objectives. Control of imaging parameters was achieved using LAS-X software (Leica). Post-acquisition processing was performed in ImageJ FIJI. All images were exported as PNG files.

### Imaging mass cytometry

Cell-ID Iridium DNA intercalator solution (201192B) was obtained from Standard BioTools. FPPE brain tissue sections for IMC analysis were deparaffinized and rehydrated as described for fluorescence imaging. Samples were equilibrated in PBS for 5 min, incubated in DNA intercalator solution (0.5 µM Cell-ID DNA Intercalator-Ir in PBS) for 30 min, then immersed in ultrapure water for 8 min. Sections were then left to fully air dry for subsequent IMC analysis. IMC data was acquired from 1 mm^2^ regions using a Standard Biotools Hyperion+ Imaging System.

Data analysis was conducted using MCDViewer, Rakaia, and ImageJ FIJI. To increase signal-to-noise of the natural abundance Te signal as described in literature^88^, a xenon background correction (Xe131) was applied to the most abundant tellurium isotope channels (Te126, Te128, Te130), which were then combined as a geometric mean to give the final Te distribution. Cell nuclei locations were identified by Ir distribution, taking the geometric mean of the two iridium isotopes (Ir191, Ir193). Max thresholds were set to 99% of max geometric mean signal for each element, while min thresholds were left at zero for better comparison with fluorescence images.

### Hematoxylin & eosin staining

Hematoxylin (HEM001.25) and Eosin Y disodium salt (EOS109.25) were purchased from BioShop Canada. Aluminium ammonium sulfate dodecahydrate (alum) was purchased from Ward’s Science (470300-118). Hematoxylin stain was formulated at 10 g/L hematoxylin, 50 g/L alum, and 0.4 g/L NaIO3 in 70:30:2 H2O/glycerol/acetic acid. Eosin stain was formulated at 2.5 g/L eosin in 80:20:1 ethanol/H2O/acetic acid. FFPE sections were deparaffinized and rehydrated as previously described. After rinsing in H2O for 5 min, sections were immersed in hematoxylin stain for 3 min, then rinsed twice in deionized water and briefly in 75% followed by 95% ethanol. Sections were immersed in eosin staining for 3 min, then washed four times in 95% ethanol. Sections were mounted in 80% glycerol. Images were acquired on a Mica Widefield Microhub automated microscope in brightfield mode. White correction was performed in FIJI.

### Cell culture

Mammalian cell lines were maintained at 37 °C in a humidified incubator with 5% CO2 atmosphere. Unless otherwise stated, high glucose DMEM (Sigma D5796) supplemented with 10% fetal bovine serum (Sigma F1051) and 1% penicillin-streptomycin (referred to as “DMEM + 10% FBS”) was used.

#### Toxicity assays

TePhe toxicity was determined using the alamarBlue viability assay (Invitrogen). 10000 N2a cells were seeded per 96-well (4 replicate wells per condition). 24 to 42 hr post-seeding, TePhe treatment was initiated with media exchange to fresh DMEM + 10% FBS containing stated concentrations of TePhe such that all groups reached endpoint simultaneously. After treatment, media was replaced with fresh DMEM + 10% FBS and cells allowed to grow for 40 hr. Media was then replaced with 1:10 alamarBlue reagent in fresh DMEM + 10% FBS and cells incubated for 1.5-2 hr, followed by measurement of resorufin absorbance on a Biotek Synergy H1 microplate reader.

#### TePhe treatment for proteomics

HEK293T cells were plated at 1.6 × 10^6^ cells per 10-cm dish and grown 2 days, at which point the media was replaced with fresh DMEM + 10% FBS containing 400 µM TePhe (or unmodified DMEM + 10% FBS for control) for 4 hr. N2a cells were plated at 4 × 10^6^ and 3 × 10^6^ cells per 10-cm dish for undifferentiated and differentiation-destined samples, respectively. After 24 hr, the undifferentiated group was treated with DMEM + 10% FBS containing 400 µM TePhe for 8 hr, while the differentiation group was exchanged into DMEM + 2% FBS + 20 µM retinoic acid and incubated for a total of 88 hr (media replenished halfway through the differentiation period), followed by TePhe treatment in DMEM + 2% FBS + 20 µM retinoic acid + 400 µM TePhe for 8 hr. Upon completion of TePhe treatment, cells were washed with PBS and lysed in PBS + 0.5% SDS + cOmplete Mini protease inhibitor cocktail. Lysate was transferred to a 2 mL microcentrifuge tube, homogenized by drawing through a 22-gauge hypodermic needle, and stored at -80 °C.

### Mass spectrometry-based proteomics

#### General procedures

All samples were subjected to a pre-treatment consisting of reduction, alkylation, and acetone precipitation. Lysates were diluted to a final concentration of 0.5 to 1.0 mg/mL protein in 100 mM Tris + 100 mM NaCl, pH 8.5 + 0.5% SDS. Disulfides were reduced by the addition of DTT (7-10 mM) for 45 min at 55 °C, cooled to RT, and alkylated with chloroacetamide (15 mM) for 25 min in the dark. Reactions were quenched with additional DTT (5-7 mM) for 5 min, followed by acetone precipitation (as described under “In-solution fluorescent OSTAC labeling of tissue lysates”). Precipitates were redissolved, quantified, and normalized before proceeding. Sequencing grade trypsin was purchased from Promega (V5113). Overnight tryptic digests were carried out in 100 mM ammonium bicarbonate (pH ∼8), at 37 °C with vigorous shaking on an Eppendorf Thermomixer C platform. Digests were quenched by acidification with 1% formic acid. Peptide clean-up was performed on C18 tips (Pierce #87784) according to the manufacturer’s instructions (with substitution of trifluoroacetic acid by formic acid). Eluted peptides (in 70% acetonitrile + 0.1% formic acid in H2O) were dried *in vacuo* and redissolved in 0.1% formic acid in H2O. Peptide concentrations were determined by micro-scale BCA assay (using 1 µL sample/standard + 4 µL working reagent and quantification by on a NanoDrop spectrophotometer at 562 nm). Solutions were stored at -80 °C until LC-MS/MS injection.

#### Preparation of bulk brain proteome

2 mm coronal brain slices were roughly crushed with a pipet tip, suspended in PBS + 1% SDS (2.5 mL), and homogenized by repeatedly drawing contents through a 22-gauge hypodermic needle. Lysates were briefly sonicated (∼10 sec), filtered through a 35 µm mesh strainer, and stored at -80 °C until further processing. Quantified lysates (450 µg input) were subjected to reduction, alkylation, and acetone precipitation. Pellets were redissolved in 3 M urea in 100 mM ammonium bicarbonate (130 µL), diluted with 100 mM ammonium bicarbonate to 390 µL, and pelleted at 10000 × *g* for 1 min to remove any undissolved particles. Cleared supernatants were quantified and further diluted to 0.5 mg/mL protein in 100 mM ammonium bicarbonate, from which 150 µL were taken for tryptic digest. CaCl2 (1 mM) and MS-grade trypsin (1.9 µg) were added and samples incubated overnight at 37 °C, 600 rpm. Peptide clean-up was performed as described above.

#### OSTAC-based biotinylation, streptavidin capture, and on-bead digest

Cell/tissue lysates were subjected to reduction, alkylation, and acetone precipitation as described above. Redissolved proteins (1.0 mg/mL in PBS + 0.5% SDS) were OSTAC labeled by addition of DBCO (150 µM) + NCS (550 µM) in rapid succession and incubating for exactly 10 min, followed by addition of BCN-BtnSO2 (10 µM) + NCS (100 µM) and incubating again for exactly 10 min. Reactions were quenched by addition of DTT (3-5 mM). A second round of acetone precipitation was performed to remove excess biotin probe. Pellets were reconstituted in PBS + 1% SDS, diluted to 0.2-0.25 mg/mL, and applied to magnetic streptavidin beads (Genscript L00936) or streptavidin agarose resin (Genscript L00353) to enrich biotinylated proteins. More detailed bead equilibration and washing procedures can be found in the Supplementary Information, Section 4. Briefly, samples were diluted to 0.25 mg/mL protein in PBS + 1% SDS and incubated with equilibrated beads for 1.5 hr at RT on an over-the-end rotator. Beads were washed with 4 × PBS + 1% SDS, 2 × PBS, and 2 × 100 mM ammonium bicarbonate. Overnight on-bead tryptic digest was performed at an enzyme/input ratio of 1:100-1:150, at 37 °C, 1200 rpm, followed by peptide clean-up and quantification.

#### LC-MS/MS data acquisition and analysis

Peptides were separated using an Easy n-LC 1200 liquid chromatography system (Thermo Fisher) connected to a 14 cm PicoFrit column with 75 µm internal diameter (New Objective) packed in-house with 1.9 µm ReproSil-Pur C18-AQ (Dr. Maisch). A gradient of 2 to 80% acetonitrile (0.1% formic acid in H2O) over 105 min followed by isocratic elution at 80% acetonitrile for 15 min, at a constant flowrate of 300 nL/min and total duration of 120 min, was applied to all samples. For bulk brain proteomics, 3.2 µL (750 ng) sample was used per LC-MS/MS injection. To compare abundances of affinity captured peptides between vehicle and TePhe-treated HEK293T samples, all samples were reconstituted to the same volume and 20% of total peptide output used for injection (4.0 µL; on average 240 ng for vehicle samples, 780 ng for TePhe-treated). For TePhe-treated N2a proteomes at varying differentiation stages, peptide concentrations were normalized to achieve consistent loading of 800 ng in 4.0 µL. To compare abundances of affinity captured peptides between vehicle and TePhe-treated (control and mutant) forebrains, all samples were reconstituted to the same volume and 70% of total peptide output injected (5.5 µL; on average 600 ng for vehicle, 800 ng for TePhe-treated).

Protein identification was performed in MaxQuant v.2.6.6.0^89,90^ via the Trillium HPC. Spectra were searched against their respective human and mouse reference proteomes (Uniprot UP000005640 and UP000000589 respectively, downloaded July 2026). For all searches, trypsin/P was selected as the digest condition with up to 2 missed cleavages allowed. Minimum peptide length was set at 7, maximum peptide mass at 4600 Da, and mass tolerance at 4.5 ppm for peptide ions and 20 ppm for fragment ions. Peptide spectrum match and protein false discovery rates were both set at 0.01. Carbamidomethyl-cysteine was set as a fixed modification while N-terminal acetylation and methionine oxidation were set as variable modifications for all searches. For OSTAC-treated samples, tryptophan side chain oxidation (+15.994914 Da) was included as a variable modification. Match-between-runs was disabled while label-free quantification (minimum of 1 ratio count for bulk brain proteome analysis, 2 ratio counts for all other experiments) was enabled for all searches.

Data processing and visualization were performed in Perseus.^91^ After removal of potential contaminants, reverse peptide sequences and those identified by site only, intensities (raw or LFQ) were converted to log2 values. Proteins appearing in at least 2 replicates of at least one condition of a particular experiment were retained, after which missing values were supplied by imputation. Differential protein expression/pull-down was assessed by Student’s t-test.

### Gene ontology analysis

Over-representation testing was performed with PANTHER.^92–94^ From RNA-seq data, all genes meeting the *padj* <0.05 cutoff for differential expression were used as input. For bulk brain proteomics, protein groups meeting both the 1.5-fold change and *p* <0.05 thresholds were used. Each was tested against the GO: Biological Process, GO: Cellular Component, and GO: Molecular Function complete annotation sets. All genes/protein groups detected in their respective experiments were used as the reference list.

Pathway analysis for protein synthesis differences between undifferentiated and differentiated N2a cells was visualized using ShinyGO 0.85.2.^95^ The list of all protein groups which could be quantified by t-testing of LFQ intensities was used as background. Down- and up-regulated proteins were tested separately. For entries where peptides were mapped to multiple proteins IDs, only the first ID generated by MaxQuant was used.

#### Analysis of Sequence Length and Phe content

Sequences were retrieved using the UniProt ID Mapping function (UniProtKB accession number as input). Where multiple IDs were associated with a particular protein group, only the first ID was used. Sequence length, number of Phe residues, and Phe content (%) were calculated in Excel and correlated with corresponding TePhe- to-vehicle fold enrichment values observed by LC-MS/MS. For proteins detected across HEK293T lysates, 4187 of 4190 entries were successfully mapped to reviewed (Swiss-Prot) sequences. For brain transcriptomic data, all entries were searched, with 15999 mapping to coding sequences with calculable Phe content. This distribution was compared against the Phe content distribution of all proteins identified by LC-MS/MS.

### Statistical testing

Statistical testing on TePhe dosing regimes and clinical chemistry parameters was performed in GraphPad Prism. TePhe dosing regimes in Fig. 2b were compared by ordinary one-way ANOVA (replicates corresponded to individual animals, *n =* 3 or 4 for each dosing regime, as indicated). Serum clinical chemistry parameters were compared by unpaired t-test with Welch’s correction (replicates corresponded to individual animals, *n* = 4 for vehicle, *n* = 3 for TePhe-treated). Differential expression analysis for RNA-seq was performed using DESeq2. T-tests on proteomics data were performed in Perseus. After log2 transformation of intensity values, protein groups were appearing in at least 2 replicates in any group were retained and subjected to imputation followed by two-sided Student’s t-test.

## Acknowledgments

We thank the Bioscience Support Facility for mouse housing, husbandry, support, and training, the Cell and Systems Biology Imaging core for access to confocal microscopy and Imaris, and the Centre for Applied Genomics for next generation sequencing. We also thank the Transgenic Core facility at the Salk Institute for Biological Studies for generating the original RARS1 loxP/loxP insertion as well as Dr. Paul Schimmel for support in the generation of the mouse line. We appreciate insightful discussions on TePhe toxicity with Prof. Mathieu Lemaire at The Hospital of Sick Children, and preliminary experiments by Dr. Shahbaz Khan and Prof. Thomas Kislinger at the Princess Margaret Cancer Centre.

HC acknowledges funding by the Natural Sciences and Engineering Research Council of Canada (NSERC, RGPIN-2023-04305), the Canadian Institutes of Health Research (CIHR, PJT 497271), the Canada Foundation for Innovation/John R. Evans Leaders Fund and the Ontario Research Fund for instrument support, as well as the Department of Chemistry and the Faculty of Arts and Science at the University of Toronto. YJB was supported by a CIHR postdoctoral fellowship, SPN by a CIHR Canada Graduate Scholarships – Master’s and Doctoral program. This research was enabled in part by support provided by Scinet, the Niagara and Trillium Cluster at the University of Toronto, and the Digital Research Alliance of Canada (alliancecan.ca).

Individual icons for figures for sourced from Biorender.

## Competing interests

MN holds a patent describing TePhe which has been licensed to Standard Biotools.

