## Supplementary Figures for "A Metabolic Labeling Strategy for Tracking Protein Synthesis in Complex Biological Systems"

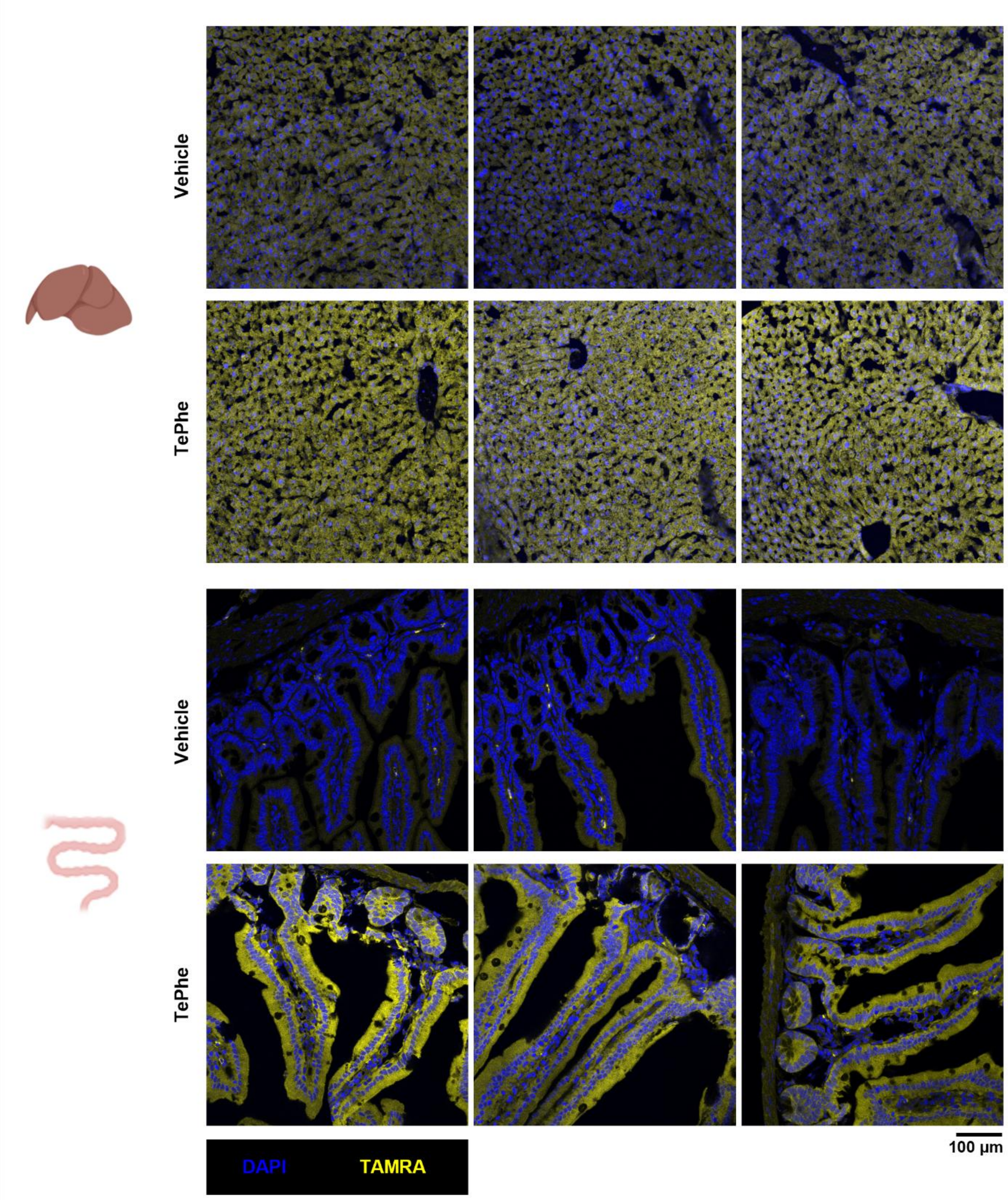


**Figure S1. Fluorescent OSTAC labeling in liver and intestine cryosections.** Confocal imaging revealed a clear signal increase in the TAMRA channel in tissues TePhe-treated tissues over vehicle. Section thickness = 8 µm.

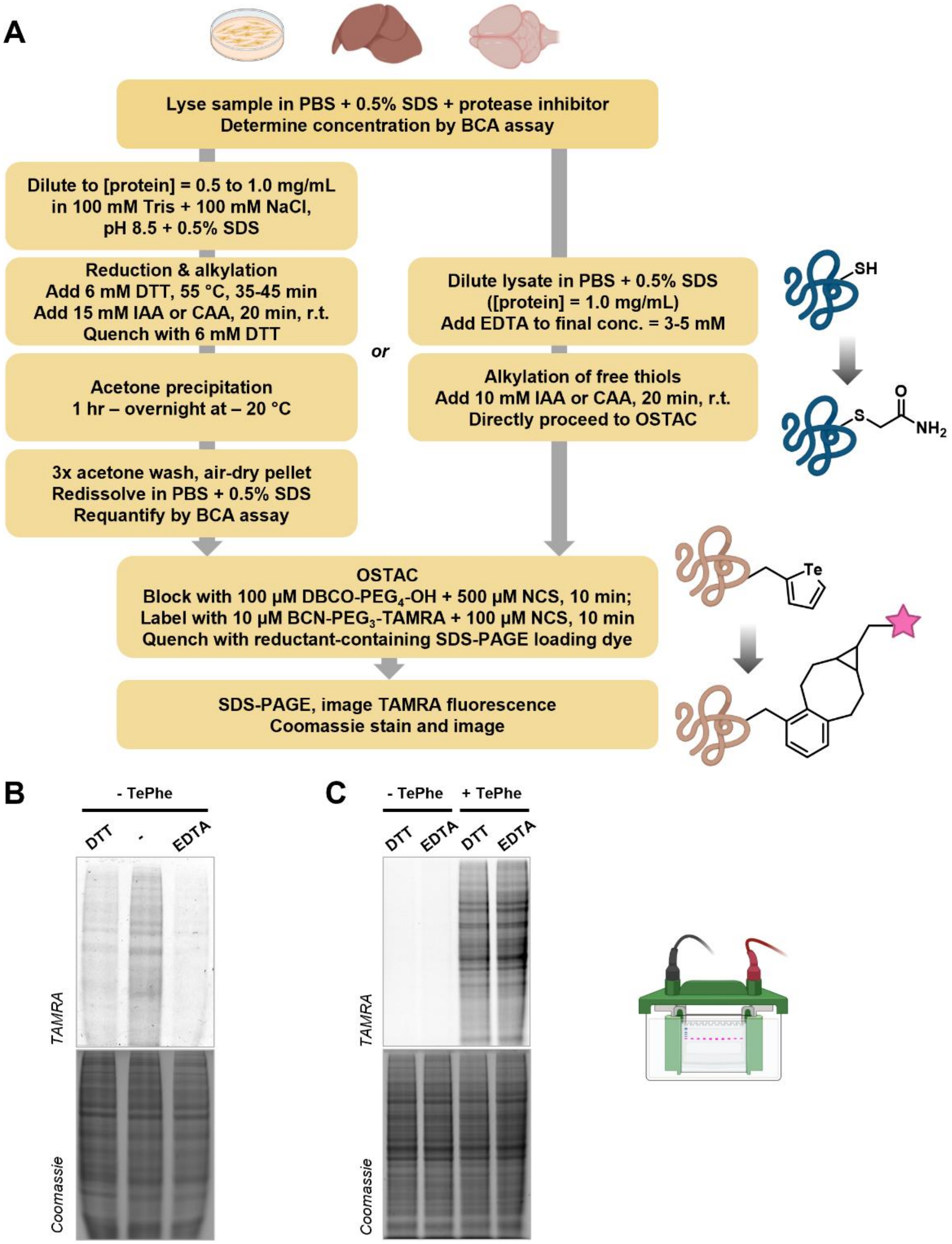


**Figure S2. In-solution fluorescent OSTAC labeling for lysates.** A) Schematic of labeling workflow. To prevent free thiols from interfering with OSTAC chemistry, iodoacetamide (IAA) alkylation was performed. Lysates could be reduced with DTT followed by IAA alkylation and acetone precipitation before OSTAC. It was later discovered that lysates could be alkylated with IAA in the presence of EDTA and subjected directly to OSTAC; EDTA mitigates OSTAC background to a similar degree as DTT pre-treatment while shortening the overall workflow. B) SDS-PAGE showing background reduction by DTT or EDTA pre-treatment method in HEK293T lysate, compared to direct OSTAC labeling. C) SDS-PAGE confirming comparable signal to noise using either pre-treatment strategy prior to OSTAC (HEK293T lysates, untreated vs. 400 μM TePhe, 4 hr).

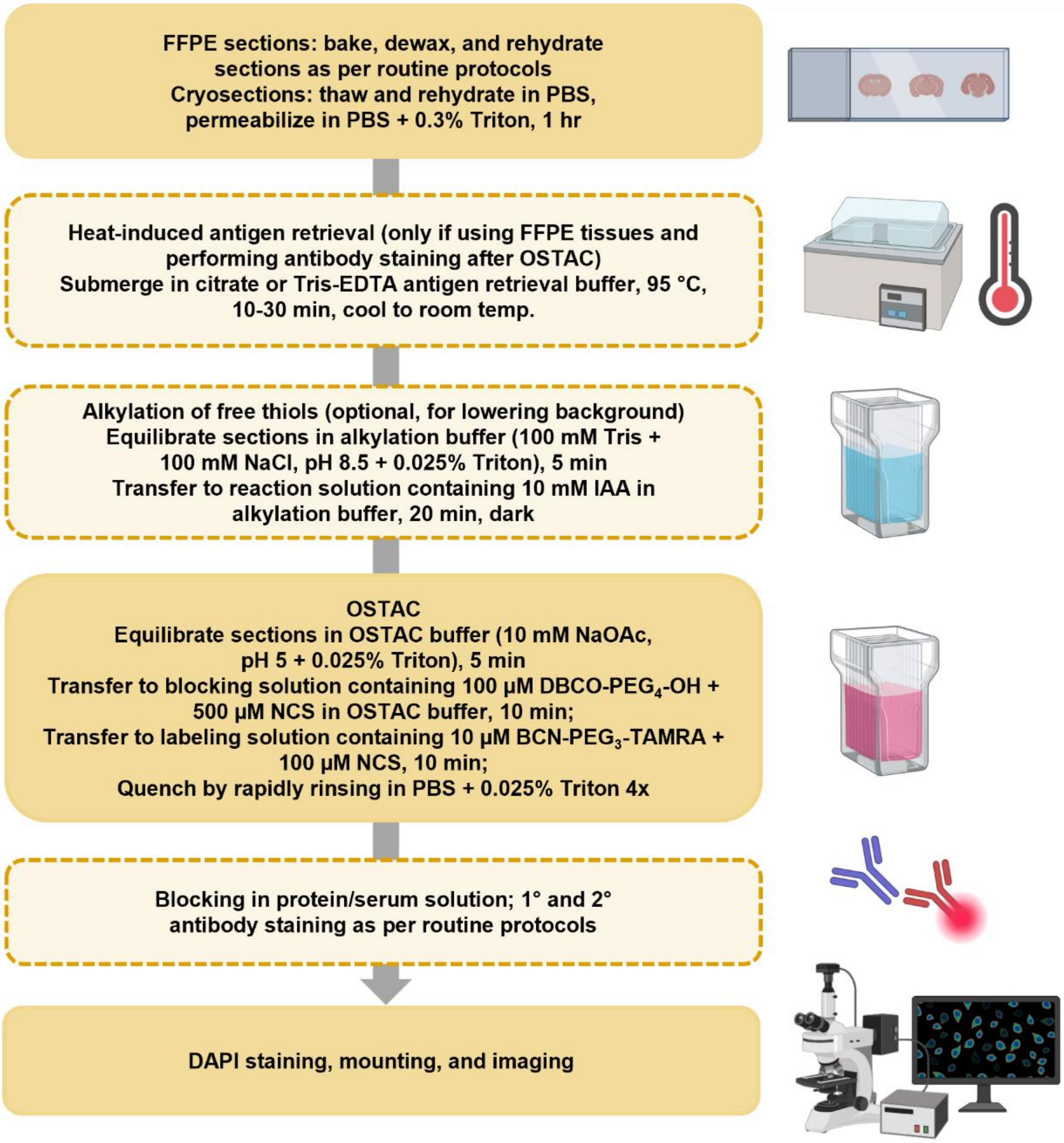


**Figure S3. Schematic of fluorescent OSTAC labeling for slide-mounted tissue sections.** Cryo- and FFPE tissues were pre-processed per routine protocols. For FFPE tissues, heat-induced antigen retrieval could be performed prior to OSTAC labeling if downstream immunostaining was desired; otherwise OSTAC chemistry could be directly performed on rehydrated tissues. Alkylation with IAA prior to OSTAC aided background reduction but could be omitted. Once the OSTAC labeling sequence was completed, tissues could be directly counterstained with DAPI and mounted for imaging, or taken to procedures for routine blocking and antibody binding.

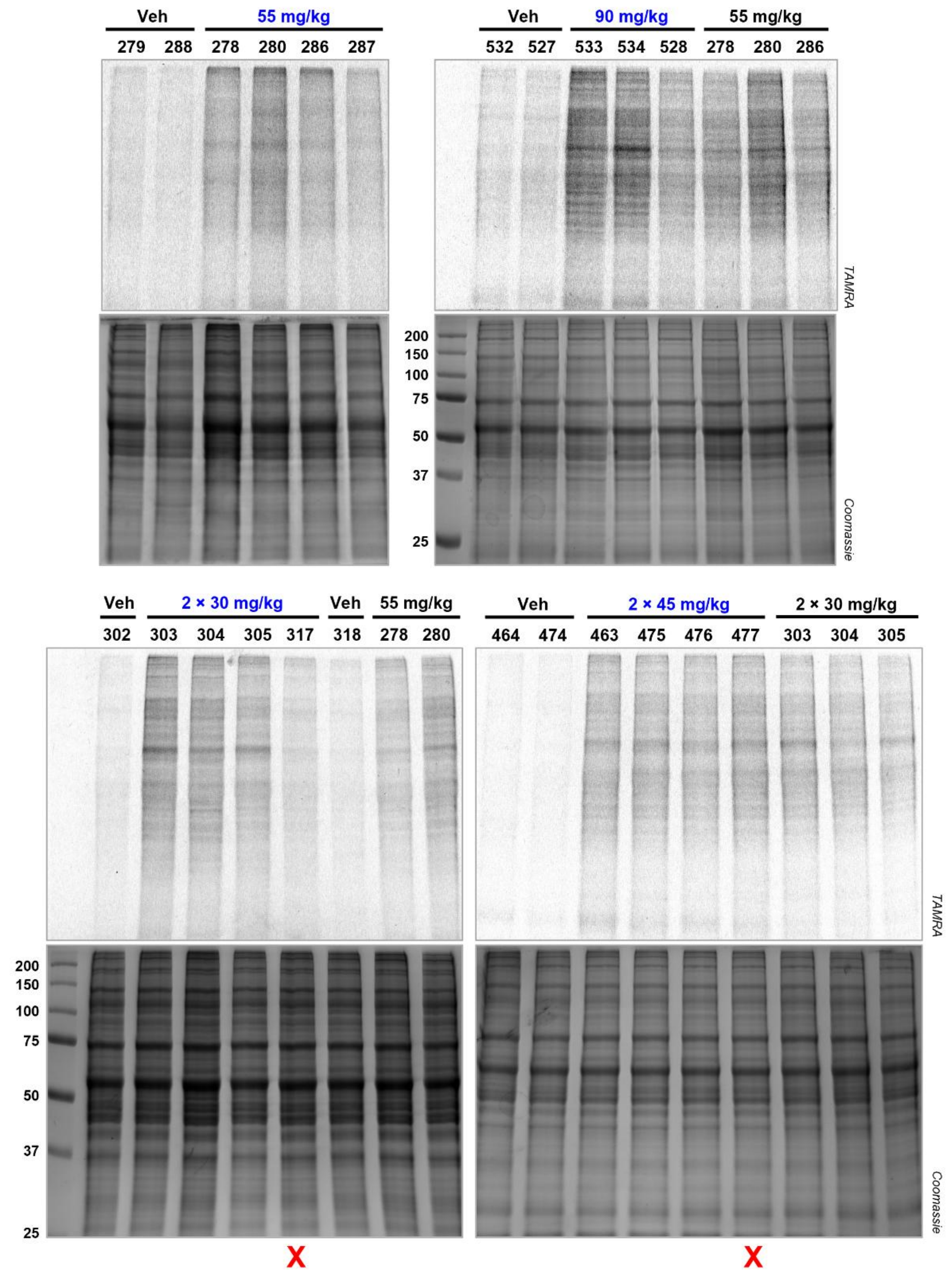


**Figure S4. In-gel fluorescence quantification of TePhe incorporation within the brainstem.** Each lane represents one animal. OSTAC signal (TAMRA fluorescence) was quantified by densitometry and normalized to Coomassie intensity. Normalized signals of TePhe-treated samples were divided by the average vehicle background to determine TePhe/Vehicle ratios plotted in Fig. 2b. The dosing regime highlighted in blue is the condition being quantified in each gel. After the initial cohort (55 mg/kg TePhe), gels were loaded with samples from an earlier cohort to ensure comparableness of TePhe/Vehicle ratios across gels from different days. Lanes marked with a red "X" are excluded from the final analysis due to unsuccessful injection or incorrect genotype.

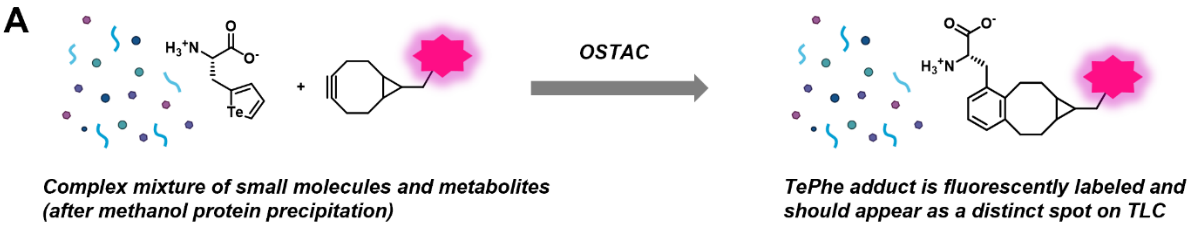


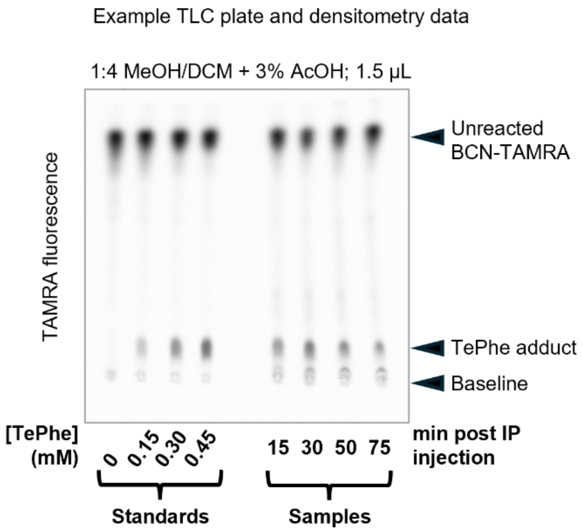


| Standard (µM TePhe) | Adduct Integration | Sample (min post injection) | Adduct Integration |
|---|---|---|---|
| 0 | 137 | 15 | 2229 |
| 150 | 1175 | 30 | 2741 |
| 300 | 2596 | 50 | 1971 |
| 450 | 3302 | 75 | 1474 |

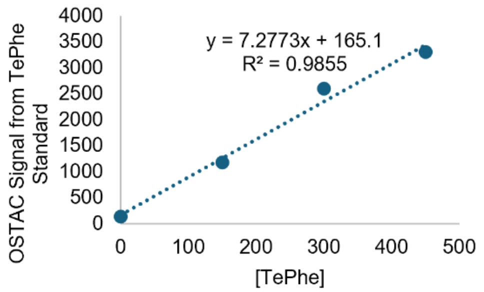


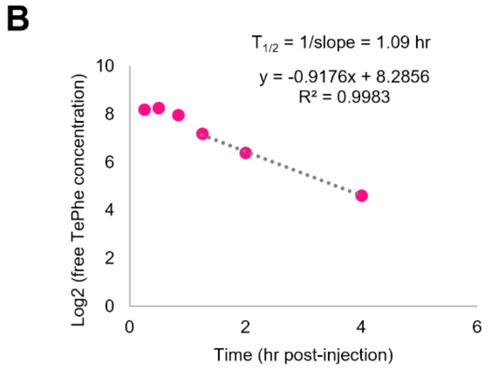


**Figure S5. Fluorescent OSTAC-based TLC assay for detection of free TePhe in biofluids.** A) Free TePhe is visualized by formation of a fluorescent OSTAC adduct with BCN-TAMRA. The example TLC shows quantitation of free TePhe in serum of mice injected with 90 mg/kg TePhe and sacrificed at indicated timepoints, using a standard curve of untreated mouse serum that has been spiked with known concentrations of TePhe. B) Semi-log plot of serum TePhe concentration data shown in Fig. 2c. Later timepoints, at which contributions from delayed absorption after IP injection are expected to be minimal, showed a linear decline and were used to estimate the half-life of TePhe in mouse serum.

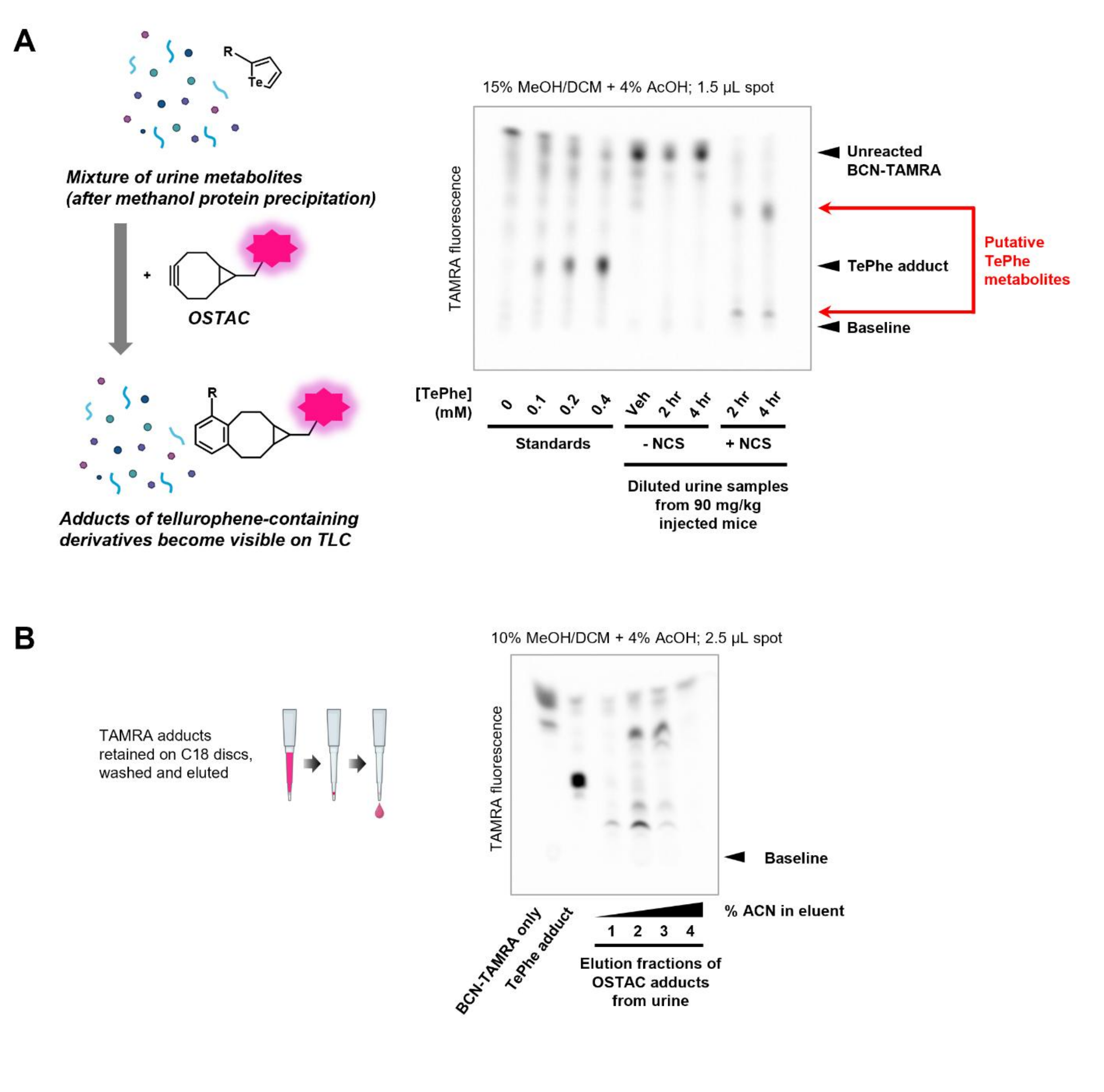


**Figure S6. OSTAC-based fluorescent TLC analysis of TePhe metabolites in mouse urine.** A) TLC analysis of urine metabolites after fluorescent OSTAC labeling shows essentially no free TePhe in urine. Two new spots were found to be present only when oxidant is added to the OSTAC labeling mixture, consistent with the behaviour of intact tellurophenes ("putative TePhe metabolites"). B) OSTAC adducts of urine metabolites with BCN-TAMRA could be concentrated using C18 discs. Elution with increasing fraction of acetonitrile in water yielded two fractions (2 and 3) enriched in putative TePhe metabolites, which were further analyzed by mass spectrometry.

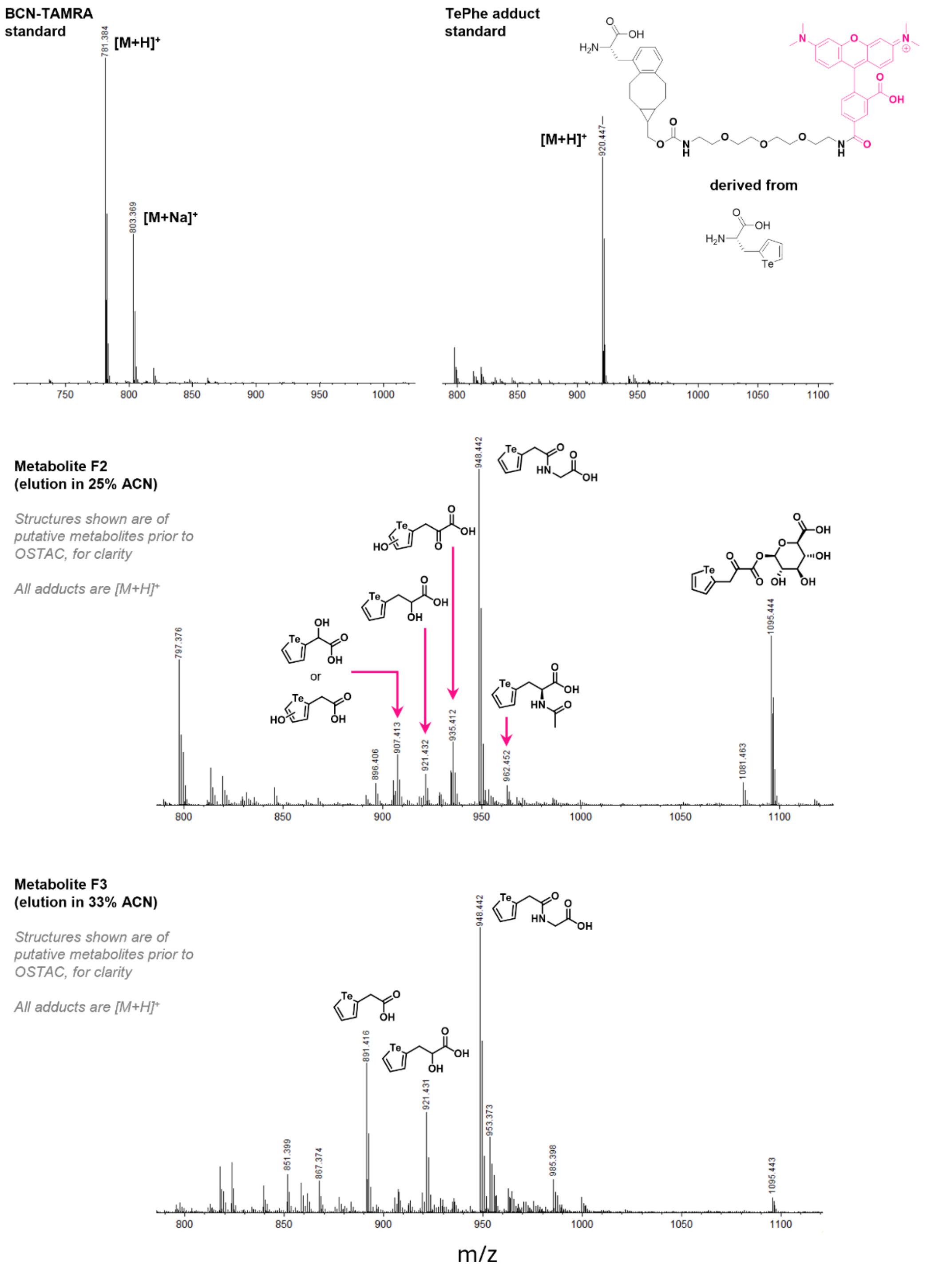


**Figure S7. MS analysis of OSTAC adducts found in urine of TePhe-treated mice.** Structures overlaid on the spectra of metabolite fractions F2 and F3 are hypothesized starting structures whose expected OSTAC adducts with BCN-TAMRA have masses matching the observed signals to within 10 ppm.

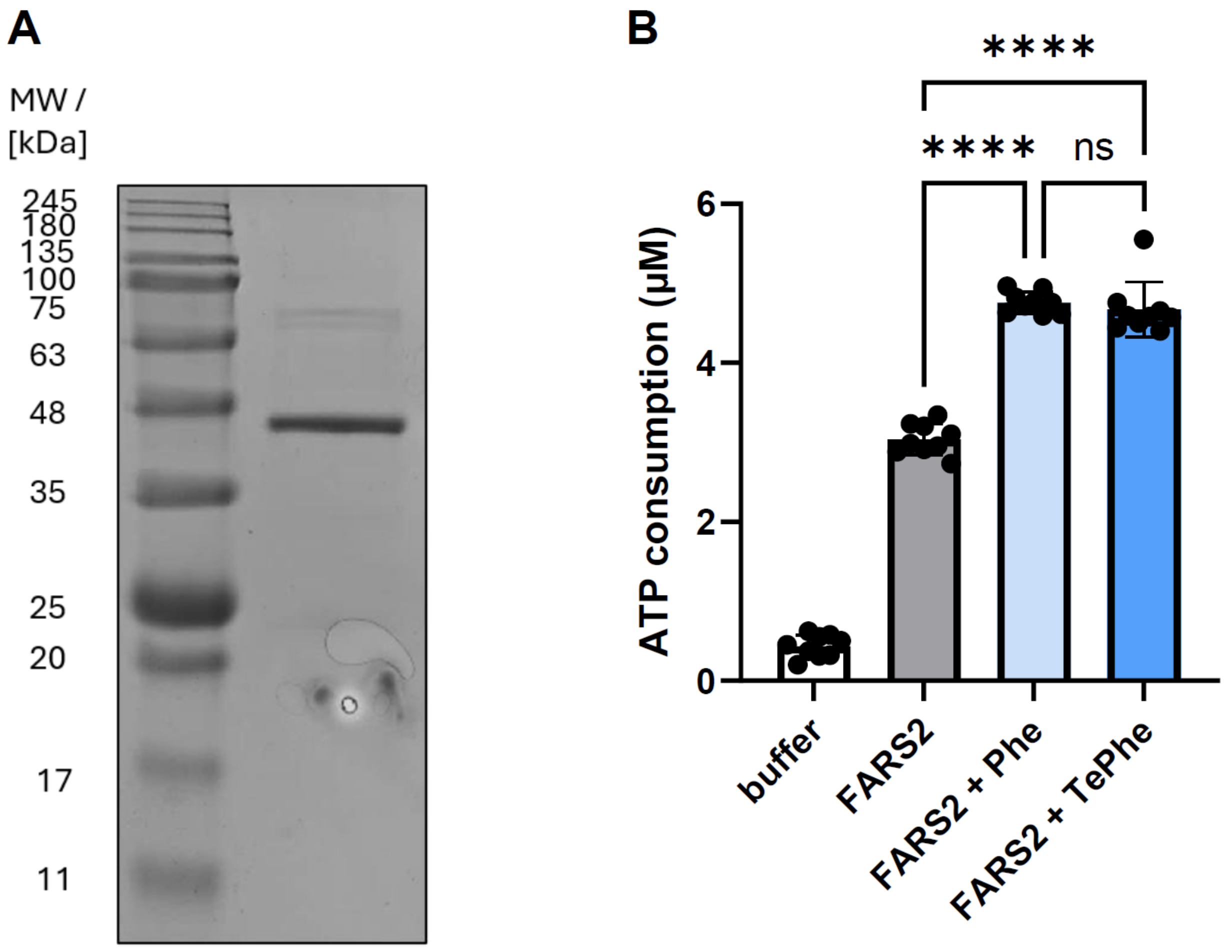


**Figure S8. Assay quantifying ability of mitochondrial phenylalanyl-tRNA synthetase to activate TePhe compared to Phe.** A) SDS PAGE confirmation of FARS2 purity following recombinant protein expression in *E. coli* and multistep purification. FARS2 was used over the cytosolic Phenylalanyl-tRNA synthetase, as the latter is a dimer of FARSA/B dimers and challenging to purify. B) A luciferase-based assay was used to measure ATP consumption as a means of quantifying the first step of the aminoacylation reaction, the activation of the amino acid to aminoacyl-AMP. Each datapoint represents one replicate in a 96-well plate, measured over two experiments ($n$ = 9, 9; one-way ANOVA, **** $p$ <0.0001).

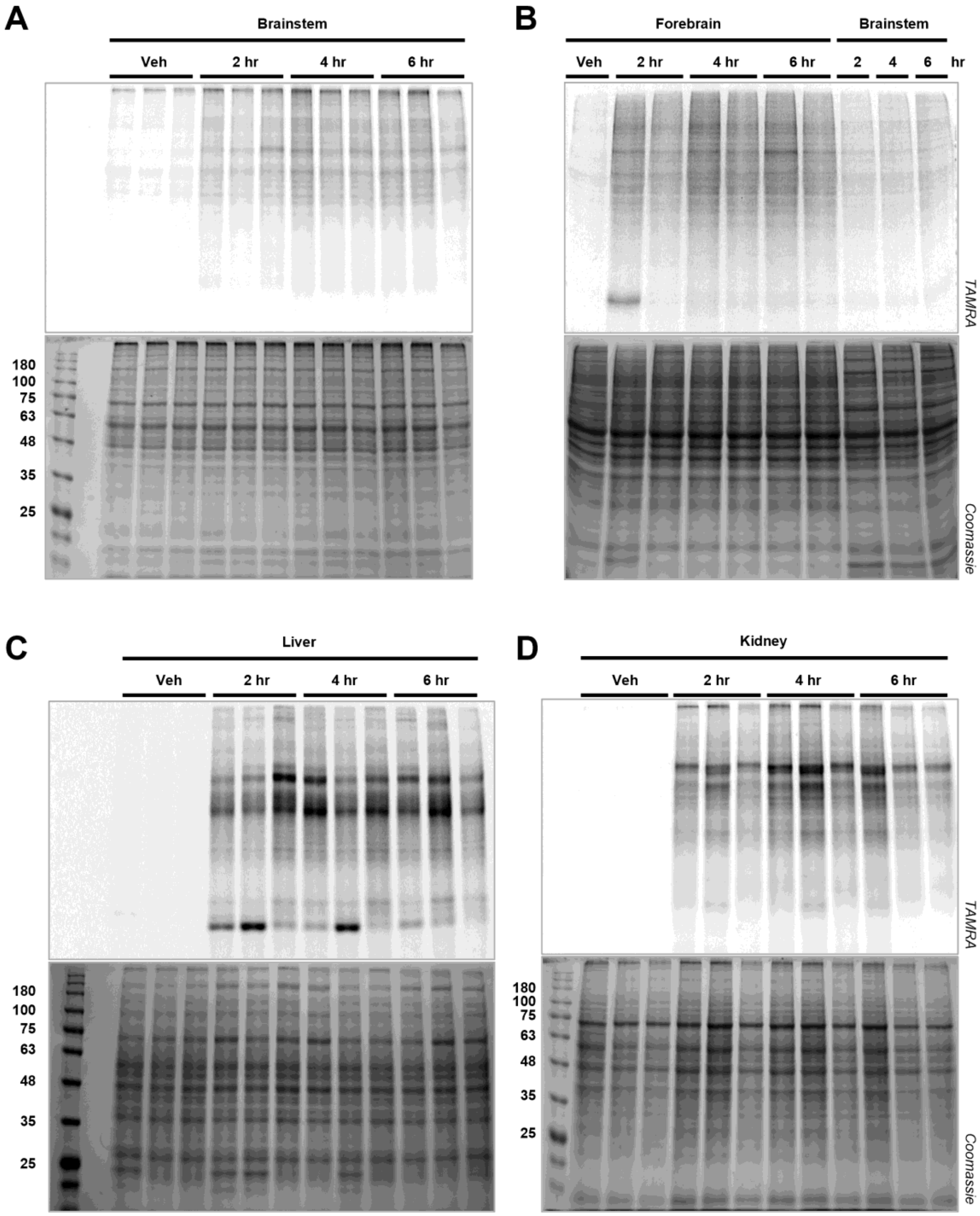


**Figure S9 (cont'd on next page)**

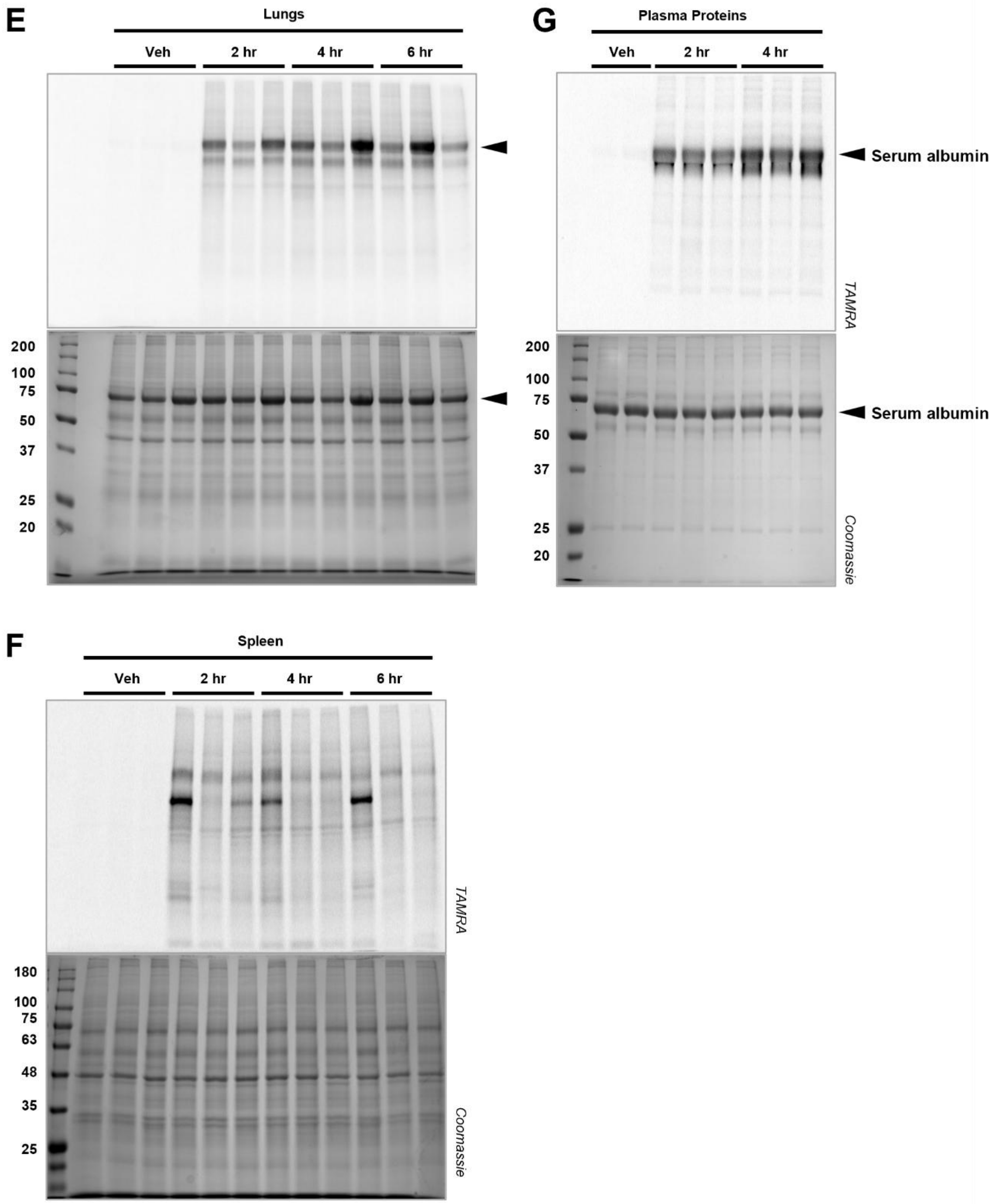


**Figure S9. In-gel fluorescent detection of TePhe incorporation in major organs.** Time course showing incorporation after a single 90 mg/kg IP injection in: A) brainstem, B) forebrain, as compared to brainstem, C) liver, D), kidney, E) lungs, F) spleen, G) plasma proteins. Note that major band observed in lung lysate is likely serum albumin due to variable entrapment of blood within lung tissue (indicated by arrows).

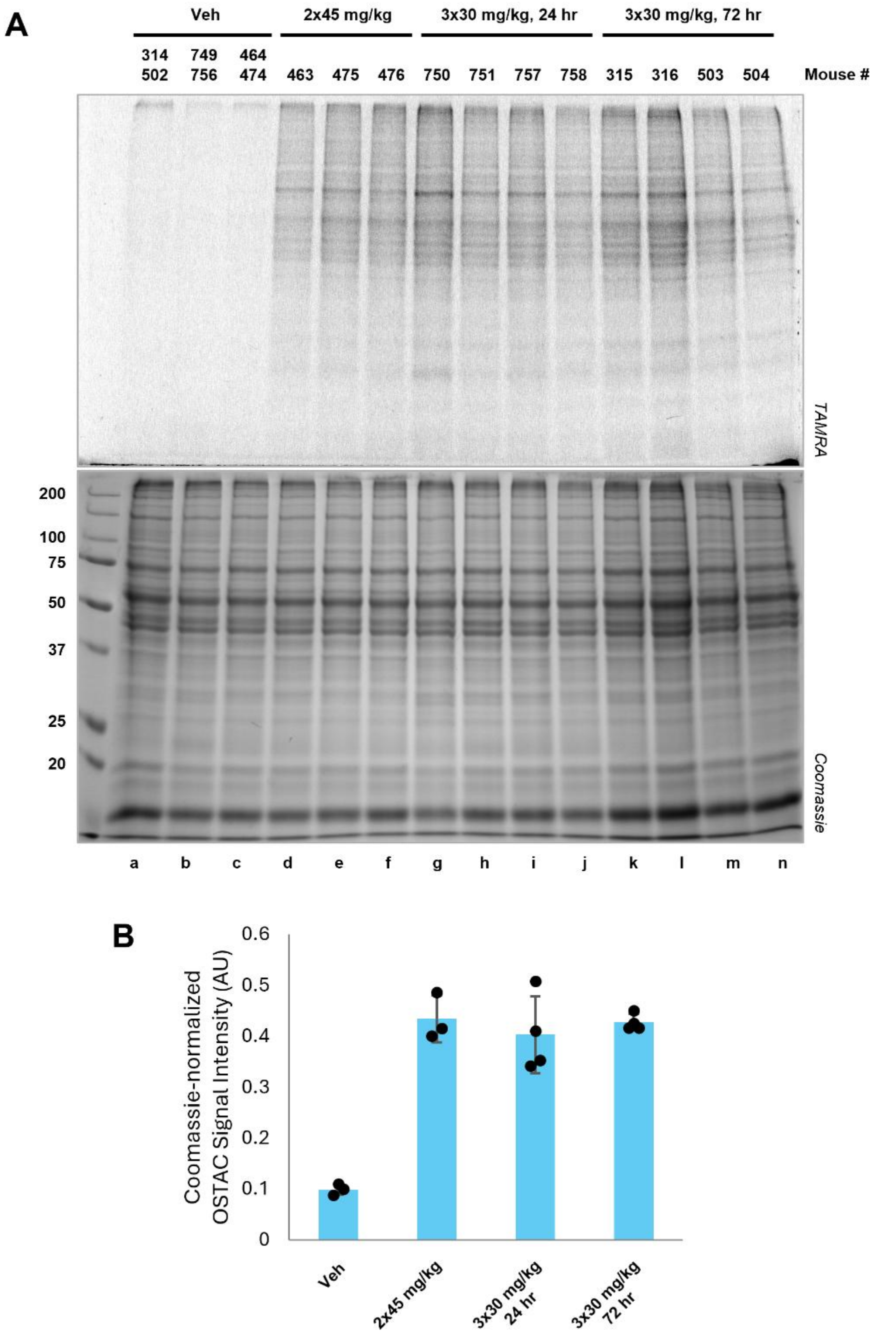


**Figure S10. In-gel fluorescent detection of TePhe incorporation in brainstem after TePhe injections spaced over long intervals.** A) Gels showing OSTAC detection of TePhe (TAMRA fluorescence) and staining for total protein content (Coomassie). Lanes a-c comprise a 1:1 mixture of lysates from 2 vehicle animals, each the corresponding controls within a TePhe-treated cohort (a corresponding to k-n, b to g-j, c to d-f). As expected, vehicle lysates across cohorts exhibit similar background labeling. Lanes d-n represent lysates from one TePhe-treated animal each. B) Fold change in Coomassie-normalized OSTAC labeling intensity in TePhe-treated groups.

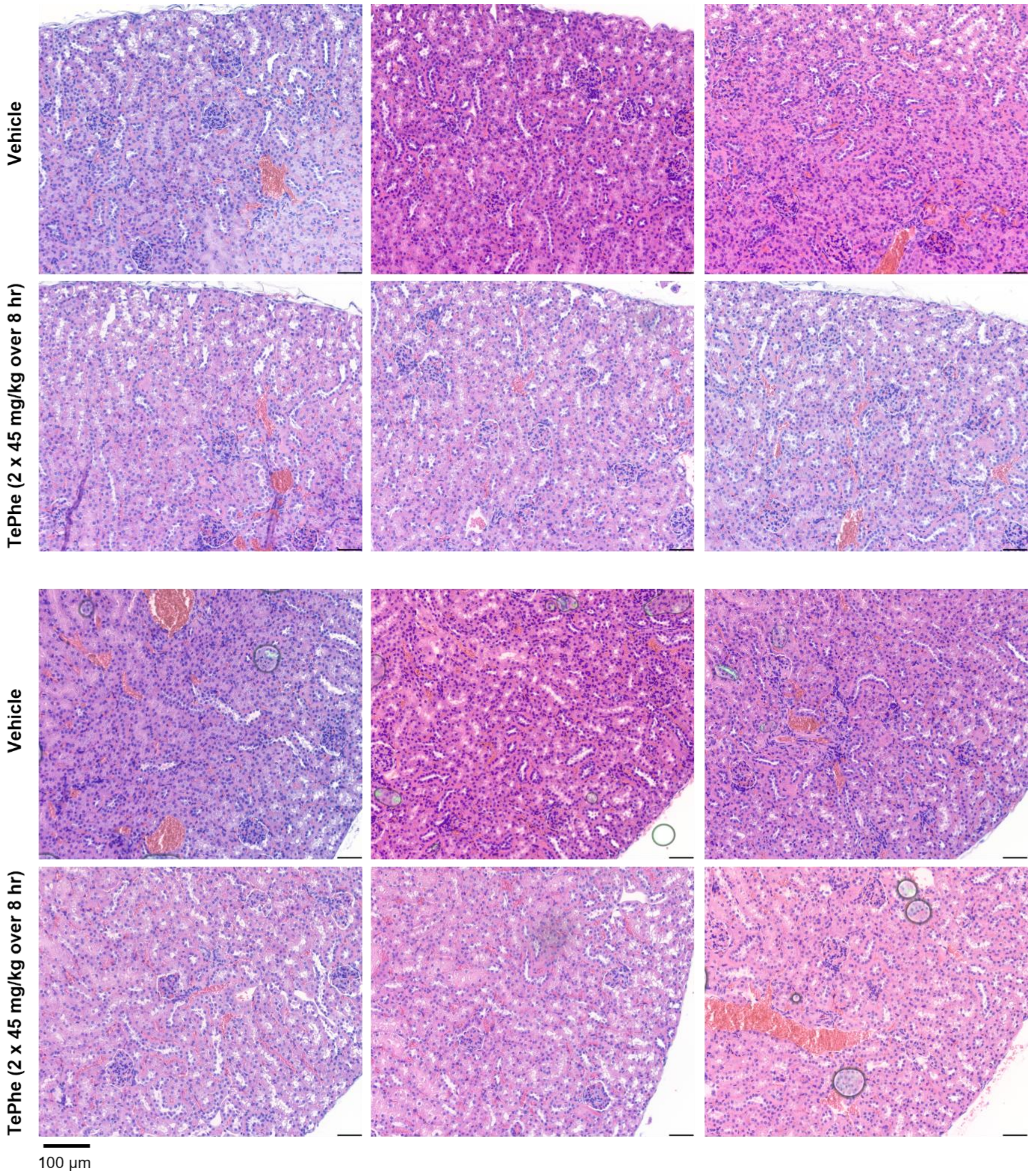


**Figure S11. Hematoxylin and eosin staining of renal cortical tissues.** No obvious morphological differences were observed between vehicle and TePhe-treated kidneys. Overall shape and spacing within glomeruli and renal tubules appear unchanged.

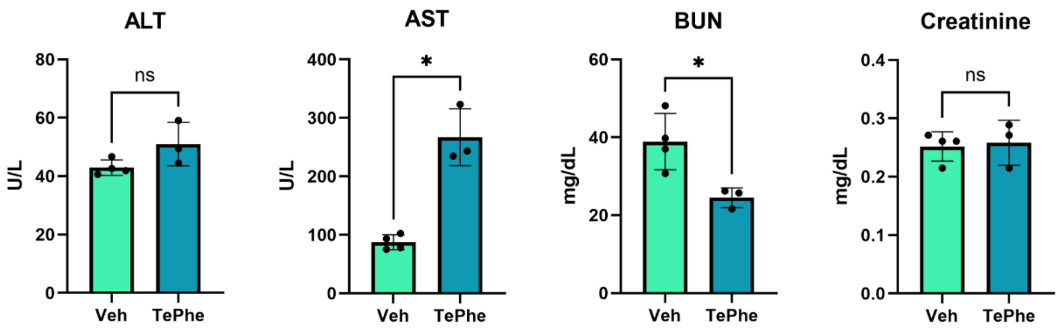


**Figure S12. Serum clinical chemistry parameters of vehicle and TePhe-treated (90 mg/kg IP injection, 6 hr) mice.** ALT = alanine aminotransferase. AST = aspartate aminotransferase. BUN = blood urea nitrogen. Serum from the same animals were used for all tests (*n* = 3 for vehicle, *n* = 4 for TePhe-treated). * $p < 0.05$, unpaired t-test with Welch's correction.

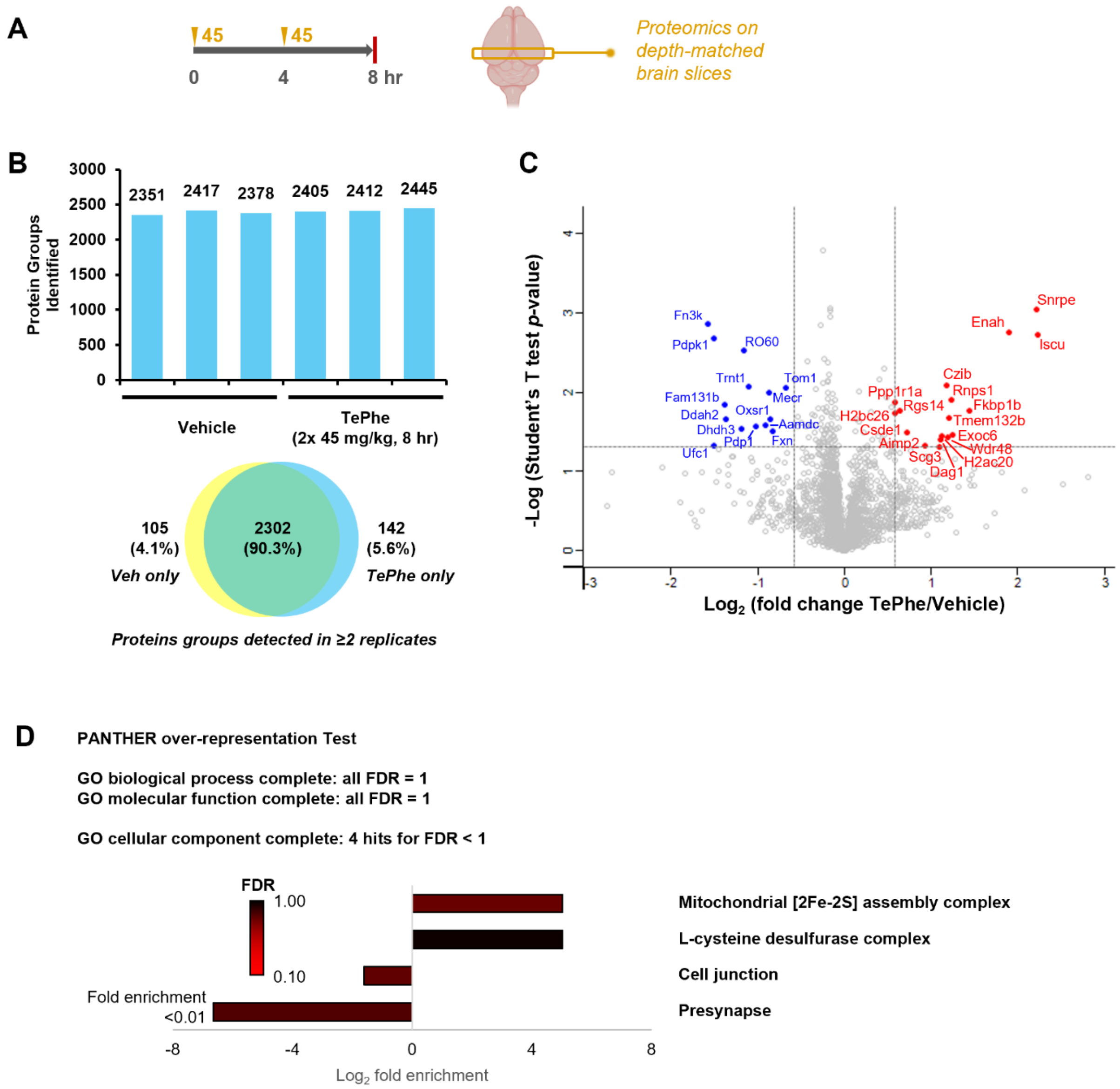


**Figure S13. Evaluation of TePhe toxicity within the mouse brain by shotgun proteomics.** A) Scheme showing TePhe dosing regime and sample collection. Mice were dosed with 2 × 45 mg/kg TePhe (or vehicle) by IP injections at 4 hr intervals and sacrificed after a total of 8 hr of TePhe exposure. Matching 2 mm slices of the midbrain were used for proteomics. B) Number and distribution of protein groups detected across samples. A total of 2738 protein groups were detected across all samples, with similar numbers detected across conditions and biological replicates. 2302 (90.3%) of protein groups appeared in at least 2 of 3 replicates in both vehicle and TePhe-treated groups. C) Volcano plot showing Student's t-test results on LFQ comparison of protein groups across vehicle and TePhe-treated tissues. Of 2474 proteins quantified, only 14 were down-regulated and 17 up-regulated in TePhe-treated samples (fold change ≥ 1.5, $p < 0.05$). D) Results of over-representation testing on combined list of all down- and up-regulated proteins. No hits were obtained when testing against the GO biological process and GO molecular function annotation sets. Low confidence hits (FDR between 0.1 and 1.0) were obtained when testing against the GO cellular component annotation set.

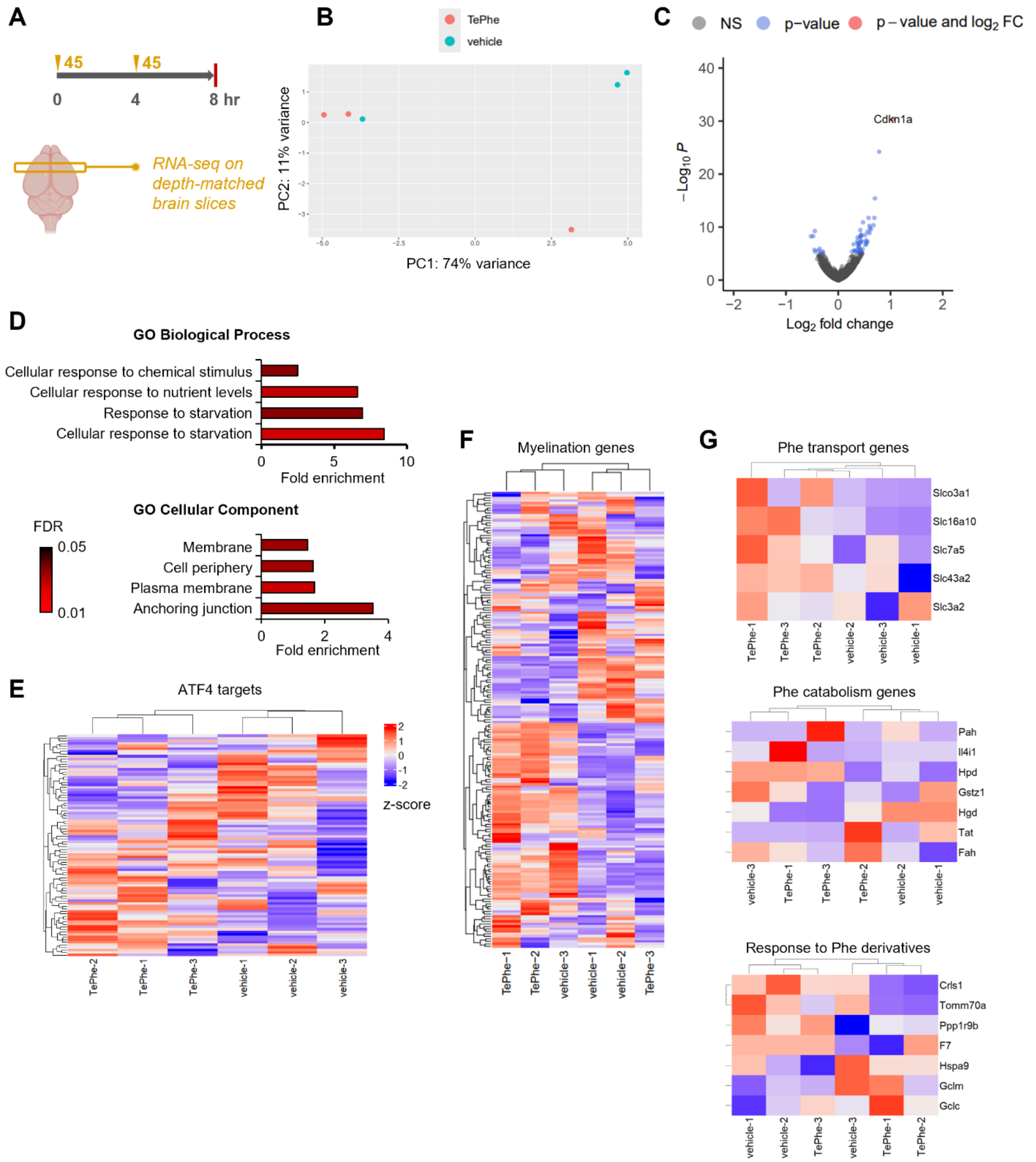


**Figure S14. Evaluation of TePhe toxicity within the mouse brain by transcriptomic analysis.** A) Volcano plot comparing TePhe-treated and vehicle proteomes. Protein groups exceeding |fold change| $\geqslant 2$ and $p < 0.05$ were considered hits. Only one significantly up-regulated gene, Cdkn1a, was identified. B) PCA analysis of vehicle and TePhe-treated transcriptomes show no clear clustering by treatment status. C) List of GO: Biological Process and GO: Cellular Component terms enriched upon overrepresentation analysis using both up- and down-regulated genes meeting the $p_{adj}$ cutoff in A). D-G) Heat maps showing expression levels of ATF4 target genes, genes involved in myelination, and genes involved in Phe metabolism. All heatmaps are colored with Z-scores.

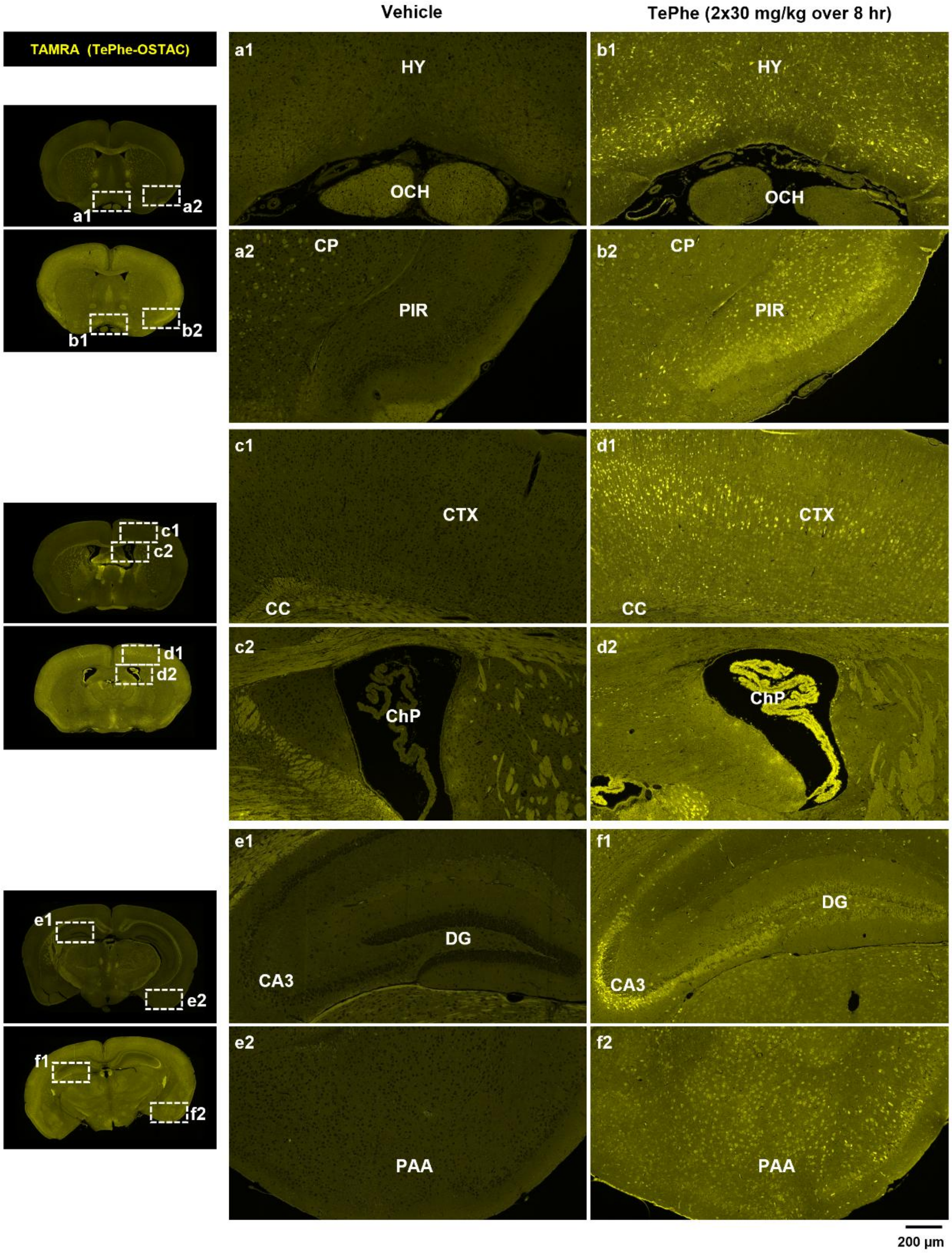


**Figure S15. TePhe incorporates widely across brain regions.** Matching regions indicated by dotted rectangles (vehicle, **a1-f1**; TePhe, **a2-f2**) are shown at higher magnification. TePhe appears to diffuse readily across brain tissue, as neuronal cell bodies with bright OSTAC signal appear across multiple regions. HY = hypothalamus. OCH = optic chiasm. CP = caudoputamen. PIR = piriform area of cerebral cortex. CTX = cerebral cortex. CC = corpus callosum. ChP = choroid plexus within lateral ventricle. DG = dentate gyrus. CA3 = Ammon's horn subfield 3. PAA = piriform-amygdalar area. Assignments were made by matching against the Allen Brain Atlas.[1]

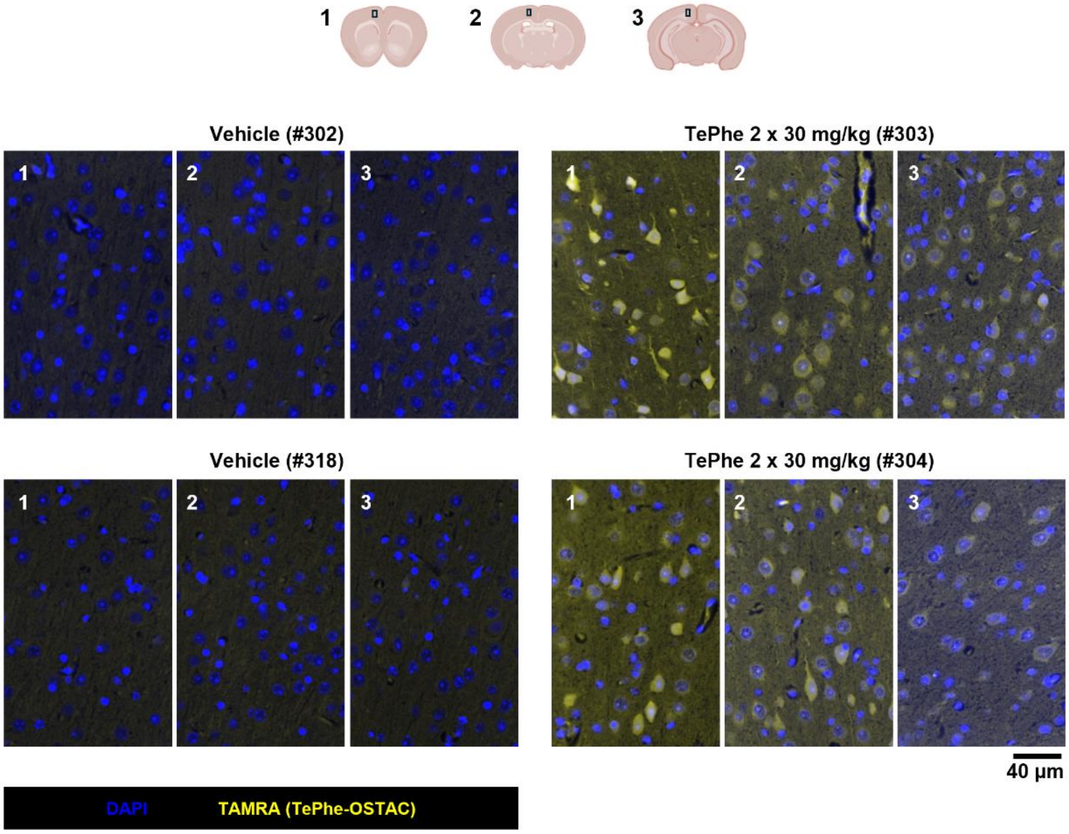


**Figure S16. TePhe signal in cell bodies of neurons within the cerebral cortex at 3 different brain depths.** Fluorescent OSTAC labeling reveals high signal intensity in the soma as well as proximal projections of individual neurons.

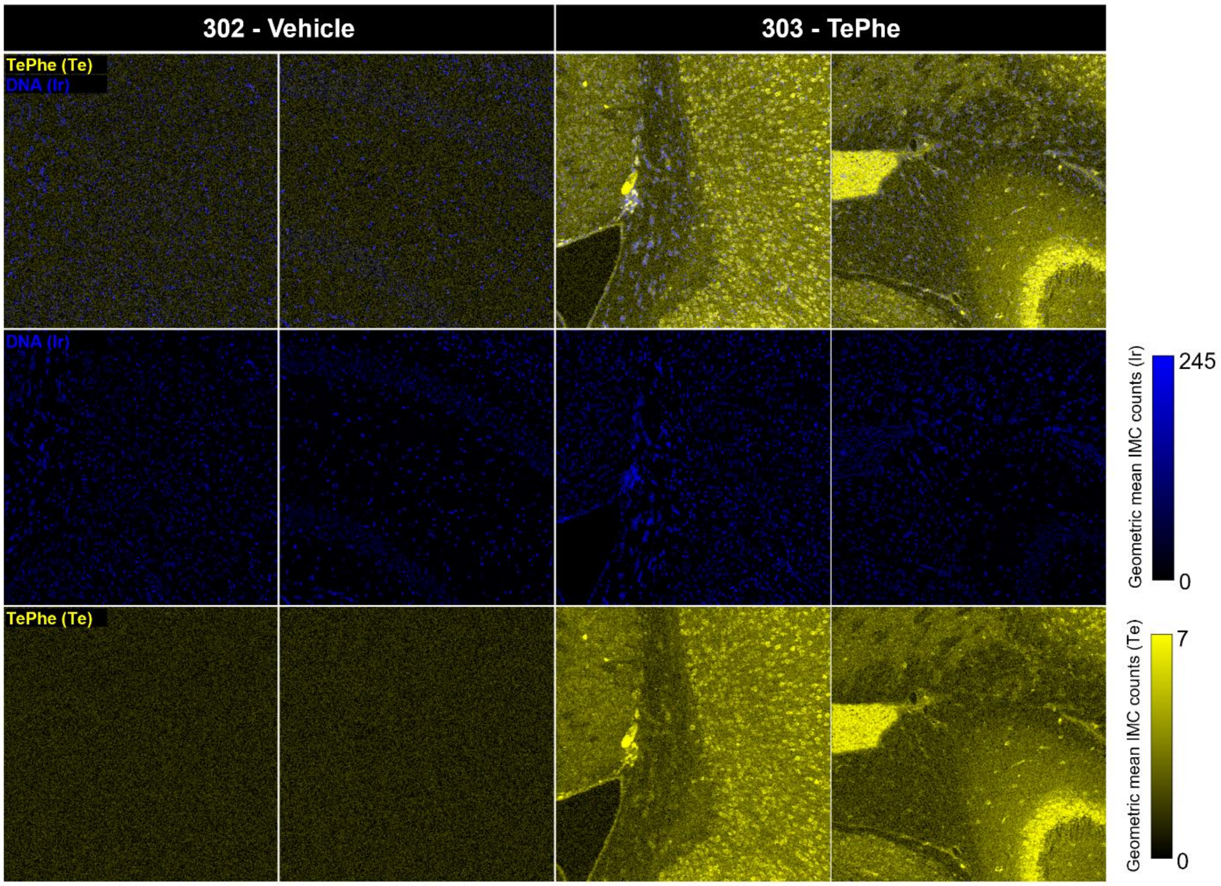


**Figure S17. Additional data associated with imaging mass cytometry (IMC) images shown in Fig. 3b.** All regions are 1 mm$^2$ in area. Two regions from a vehicle brain, of similar locations to those from the TePhe-treated brain, were imaged, showing low, uniform Te background in contrast to obvious morphological features visualized in the TePhe-treated sample. Ir = iridium-based DNA intercalator for identification of nuclei. Each channel registered intensity information for their respective elements only; false colours were assigned for intuitive viewing. Te signal = geometric mean of $^{126}$Te, $^{128}$Te, $^{130}$Te ($^{131}$Xe corrected). Ir signal = geometric mean of $^{191}$Ir, $^{193}$Ir. Images were thresholded to 99% max of the geometric mean signal in the based on the region with the highest signal.

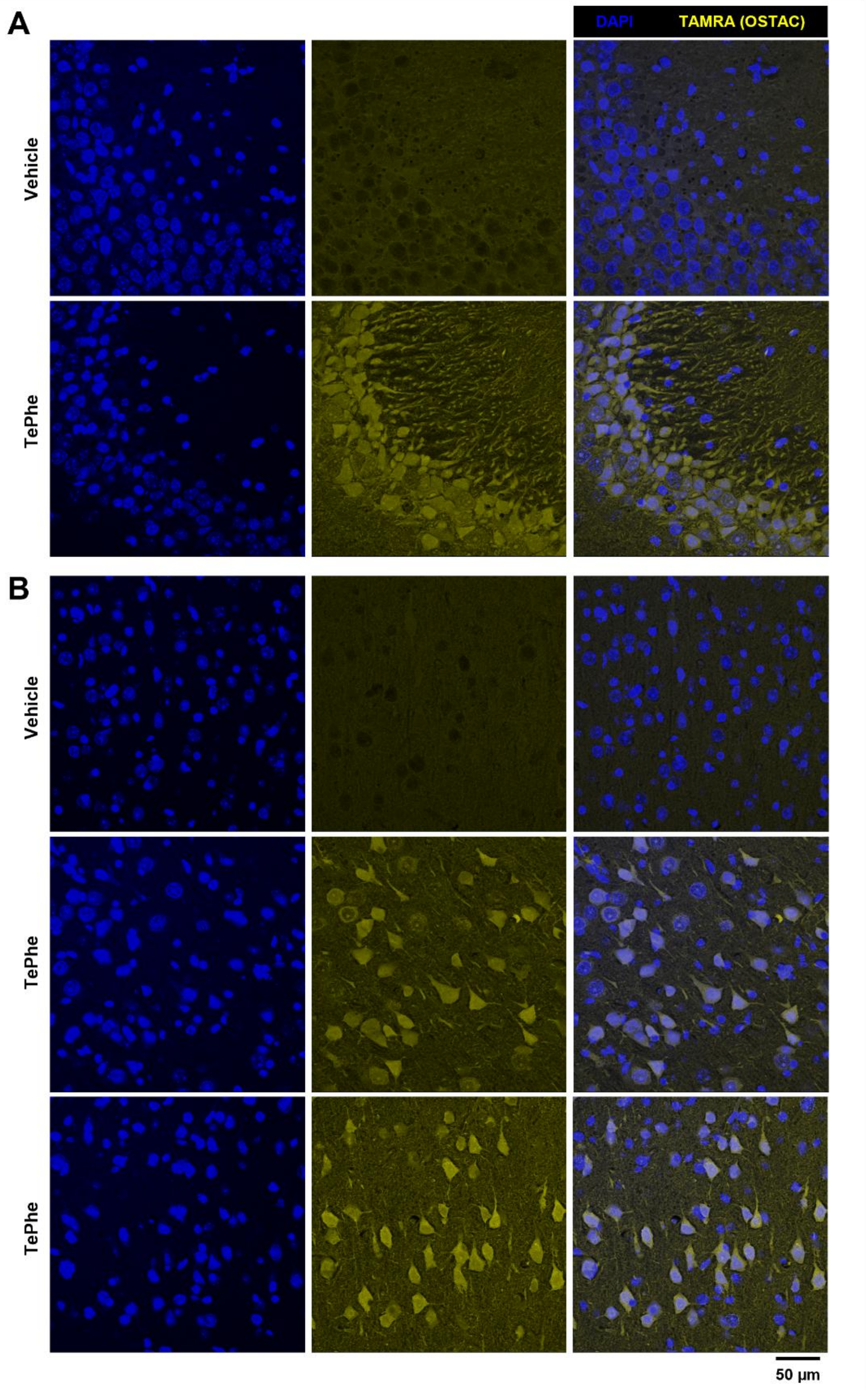


**Figure S18. Additional confocal imaging data from fluorescent OSTAC labeling of 20 µm sections.** A) CA3 region of vehicle brain compared with TePhe-treated (2 x 30 mg/kg over 8 hr) CA3 region shown in Fig. 3c, showing uniformly low background in the control sample. B) Comparison of vehicle and TePhe-treated cortical regions.

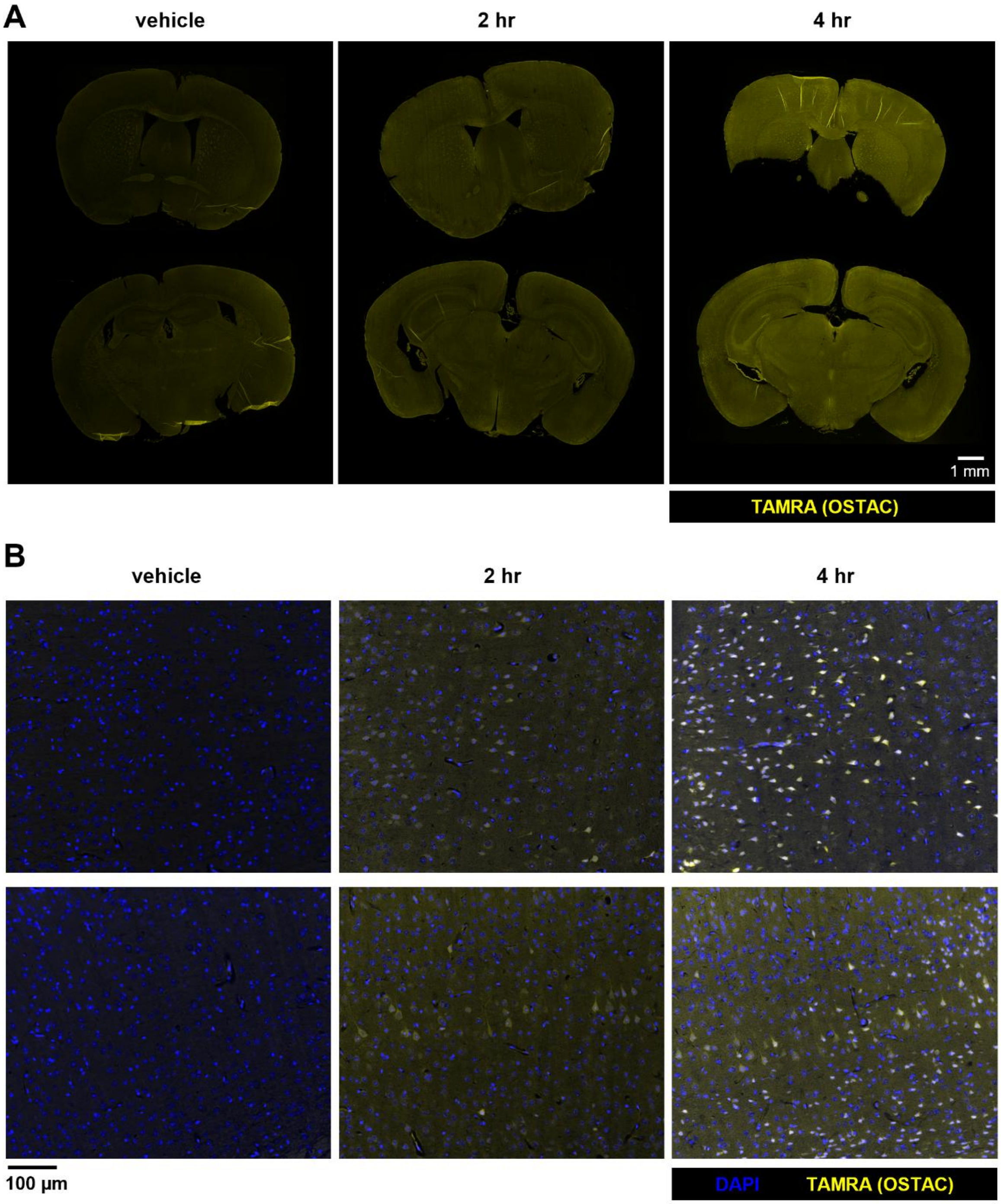


**Figure S19. Additional data from time course experiment shown in Fig. 3d.** A) Whole section view of vehicle and TePhe-treated (90 mg/kg) sections after fluorescent OSTAC labeling, showing increasing TAMRA intensity across the brain with increasing labeling time. B) Cortex visualized at high magnification, showing strong signal in neuronal cell bodies visible as early as 2 hr.

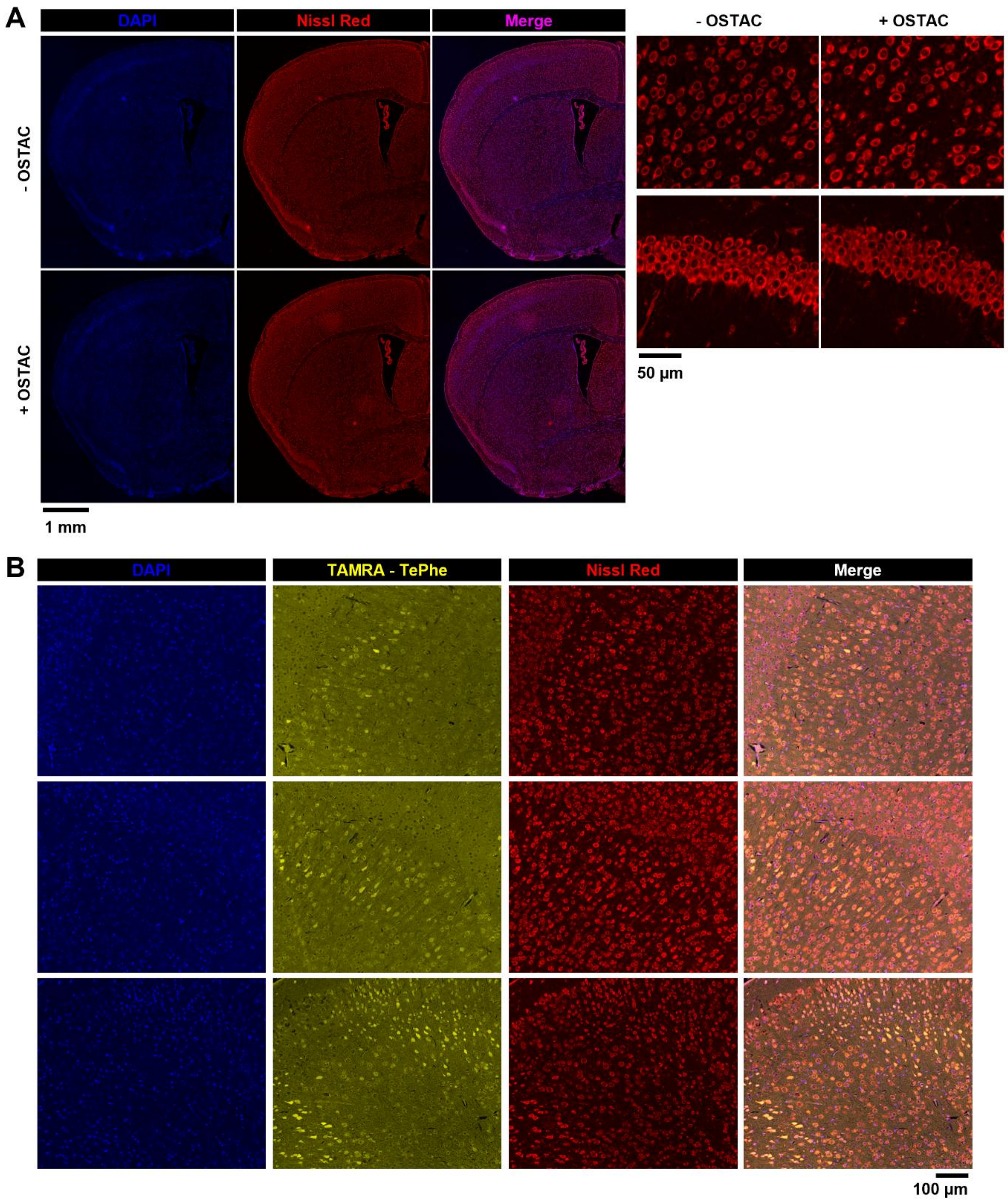


**Figure S20. Sequential combination of OSTAC and Nissl staining.** A) Serial sections were either untreated or subjected to OSTAC conditions prior to application of Nissl Red stain. Nissl staining pattern and intensity are effectively unchanged by OSTAC treatment. Left: coronal section of brain hemisphere. Right: neurons in cortex and hippocampus visualized at higher magnification. B) Additional images of the mouse cerebral cortex showing combined OSTAC and Nissl staining, with co-localization of TePhe and Nissl signal indicating protein synthesis in neuronal cell bodies.

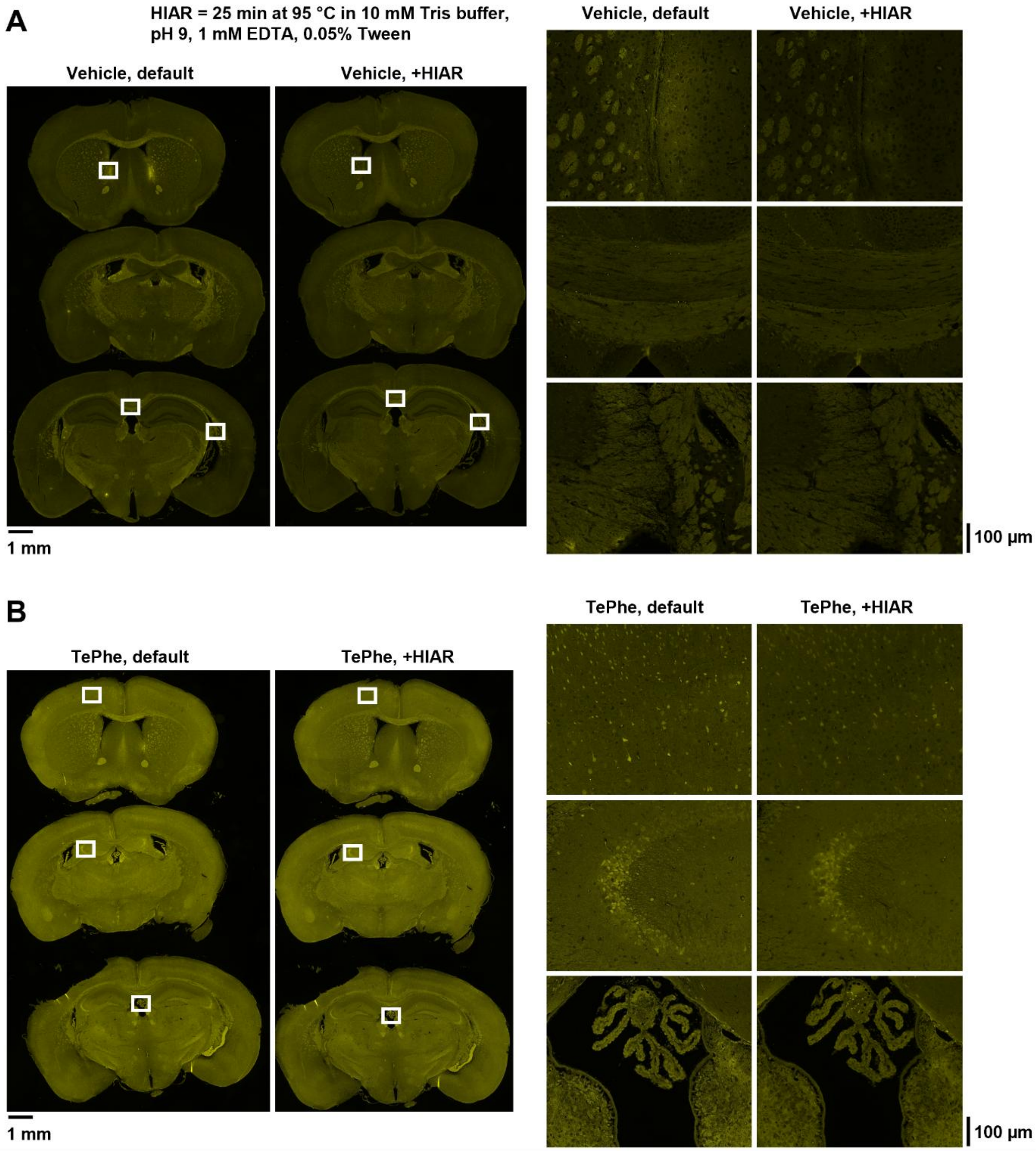


**Figure S21. Effect of heat-induced antigen retrieval (HIAR) on OSTAC labeling.** Serial sections of mouse brain (A, vehicle; B, TePhe) were subjected to fluorescent OSTAC labeling with BCN-TAMRA (yellow), either directly after tissue rehydration (default) or after heating at 95 °C in 10 mM Tris buffer, pH 9, 1 mM EDTA, 0.05% Tween 20 (HIAR). Right panel shows magnified regions indicated by white boxes on whole brain sections. HIAR conditions used appear to have no effect on the OSTAC background, and only slightly reduces the TePhe signal/sharpness of morphological features. Similar results are obtained after HIAR using 10 mM citrate buffer, pH + 0.5% Tween.

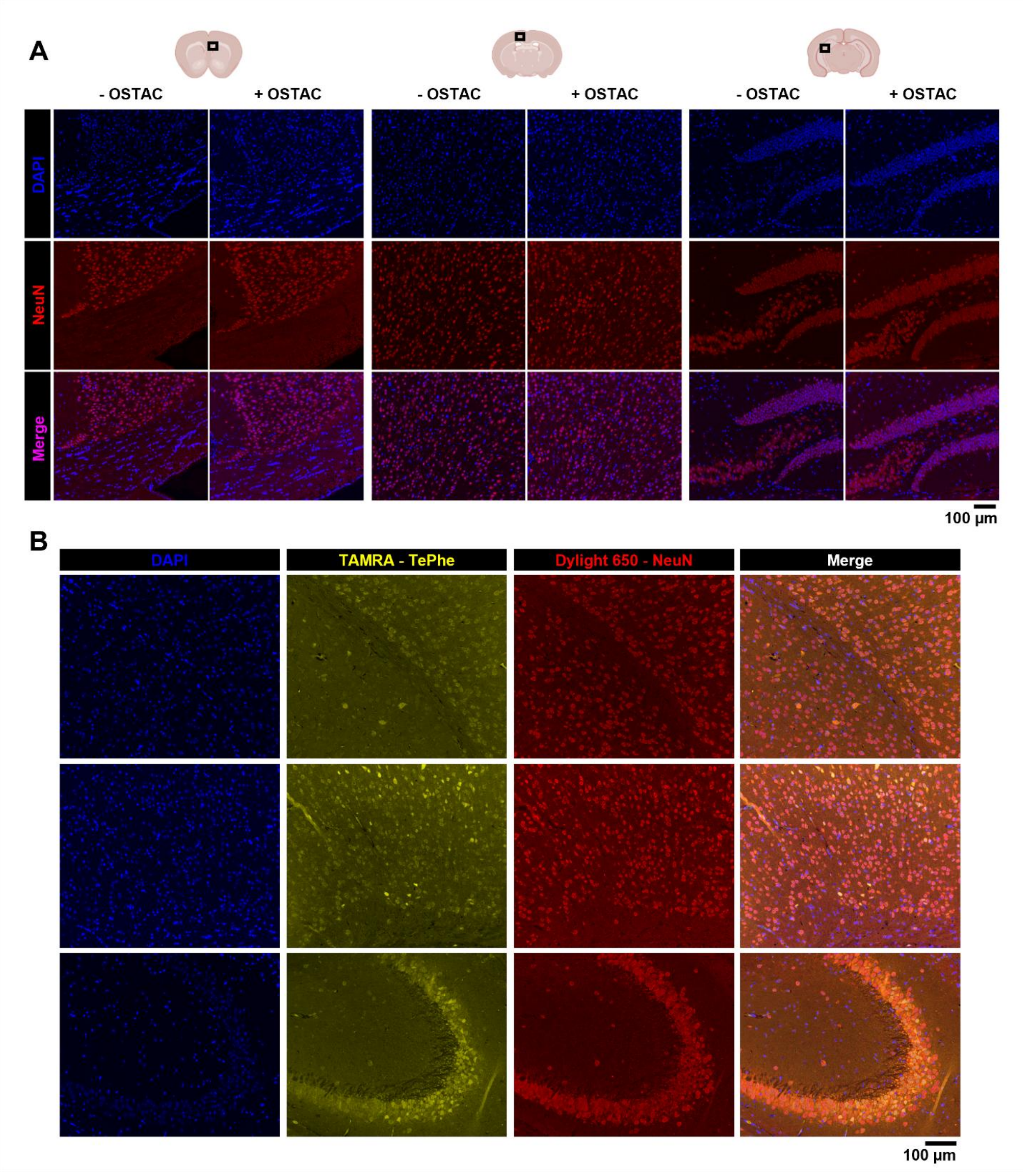


**Figure S22. Compatibility of OSTAC with NeuN (neuronal cell marker) immunofluorescence.** A) Sections were untreated or subjected to OSTAC conditions before NeuN staining. OSTAC has minimal effect on NeuN staining pattern and intensity. Strong signal is observed in neuronal nuclei of the cerebral cortex and hippocampus while being absent from the corpus callosum, which is composed primarily of oligodendrocytes. B) Additional images showing visualization of protein synthesis by OSTAC and identification of neurons by NeuN immunofluorescence in mouse cerebral cortex and hippocampal CA3 region.

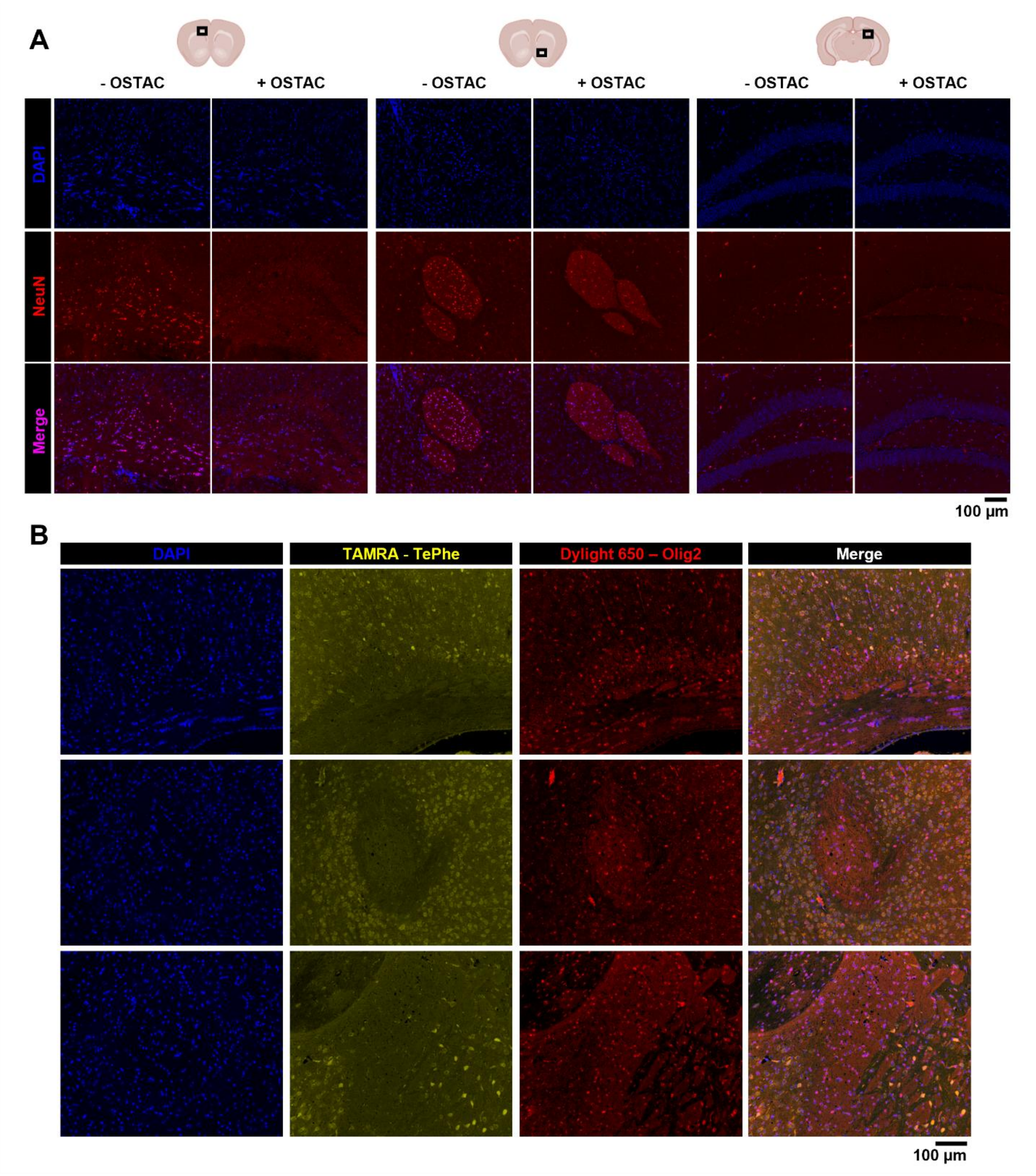


**Figure S23. Compatibility of OSTAC with Olig2 (oligodendrocyte marker) immune-fluorescence.** A) Sections were untreated or subjected to OSTAC conditions before Olig2 staining. OSTAC causes a notable reduction in Olig2 signal intensity, but residual signal maintains the correct nuclear localization in oligodendrocytes. B) Additional images showing combination of OSTAC and Olig2 staining in the corpus callosum and white matter tracts of the brain. Although Olig2 immunofluorescence is diminished by OSTAC treatment, the residual signal remains serviceable for identification of oligodendrocytes upon threshold and contrast adjustment.

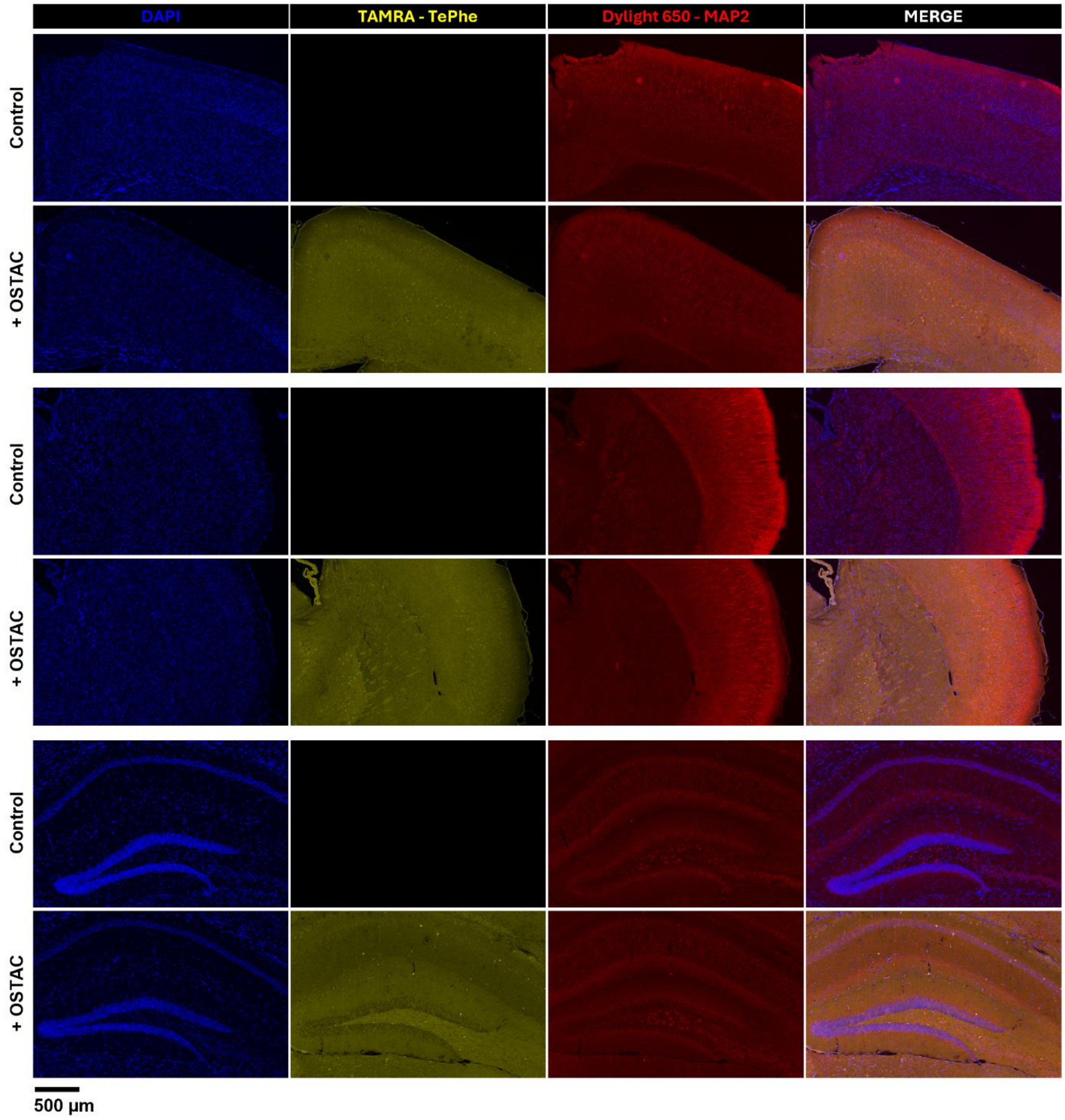


**Figure S24. Compatibility of OSTAC with MAP2 (microtubule-associated protein 2) immunofluorescence.** Control sections (vehicle brain, not OSTAC-treated) and fluorescently OSTAC-labeled sections (TePhe-treated brain) were subjected to MAP2 antibody staining. TePhe incorporation and OSTAC treatment do not appear to hinder MAP2 staining; the distinct patterns of MAP2 along neuronal projections in the cerebral cortex and hippocampus are retained.

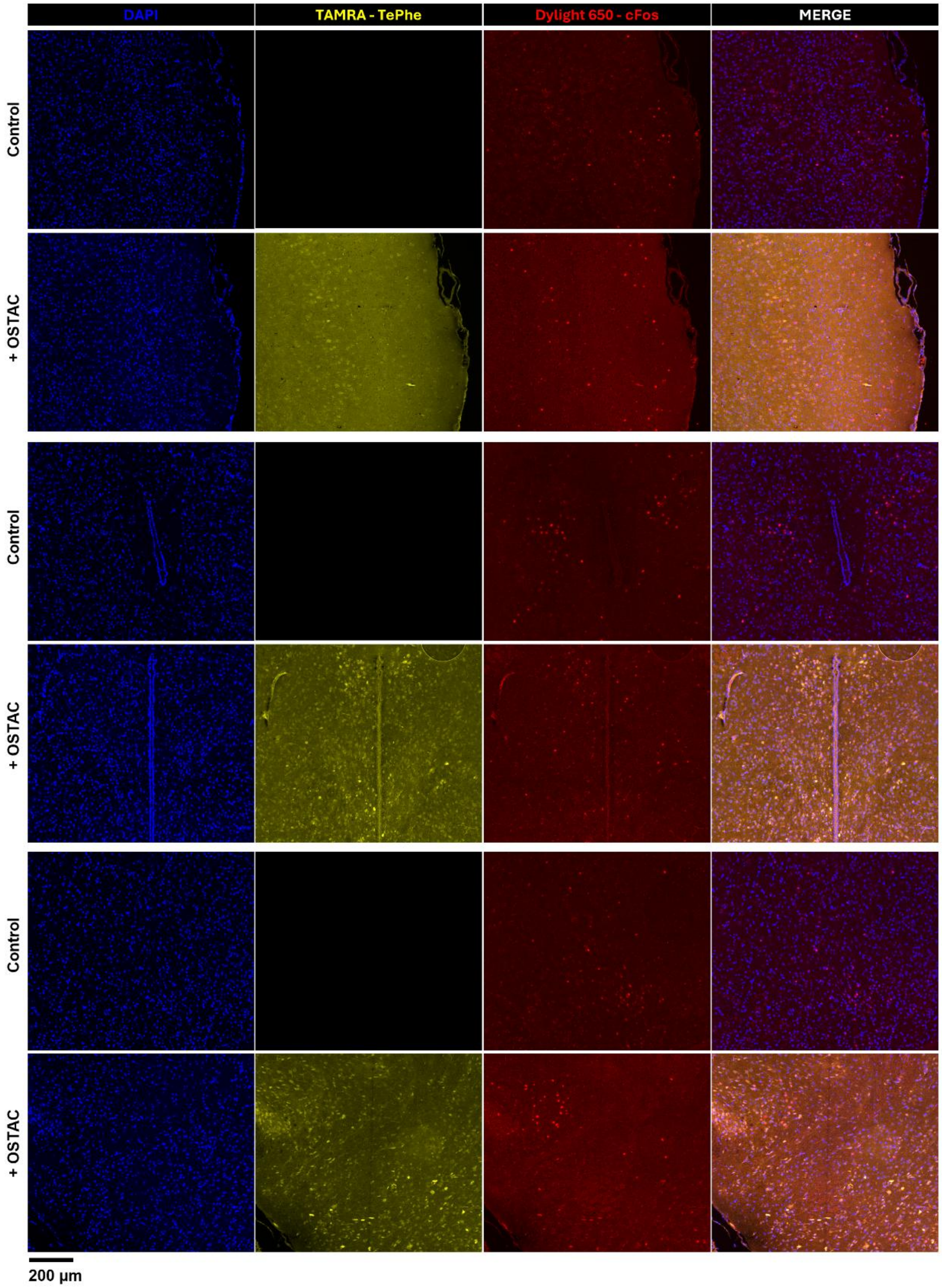


**Figure S25. Compatibility of OSTAC with c-Fos (transcription factor and cellular activity marker) immunofluorescence.** Control sections (vehicle brain, not OSTAC-treated) and fluorescently OSTAC-labeled sections (TePhe-treated brain) were subjected to c-Fos antibody staining. TePhe incorporation and OSTAC treatment do not appear to hinder staining; the distinct nuclear pattern of c-Fos can be seen in both samples. Background is very slightly elevated in the OSTAC-treated sample.

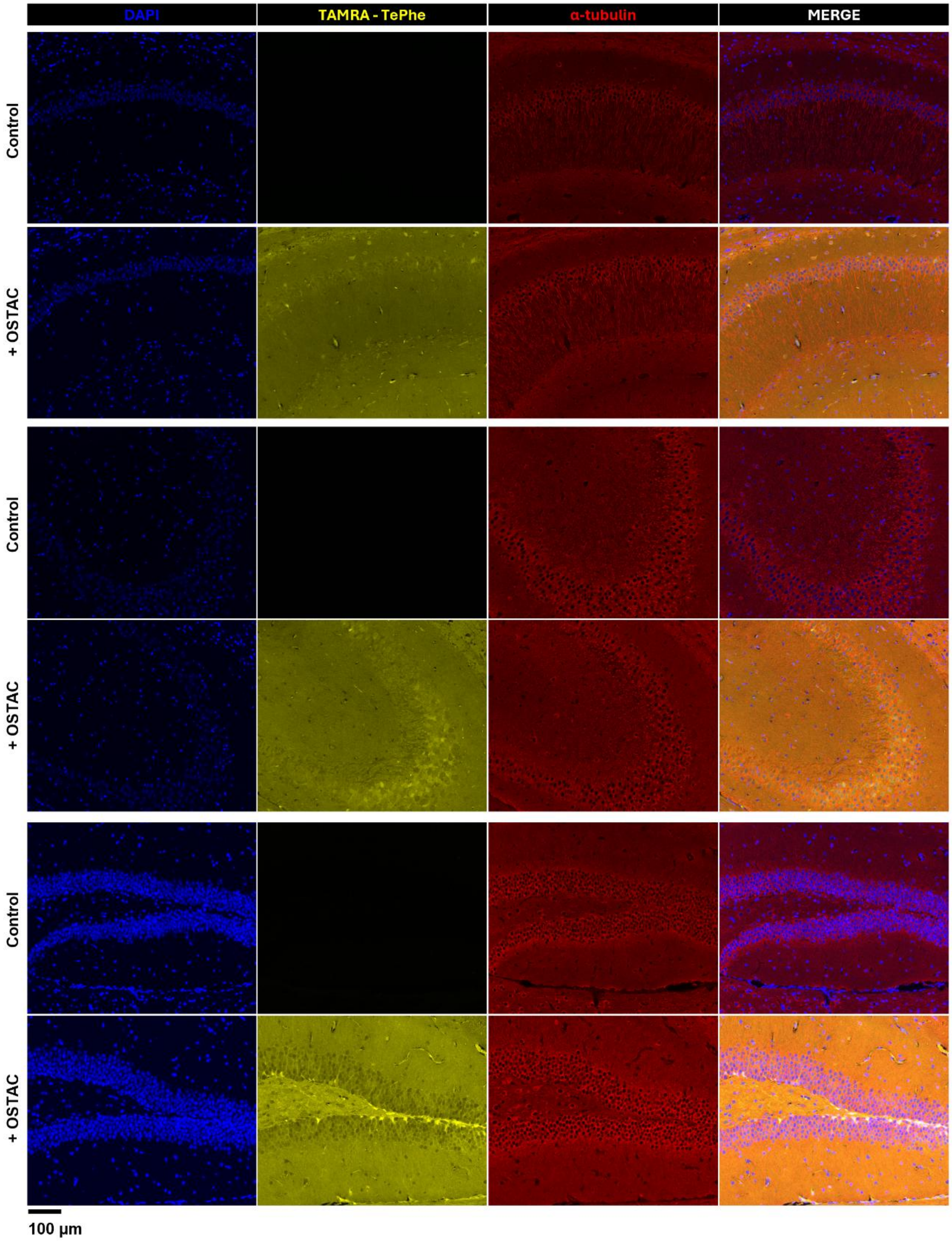


**Figure S26. Compatibility of OSTAC with α-tubulin (microtubule marker) immune-fluorescence.** Control sections (vehicle brain, not OSTAC-treated) and fluorescently OSTAC-labeled sections (TePhe-treated brain) were subjected to α-tubulin antibody staining. TePhe incorporation and OSTAC treatment do not appear to alter the distinct staining patterns visible within the hippocampus.

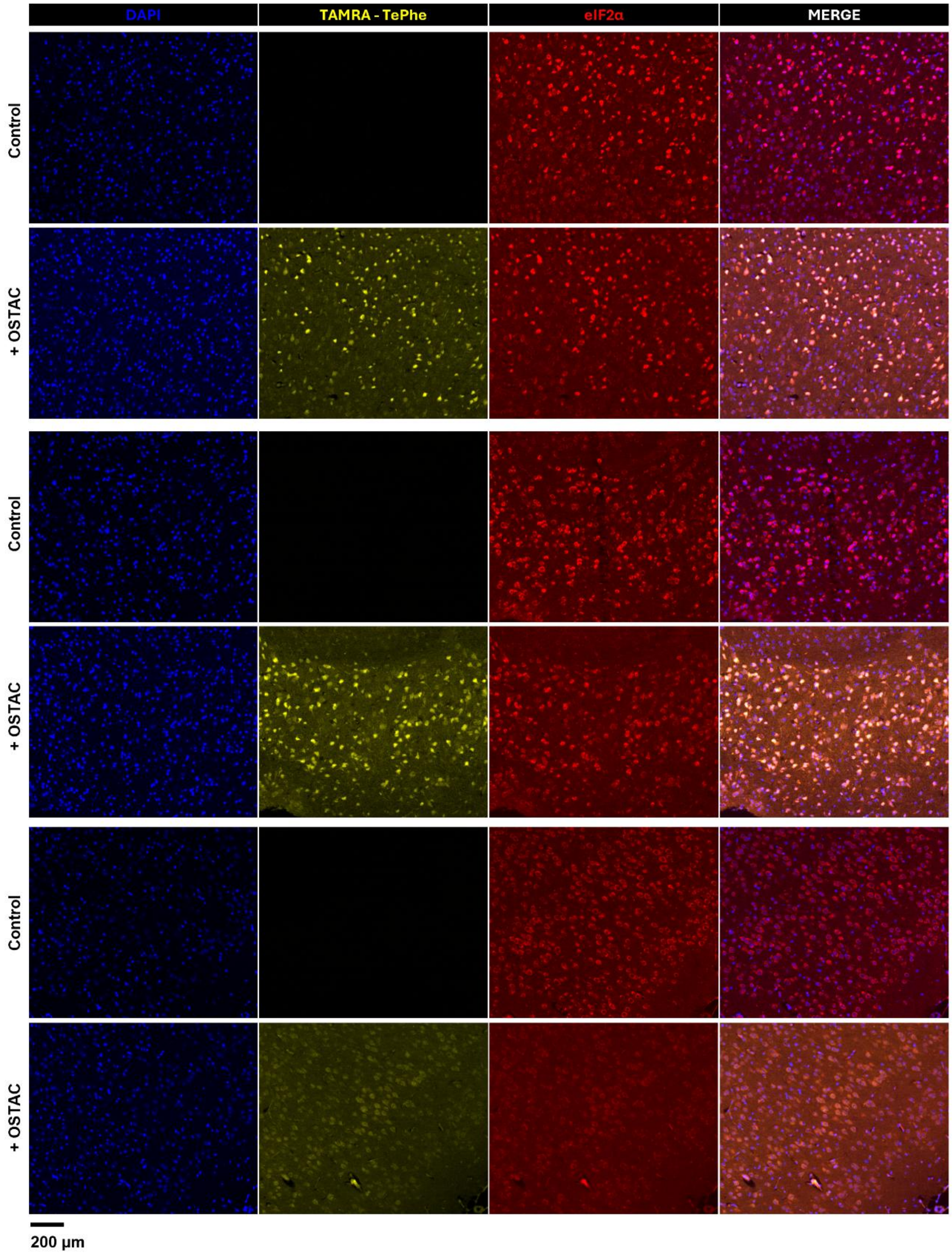


**Figure S27. Compatibility of OSTAC with eIF2α (translation initiation factor) immunofluorescence.** Control sections (TePhe-treated brain, not OSTAC-treated) and fluorescently OSTAC-labeled sections (TePhe-treated brain) were subjected to eIF2α antibody staining. Consecutive rows show matched brain regions. OSTAC treatment does not appear to alter the overall patterns of eIF2α.

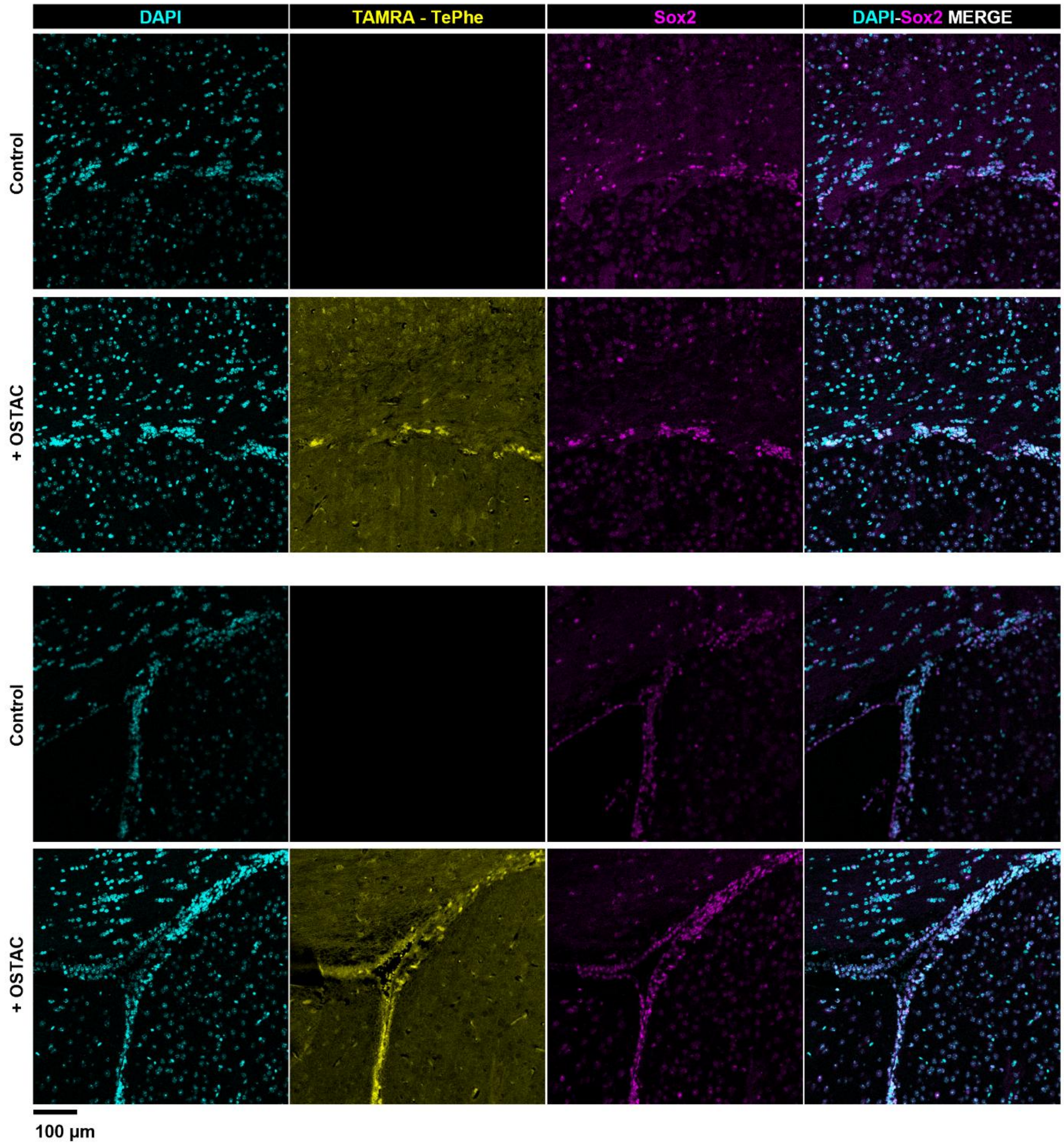


**Figure S28. Compatibility of OSTAC with Sox2 (neural stem and progenitor cell marker) immunofluorescence.** Control sections (vehicle brain, not OSTAC-treated) and OSTAC-labeled sections (TePhe-treated brain) were subjected to Sox2 antibody staining. Overlay of DAPI and Sox2 channels shows that OSTAC has minimal effect on the nuclear pattern of Sox2 staining or its localization.

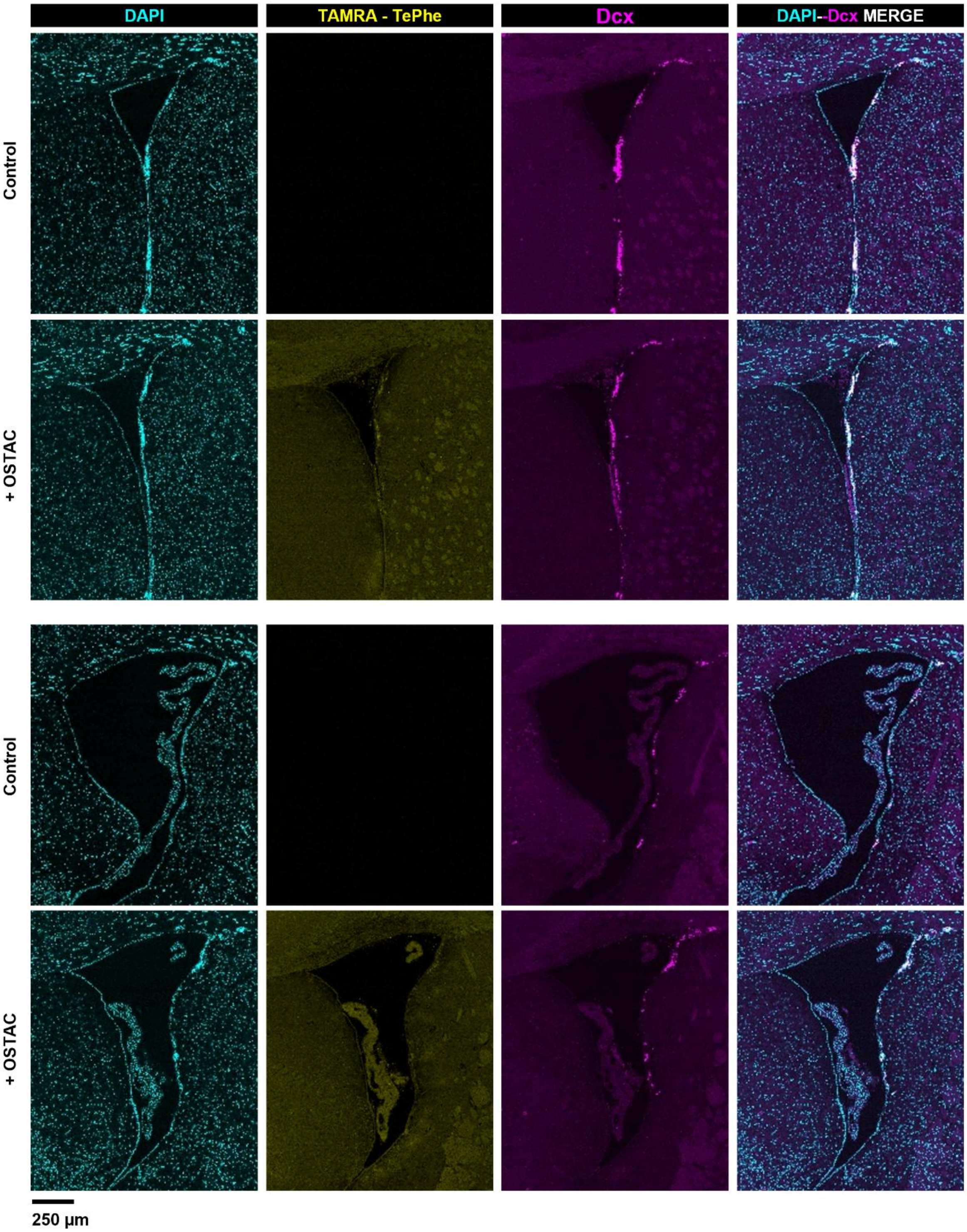


**Figure S29. Compatibility of OSTAC with doublecortin (Dcx; microtubule-associated protein and marker of migrating neuroblasts) immunofluorescence.** Control sections (vehicle brain, not OSTAC-treated) and OSTAC-labeled sections (TePhe-treated brain) were subjected to Dcx antibody staining. Overlay of DAPI and Dcx channels shows that the Dcx staining pattern surrounding the lateral ventricles is unchanged by OSTAC treatment.

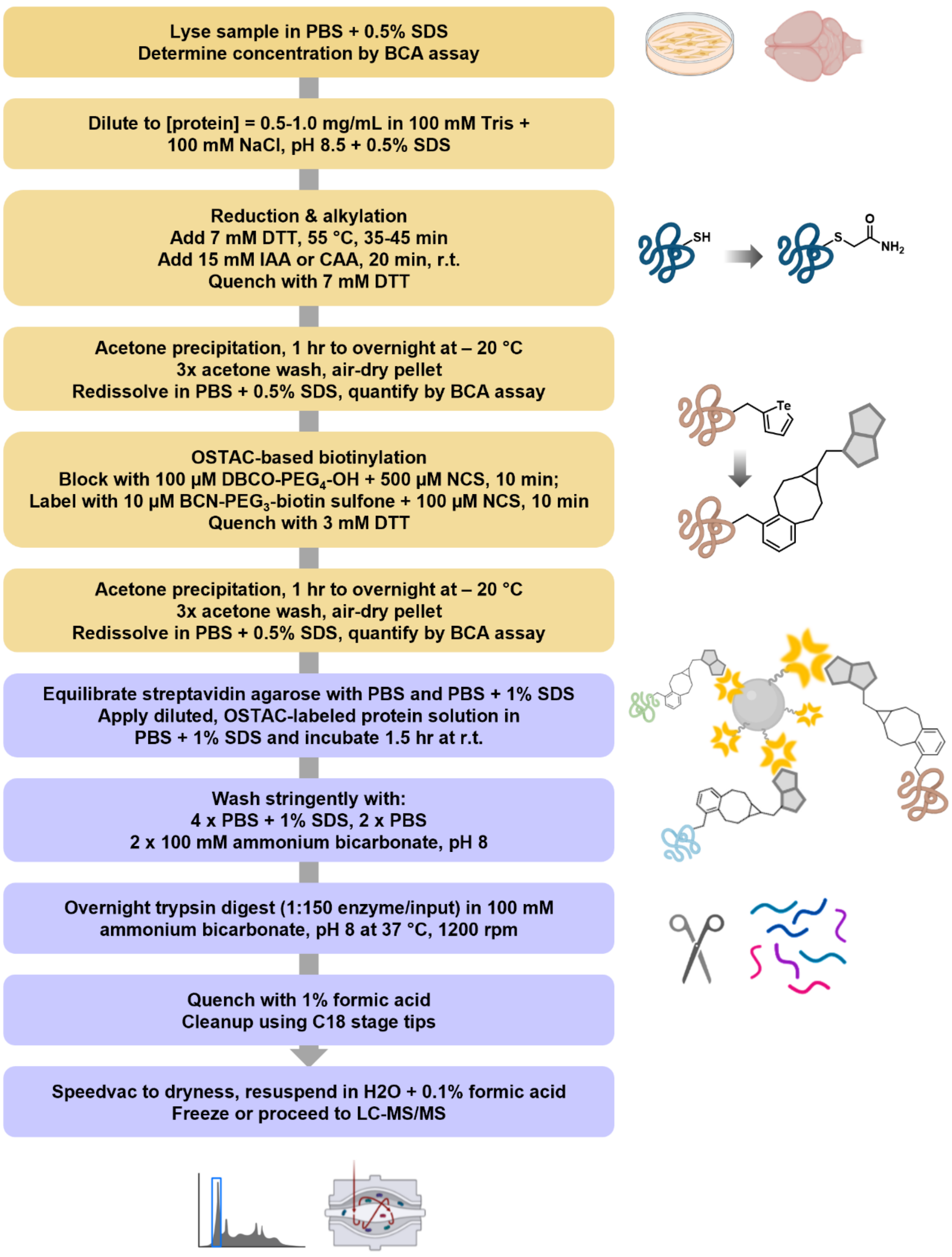


**Figure S30. Schematic of OSTAC-based biotinylation of newly synthesized proteins followed by affinity capture for LC-MS/MS.**

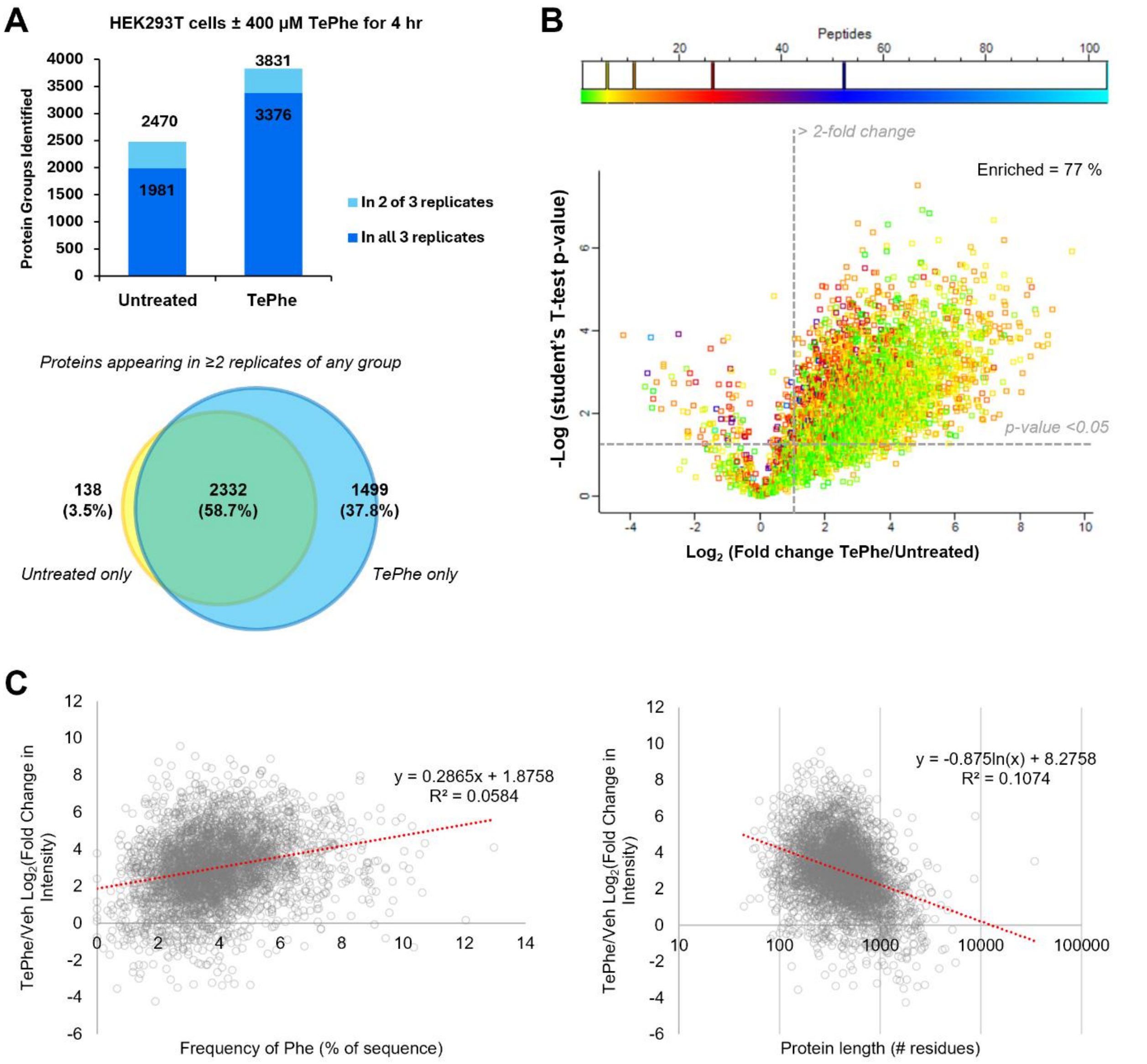


**Figure S31. OSTAC-based proteomics using HEK293T lysates.** A) Proteins groups detected after OSTAC-based enrichment for TePhe-containing proteins in vehicle and TePhe-treated (400 µM, 4 hr) cell lysates. Venn diagram shows protein groups identified in either or both conditions (a protein group is counted as identified if appearing in ≥ 2 of 3 replicates). B) Volcano plot comparing protein intensities (Student's t-test) between vehicle and TePhe-treated lysates, showing enrichment of newly synthesized proteins over background in TePhe-treated samples. C) Degree of enrichment has minimal correlation with Phe content or length of the target sequence. OSTAC-based enrichment does not appear to biased for longer proteins or those with more Phe residues.

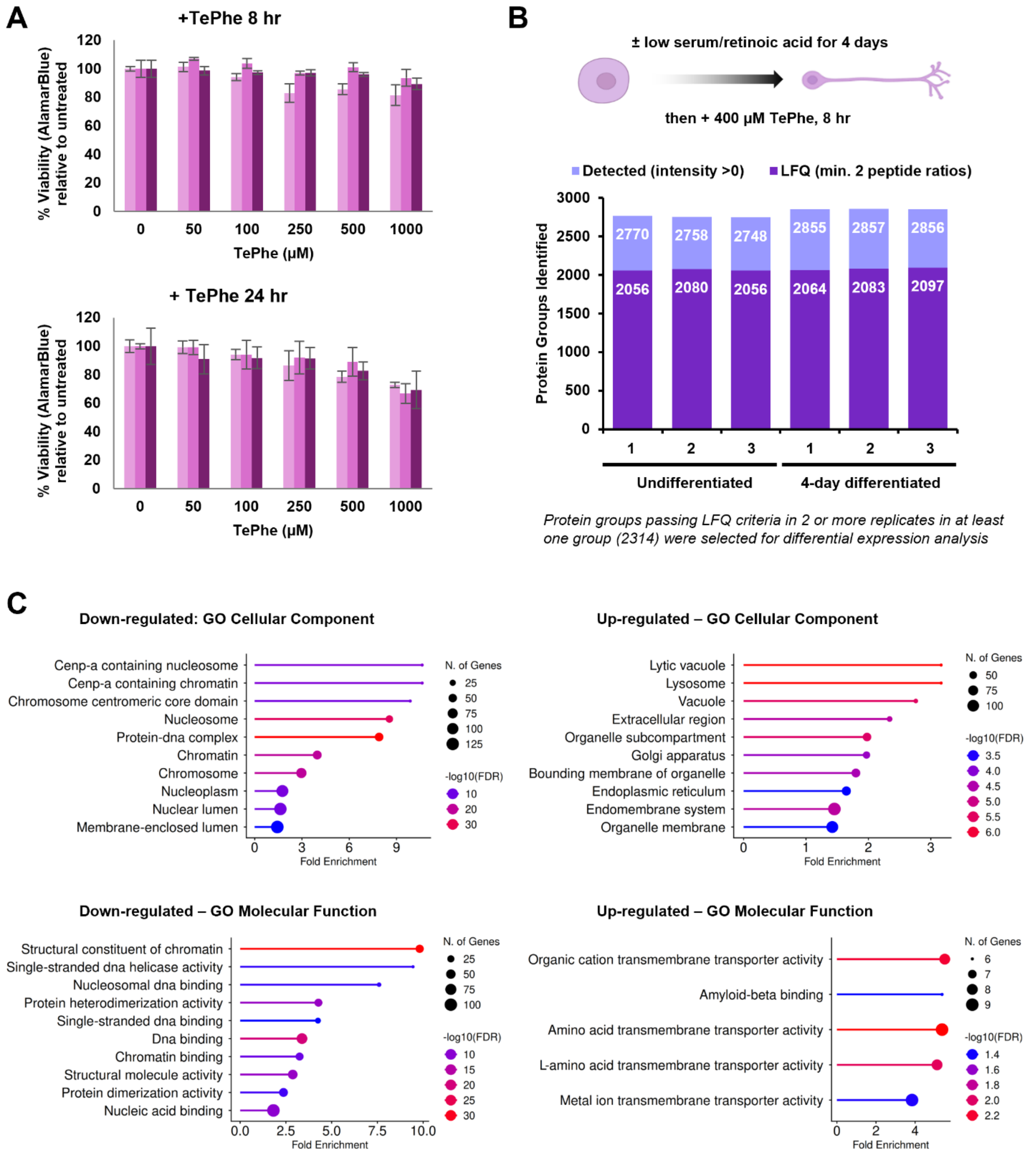


**Figure S32. OSTAC-based proteomics applied to differentiation of N2a murine neuroblastoma cells.** A) AlamarBlue assay to determine TePhe toxicity in N2a cells. Bars indicate 3 biological replicates (error bars indicate standard deviation between 4 technical replicates). B) Protein groups identified across biological replicates in undifferentiated and differentiated groups. C) Additional pathway analysis results for significantly down- and up-regulated proteins shown in Fig. 6a. Expected changes relating to down-regulation of DNA replication and up-regulation of transport-associated cellular components are observed.

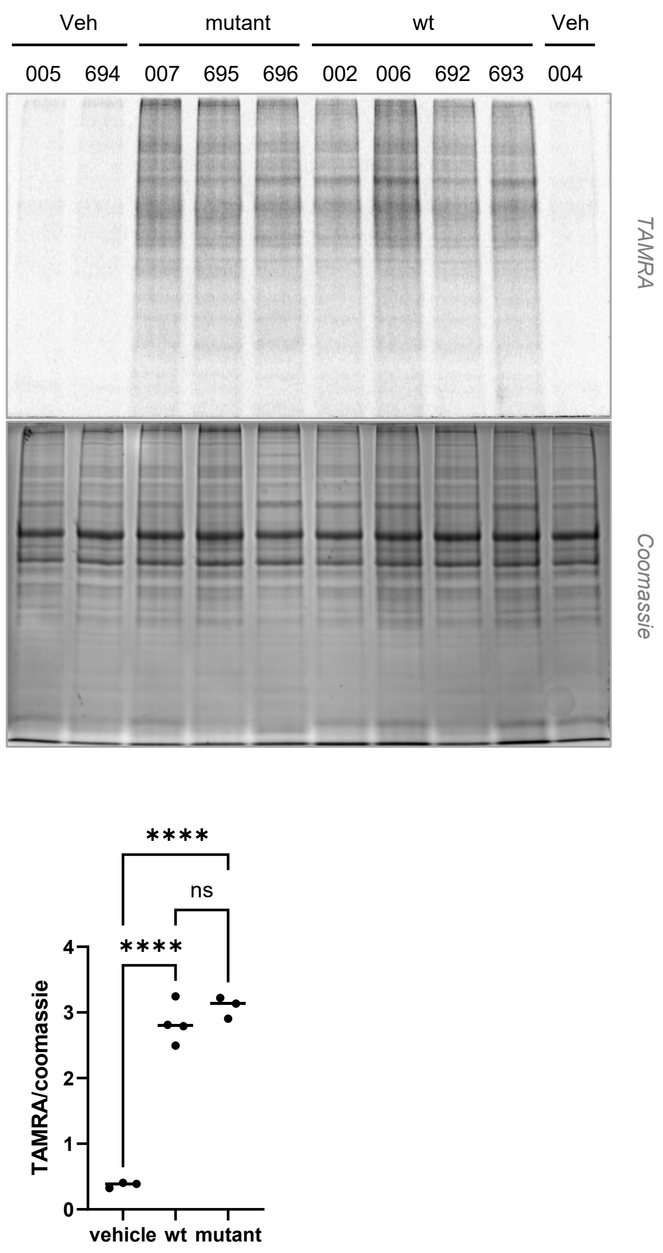


**Figure S33. In-gel fluorescent detection of TePhe incorporation to compare bulk protein synthesis.** Brain stem incorporation after a 2x 45 mg/kg IP injection in wt and mutant mice. Quantification of TAMRA signal relative to Coomassie. Each datapoint represents one mouse (*n* = 3, 4, 3; one-way ANOVA, **** *p* <0.0001).
