## Supplementary Information for "A Metabolic Labeling Strategy for Tracking Protein Synthesis in Complex Biological Systems"

**Table of Contents**


| | |
|---|---|
| **Section 1. TePhe synthesis** | 2 |
| Procedures for large-scale TePhe synthesis | 2 |
| NMR spectra of TePhe | 4 |
| **Section 2. Table of ages and genotypes of experimental animals** | 5 |
| **Section 3. List of stains and antibodies** | 6 |
| **Section 4. Procedures for equilibrating/washing streptavidin beads** | 7 |
| **References** | 8 |

### Section 1. TePhe synthesis

#### *General*

Large-scale TePhe synthesis was carried out by simplifying an existing synthetic route.[1] L-propargylglycine was purchased from Aaron Chem (AR002NB0). (Bromoethynyl)triiso-propylsilane was prepared as previously described.[1] All other reagents and solvents were purchased from Sigma. TLC plates (TL-BM3101) were purchased from Santai. Sonication was carried out using a Transsonic 420 bath sonicator. NMR spectra were acquired on a 500 MHz Agilent DD2 spectrometer equipped with an XSens C13 cold probe.

#### *Synthesis of (S)-2-amino-7-(triisopropylsilyl)hepta-4,6-diynoic acid* (1).

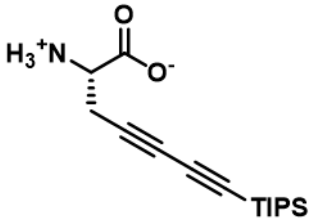


A 50 mL RBF under argon atmosphere was charged with a mixture of degassed $H_2O$ (20 mL) and *n*-butylamine (7 mL) and cooled using an ice bath. CuCl (56 mg, 0.57 mmol, 0.05 eqv) and hydroxylamine hydrochloride (330 mg, excess to Cu catalyst) were added in rapid succession, which produced a transient blue colour followed by return to a clear and colourless solution. L-propargylglycine solid (1.27 g, 11.4 mmol, 1.0 eqv) was then added and the mixture stirred until all solids dissolved. (Bromoethynyl)triisopropylsilane (2.97 g, 11.4 mmol, 1.0 eq.) was injected over the course of 5 min. The mixture was stirred on ice for 25 min, after which the contents were allowed to warm to room temperature. The reaction initially formed a yellow emulsion which cleared by 2.5 hr. After 4 hr of stirring, the reaction was quenched by transfer to a separatory funnel charged with 30 mL $H_2O$ and 50 mL 2M HCl. The mixture was extracted three times with diethyl ether (150, 60, 60 mL). The combined organic phase was washed twice with 0.5M HCl (50 mL), once with brine (50 mL), and dried over $Na_2SO_4$. Filtered product was concentrated under reduced pressure to yield a thick yellow gel which transformed into a foamy, off-white solid after drying overnight (3.5 g). This material was used directly for the next step of synthesis.

#### *Synthesis of L-2-tellurienyllalanine (TePhe).*

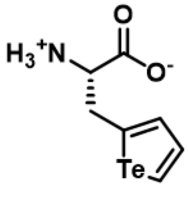


A 150 mL RBF was charged with Te metal powder (Sigma 26418, 0.896 g, 7.0 mmol, 1.2 eqv to **1**) and sodium borohydride (2.27 g, 6.0 mmol, 8.5 eqv to Te). The flask was purged with argon, after which degassed $dH_2O$ (42 mL) was added over rapid stirring. Two half-filled argon balloons were connected to the reaction vessel (leaving volume for hydrogen gas evolution). The mixture was allowed to stir for 1 hr at room temp. followed by heating to 44 °C. The reaction mixture gradually changed from gray-black to dark purple, then to light pink as Te was reduced.

Concurrently, TIPS-deprotection of **1** was carried out in a 100 mL RBF. On ice, **1** (1.78 g, 6.1 mmol, 1.0 eqv) was dissolved in anhydrous THF (8.0 mL), followed by addition of tetrabutylammonium fluoride (TBAF) solution (1 M in THF, 8.0 mL, 8.0 mmol, 1.3 eqv). The mixture was briefly sonicated to break up solids, followed by stirring for 40 min at 0 °C. The mixture was then concentrated under reduced pressure, the flask purged with argon, and the contents resuspended in degassed ethanol (45 mL).

Resuspended **1** was transferred by syringe to the vessel containing the Te reduction reaction. The injection rapidly caused the reaction mixture to turn orange. The mixture was allowed to stir under argon for 17 hr at 44 °C, after which the solution was found to be dark purple-black in colour. The mixture was removed from heat and allowed to stir, open to atmosphere, for 20 min. Excess sodium borohydride was quenched by addition of acetone (10 mL). The reaction mixture was acidified to pH 7 by addition of 2M HCl (25 mL) and stirred for an additional 45 min to fully precipitate unreacted Te. Reaction contents were filtered over a tightly-packed celite plug. The resulting yellow solution was concentrated under reduced pressure to remove organic solvents. The remaining aqueous solution was transferred to a new vessel, leaving behind small amounts of separated, oily material. The aqueous layer was flash frozen in liquid nitrogen and lyophilized to dryness (48 hr) to yield crude TePhe for purification by column chromatography.

Lyophilized solids were resuspended in methanol (50 mL) and sonicated for 20 min to form a very fine, cream-coloured suspension, which was mixed with minimal silica for dry loading. The mixture was dried *in vacuo* to yield a chalky solid, which was crushed until a fine, loose powder was obtained. A 5 cm diameter column was packed with a 2 cm tall silica plug, which was equilibrated with a gradient of methanol in dichloromethane (0, 5, 10, 15, 20, 30, 40, 50, 30, 20, 10, 5, and 0%, 100 mL each). The crude-silica mixture was dry loaded onto the column, followed by elution with a gradient of 0, 5, 7.5, 7.5, 10, 12.5, 15, 17.5, 20, 23, 25, 28, 30, 33, 37, 40% (100-200 mL each) methanol in dichloromethane. Starting at 10% methanol, eluent was collected in 10 mL fractions, with fractions 24-92 exhibiting UV activity on TLC corresponding to TePhe. Groups of 10 to 12 consecutive fractions were pooled, concentrated, and assessed by $^1$H NMR spectroscopy. Combined pure fractions (fractions 50-92) yielded a fluffy white solid (835 mg, 3.1 mmol, 51%). Impure fractions containing traces of tetrabutylammonium salts were pooled for re-purification by subsequent rounds of chromatography.

**$^1$H NMR** (500 MHz, MeOD-d4): δ 8.87 (dd, *J* = 7.0, 1.2 Hz, 1H, tellurophene -Te-C***H***-CH-CH-), 7.62 (dd, *J* = 7.0, 3.9 Hz, 1H, tellurophene -Te-CH-C***H***-CH-), 7.51 (dt, *J* = 3.9, 1.2 Hz, 1H, tellurophene -Te-CH-CH-C***H***-), 3.74 (dd, *J* = 7.0, 4.5 Hz, 1H, amino acid Hα), 3.48 (ddd, *J* = 15.2, 4.5, 1.2 Hz, 1H, amino acid Hβ), 3.40 (ddd, *J* = 15.2, 7.0, 1.1 Hz, 1H, amino acid Hβ′).

**$^{13}$C NMR** (126 MHz, MeOD-d4): δ 173.22, 143.04, 139.34, 137.44, 128.67, 58.29, 39.09.

$^{1}H$ NMR, 500 MHz, MeOD, 298 K

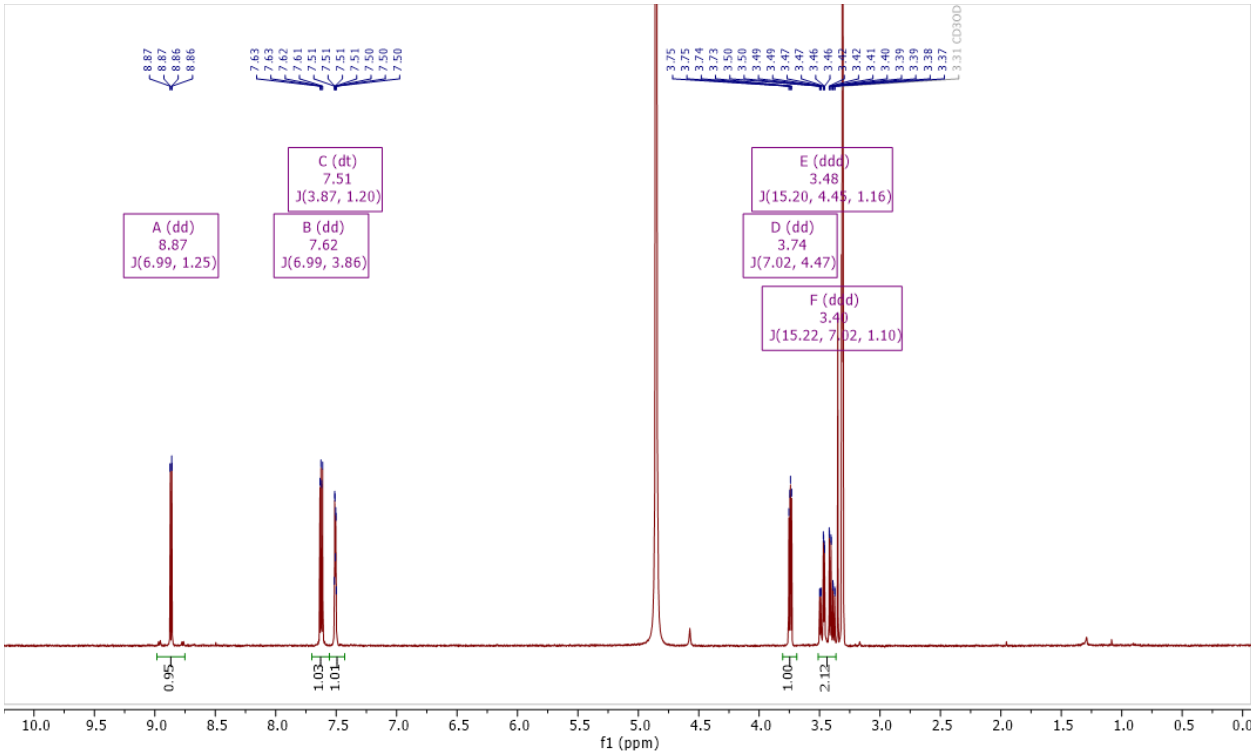


$^{13}C$ NMR, 126 MHz, MeOD, 298 K

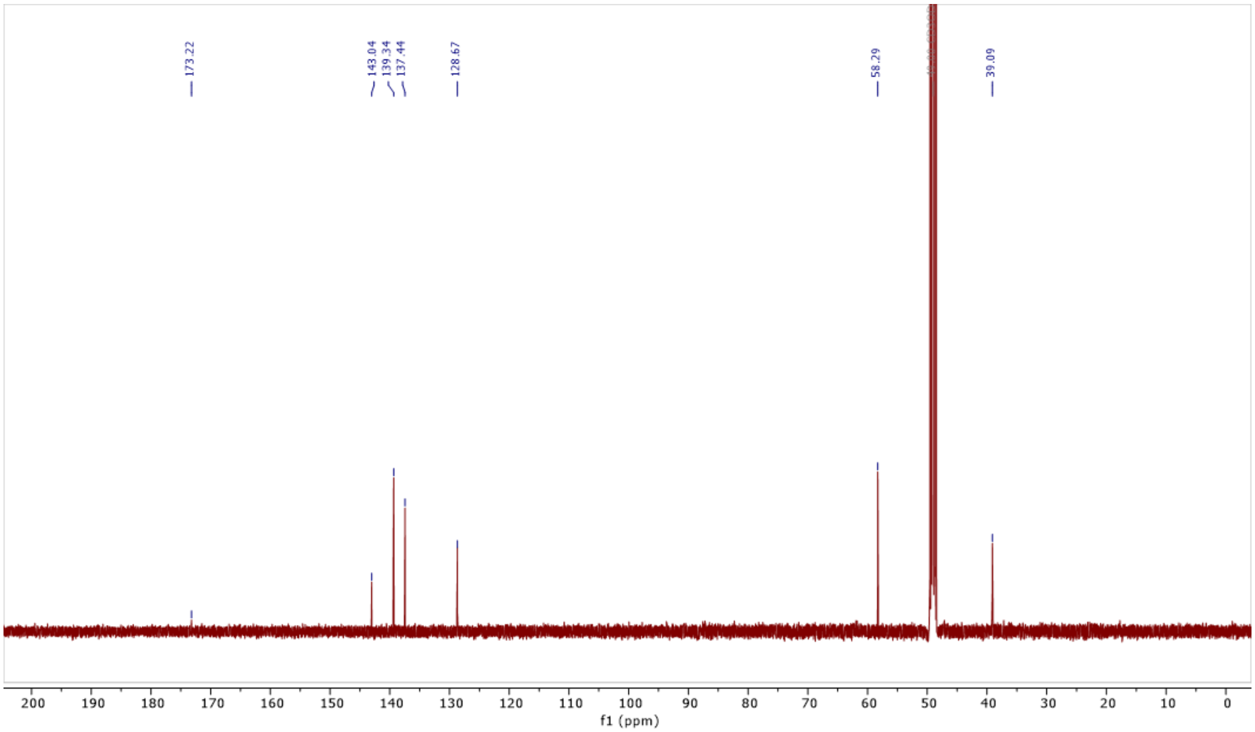

**Section 2. Table of ages and genotypes of experimental animals.**

| Experiment | Age | Sex | Genotype(s)* |
|---|---|---|---|
| IV injection, 60 mg/kg, 24 hr<br>Fluorescence microscopy of liver/intestines | 3 mo. | F | 6J/6J Cre$^-$ *Rars1*$^{loxP/-}$ |
| IP injection, 55 mg/kg, 8 hr<br>Brainstem TePhe content quant. | 9 mo. | F | 6J/6J Cre$^-$ *Rars1*$^{loxP/loxP}$ |
| IP injection, 90 mg/kg, 8 hr<br>Brainstem TePhe content quant. | 8 mo. | F/M | 6J/6J Cre$^-$ *Rars1*$^{loxP/loxP}$ |
| IP injection, 2×30 mg/kg over 8 hr<br>Brainstem TePhe content quant.<br>Fluorescence microscopy of brain sections | 8 - 10 mo. | F | 6J/6J Cre$^-$ *Rars1*$^{loxP/-}$<br>6J/6J Cre$^-$ *Rars1*$^{loxP/loxP}$ |
| IP injection, 2×45 mg/kg over 8 hr<br>Brainstem TePhe content quant. | 7 - 8 mo. | F/M | 6J/6J Cre$^-$ *Rars1*$^{loxP/-}$<br>6J/6J Cre$^+$ *Rars1*$^{-/-}$<br>6J/6J Cre$^+$ *Rars1*$^{loxP/-}$ |
| IP injection, 3×30 mg/kg over 24 hr<br>Brainstem TePhe content quant. | 10 mo. | F | 6J/6J Cre$^-$ *Rars1*$^{-/-}$<br>6J/6J Cre$^+$ *Rars1*$^{-/-}$<br>6J/6J Cre$^+$ *Rars1*$^{loxP/-}$ |
| IP injection, 3×30 mg/kg over 72 hr<br>Brainstem TePhe content quant.<br>Fluorescence microscopy of brain sections | 8 - 10 mo. | M | 6J/6J Cre$^-$ *Rars1*$^{loxP/loxP}$ |
| IP injection, 90 mg/kg, 0-4 hr timecourse blood collection and determination of free TePhe in serum | 2.5 - 4 mo. | M | 6J/6N Cre$^-$ *Rars1*$^{loxP/-}$<br>6J/6J Cre$^-$ *Rars1*$^{loxP/-}$<br>6J/6J Cre$^-$ *Rars1*$^{loxP/loxP}$ |
| IP injection, 90 mg/kg, 0-4 hr timecourse Fluorescence microscopy of brain sections | 7 - 9 mo. | F/M | 6J/6J Cre$^-$ *Rars1*$^{loxP/-}$<br>6J/6J Cre$^+$ *Rars1*$^{-/-}$<br>6J/6J Cre$^+$ *Rars1*$^{loxP/-}$ |
| IP injection, 90 mg/kg, 6 hr<br>Blood and urine collection for clinical chemistry | 3 mo. | F/M | 6J/6J Cre$^-$ *Rars1*$^{loxP/loxP}$ |
| IP injection, 90 mg/kg, 2-4 hr<br>Urine collection for detection of metabolites | 4 mo. | F | 6J/6J Cre$^-$ *Rars1*$^{loxP/loxP}$ |
| IP injection, 2 × 45 mg/kg over 8 hr<br>Brain tissue for RNA-seq and proteomics | 4 mo. | F | 6J/6N Cre$^-$ *Rars1*$^{loxP/-}$ |
| IP injection, 2 × 45 mg/kg over 8 hr<br>Comparison of protein synthesis in Wt and dLZ brains by proteomics | 2.5 mo. | M | 6J/6J Cre$^+$ *Rars1*$^{-/-}$ (TePhe)<br>6J/6J Cre$^+$ *Rars1*$^{loxP/-}$ (Veh only)<br>6J/6J Cre$^+$ *Rars1*$^{loxP/loxP}$ (TePhe) |

All animals are from the Rars1$^{tm1Cuiut}$ line. Cre$^+$ refers to animals having at least one copy of *Emx1*$^{tm1(cre)Krj}$, while Cre$^-$ animals have no copies. 6J and 6N refer to C56BL/6 substrains differing at the *n-Tr20* locus (neuronal tRNA-Arg-TCT-4-1, C50T in 6J). All 6J/6N individuals are the offspring of an inbred C56BL/6J parent from our colony and a JAX C56BL/6N parent, each homozygous for their respective alleles. Genotype is not expected to impact the outcome of TePhe experiments.

**Section 3. List of Stains and Antibodies**

| Target/Antigen | Supplier and Catalogue # | Description | Clone/Lot information | Dilution |
|---|---|---|---|---|
| Nissl bodies | Invitrogen N21482 | NeuroTrace 530/615 Red Fluorescent Nissl Stain | - | 1:200 |
| NeuN | ProteinTech 26975-1-AP | Rabbit polyclonal antibody | Lot 00105221 | 1:200 |
| MAP2 | Cell Signalling 4542S | Rabbit polyclonal antibody | Lot 4 | 1:1000 |
| Olig2 | EMD Millipore AB9610 | Rabbit polyclonal antibody | Lot 3857662; Lot 4267458 | 1:150 |
| cFos | Synaptic Systems 226 008 | Rabbit monoclonal antibody | Clone Rb108B5 Lot 1-45 | 1:550 |
| eIF2α | Cell Signalling 9722S | Rabbit polyclonal antibody | Lot 15 | 1:250 |
| α-tubulin | Cell Signalling 2125S | Rabbit monoclonal antibody | Lot 15 | 1:60 |
| Sox2 | Protein Tech 11064-1-AP | Rabbit polyclonal antibody | Lot 00098845 | 1:150 |
| Doublecortin (Dcx) | Cell Signalling 4604S | Rabbit polyclonal antibody | Lot 7 | 1:600 |
| Rabbit isotype control | BioLegend 910801 | Rabbit polyclonal antibody | Lot B429063 | As needed to match 1° antibody used |
| Dylight 650-conjugated goat-anti-rabbit IgG | CedarLane CLCC46018 | Goat polyclonal antibody | 98-90-072023 | 1:250 |

**Section 4. Procedures for equilibrating/washing streptavidin beads**

Magnetic streptavidin beads (Genscript L00936) were used for enrichment of OSTAC-biotinylated proteins from HEK293T lysate. Bulk beads were transferred to a microcentrifuge tube, diluted in PBS (up to 100 µL beads per 1.5 mL), and mixed well by repeated inversion. Beads were manually immobilized to one side of the tube using a bar magnet to allow removal of the supernatant. The PBS wash was repeated once. Beads were then suspended in PBS + 1% SDS (1.5 mL), vortexed vigorously, and further mixed on an over-the-end rotator for 5 min. The supernatant was removed and the PBS + 1% SDS wash repeated 3 more times. On the $4^{th}$ wash, suspended beads were aliquoted into 2 mL tubes (approximately 12 µL settled bead volume per sample of 480 µg protein input.) Beads at this stage were considered equilibrated in PBS + 1% SDS and ready for binding. OSTAC-treated HEK293T lysates were diluted to 0.25 mg/mL in PBS + 1% SDS and applied to washed beads. Samples were incubated at room temp. for 1 hr 15 min with over-the-end rotation. Unbound proteins were removed by washing with 4 × PBS + 1% SDS (1.5 mL), 2 × PBS (1.5 mL), and 2 × 100 mM ammonium bicarbonate (1.5 mL). Each wash step consisted of resuspension in desired wash buffer, mixing by repeated inversion of the tube, and agitation on a tilting and rotating platform for 5 min. Washed beads were suspended in fresh 100 mM ammonium bicarbonate (200 µL), to which was added sequencing grade trypsin (3.5 µg). Slurries were incubated at 37 °C, 1200 rpm for 18 hr before proceeding to peptide clean-up as described in the Methods section.

Non-magnetic streptavidin agarose resin (Genscript L00353) was used for the capture of OSTAC-biotinylated proteins from N2a and mouse brain lysates. Bulk beads were transferred to a microcentrifuge tube, diluted in PBS (up to 100 µL beads per 1.5 mL), and mixed well by repeated inversion. Beads were pelleted at 2000 × g for up to 5 min (beads pelleted relatively slowly in non-detergent-containing buffer), transferred to a tube rack, and allowed to settle for a few minutes. Supernatant was carefully removed (~50 µL residual volume). The PBS wash was repeated once. Beads were then equilibrated with 4 washes of PBS + 1% SDS (1.5 mL). Each wash consisted of vigorous mixing (4 cycles of alternating vortex (5 sec) and manual inversion of the tube (8x), adapted from Kottala et al.[2]), and pelleting at 2000 × g for 1 min. During the $4^{th}$ wash, suspended beads were aliquoted into 2 mL tubes (approximately 8 µL settled beads per sample). OSTAC-treated lysates were diluted to 0.25 mg/mL (450 µg input for N2a lysates, 290 µg input for brain lysates), applied to equilibrated beads and incubated at room temp. for 1.5 hr with over-the-end rotation. To remove unbound proteins, beads were first washed with 4 × PBS + 1% SDS (1.8 mL). Each wash consisted of resuspension in wash buffer, 4 cycles of alternating vortex (5 sec) and manual inversion of the tube (8x), and further mixing on an over-the-end rotator for 5 min. Beads were pelleted at 2000 × g for 1 min and allowed to settle for 1 min. Beads were then washed with 2 × PBS (1 mL) and 2 × 100 mM ammonium bicarbonate (1.5 mL); after each resuspension, beads were mixed briefly by vortex. Beads pelleted less readily as SDS concentration decreased; later washes required centrifugation for up to 5 min. Washed beads were resuspended in fresh 100 mM ammonium bicarbonate (120 µL) and digested with addition of trypsin (3.0 µg for N2a lysates, 2.4 µg for brain lysates) at 37 °C, 1200 rpm for 16 hr before proceeding to peptide clean-up.
